# Engineering Ca^2+^-dependent DNA polymerase activity

**DOI:** 10.1101/2023.05.11.540423

**Authors:** Bradley W. Biggs, Alexandra M. de Paz, Namita J. Bhan, Thaddeus R. Cybulski, George M. Church, Keith E. J. Tyo

## Abstract

Advancements in synthetic biology have provided new opportunities in biosensing with applications ranging from genetic programming to diagnostics. Next generation biosensors aim to expand the number of accessible environments for measurement, increase the number of measurable phenomena, and improve the quality of the measurement. To this end, an emerging area in the field has been the integration of DNA as an information storage medium within biosensor outputs, leveraging nucleic acids to record biosensor state over time. However, slow signal transduction steps, due to the timescales of transcription and translation, bottleneck many sensing-DNA recording approaches. DNA polymerases (DNAPs) have been proposed as a solution to the signal transduction problem by operating as both the sensor and responder, but there is presently a lack of DNAPs with functional sensitivity to many desirable target ligands. Here, we engineer components of the Pol δ replicative polymerase complex of *Saccharomyces cerevisiae* to sense and respond to Ca^2+^, a metal cofactor relevant to numerous biological phenomena. Through domain insertion and binding site grafting to Pol δ subunits, we demonstrate functional allosteric sensitivity to Ca^2+^. Together, this work provides an important foundation for future efforts in developing DNAP-based biosensors.

## Introduction

Progress within synthetic biology has led to a remarkable expansion in biosensing technology, both with respect to the range of detectable analytes (inputs) and functional responses (outputs) that are possible. Today, not only can protein^1–3^, nucleic acids^4–6^, metabolites^7–11^, cofactors^12–15^, ions^16–19^, and light^20–22^ be sensed, but outputs ranging from colorimetric changes (for rapid and affordable diagnostics)^23–25^ to dynamically controlled gene expression (for genetic programs)^26–30^ have been realized. Many of these approaches have been built upon only a handful of protein classes with transcription factors^31–34^, proteases^35–37^, T7 RNA polymerase^38–40^, GPCRs^41–43^, and CRISPR-Cas systems^44–46^ acting as the foundation for numerous technologies^47^. Inspired by technical advances and reduced costs in genomics, an emerging area within biosensing is recording the biosensor output into a DNA sequence^48^. For phenomena that occur in visually occluded locations, this approach holds the potential to alleviate issues posed by current standards for measurement such as microscopy and could provide informationally dense, minimally disruptive, and possibly continuous sensing-recording.

Several modalities have been explored within the biosensing-DNA recording paradigm, harnessing recombinases^49^, base editors^50^, nucleases^51–53^, and polymerases^54^. Each of these systems requires induction of expression to record an event. Therefore, these strategies cannot sense and record biological phenomena that occur on timescales shorter than the combined time of transcription, potentially translation, and the subsequent DNA modification step. One proposed solution is to mimic how biological systems respond quickly to environmental changes, such as nutritional shifts, and to instead utilize post-translational regulation (i.e. allostery, post-translational modification)^55–57^ in combination with an enzyme that can directly act on DNA. Allosteric regulation or altering the activity of a DNA modifying enzyme, such as the error rate of a DNA polymerase (DNAP) where the frequency of errors in the copied DNA correlates with the signal, could provide more direct signal transduction by filling both the sensor and recorder roles^58^ and thus provide a foundation for a new class of biosensors.

While there are several possible implementations within the allosteric DNAP biosensor concept, the simplest from a technical execution standpoint is a two-polymerase system, where one DNAP is functionally ligand-sensitive while the other is insensitive (Figure 1a, green and gray enzymes respectively). Within this framework, as described below, it is also important that the two polymerases have disparate error rates. The fundamental first step for this approach, which is the focus and scope of this work, is the identification or engineering of a DNAP to be functionally sensitive to the molecule of interest that can be paired with a reference (insensitive) polymerase. If both polymerases are allowed to copy a known DNA template, and if the chosen pair of DNAPs possess different replication characteristics (e.g., error rates), then the presence or absence of the ligand can be distinguished by the resultant DNA. In the absence of the ligand, the output DNA will represent the combined function of both DNAPs (green and gray bars in Figure 1a). In the presence of the ligand, the output DNA will primarily represent the replication characteristics of the insensitive DNAP (Figure 1a, gray bars), as the engineered DNAP (green) will have reduced activity. The resulting DNA can then be read as a molecular “ticker-tape,” reporting on the presence or absence of the ligand within the given biological context over time^58^. We recently accomplished a two-polymerase system *in vitro* using an engineered deoxynucleotidyl transferase (TdT) to record calcium signal^59^. However, TdT creates single stranded DNA as an output, which could potentially be targeted for degradation in certain biological contexts. Therefore, in this work we sought to engineer a template-dependent alternative that would generate double stranded DNA products.

**Figure 1.**
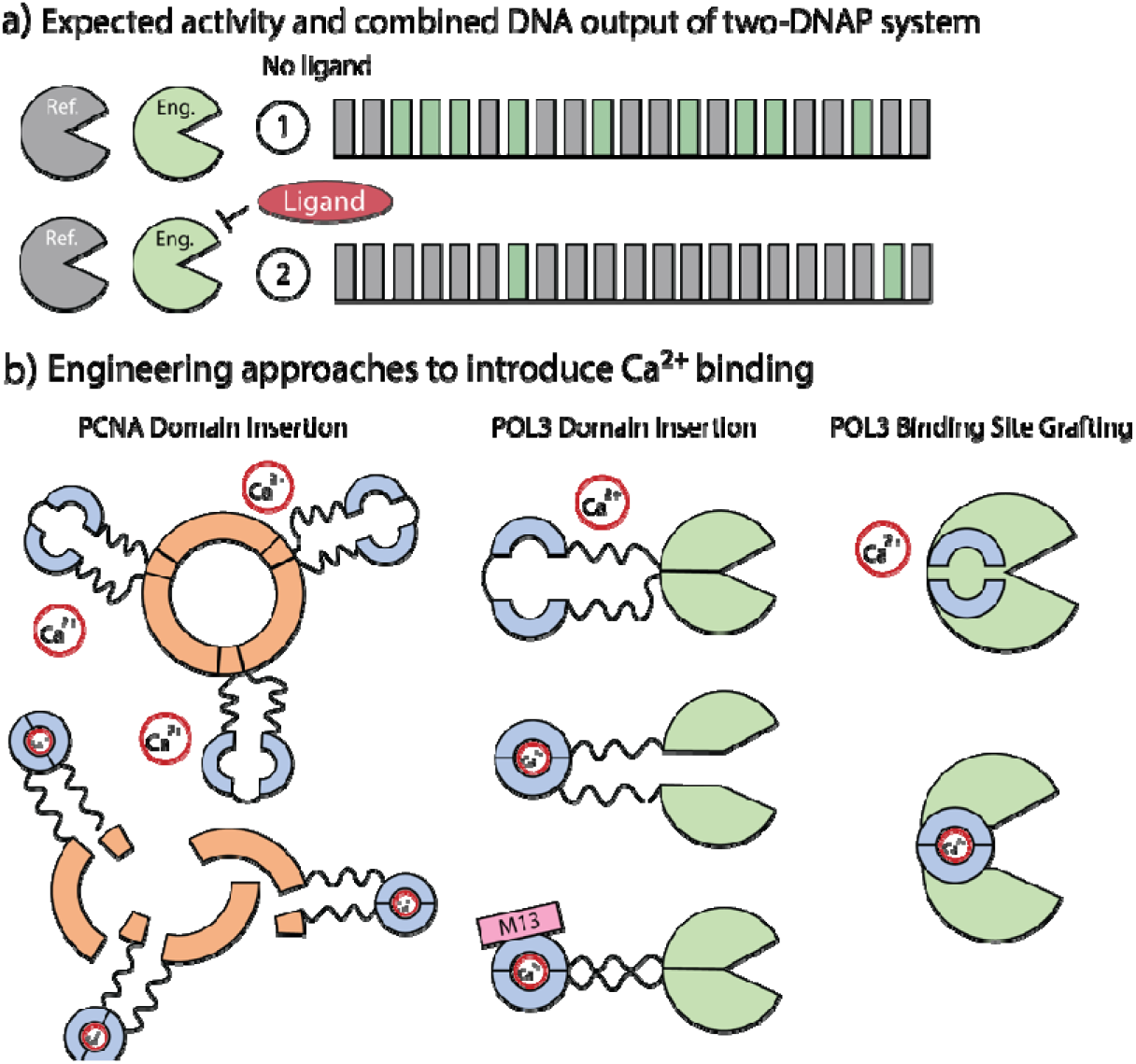
Outline of DNAP biosensor and recording. **a)** Depicts expected function of a two-DNAP system, where reference DNAP (Ref.) in gray is unaffected by the presence of a given ligand and maintains typical activity, while th activity of an engineered (Eng.) DNAP in light green is ligand sensitive. DNA output is shown for the case where eac polymerase has distinct replication characteristics (e.g., replication error rate). In the absence of the ligand, the resultant DNA is a mixed representation of each DNAP’s activity, but in the presence of the ligand, the output DNA predominantly represents the reference DNAP (gray). **b)** Three engineering approaches to create DNAP Ca^2+^ sensitivity. On the left is CaM domain (light blue) insertion into PCNA (orange), where the trimeric ring and typical protein function is disrupted in the presence of Ca^2+^. In the middle is CaM domain (light blue) insertion into Pol3 (light green), the catalytic subunit of the Pol δ complex, where Pol3’s activity is disrupted in the presence of Ca^2+^ and potentially restored in the presence of M13, a peptide which binds Ca^2+^-loaded CaM. On the right is binding site grafting, where the consensus residues of CaM’s EF-hand Ca^2+^-binding site (light blue) are grafted onto the exonuclease domain of Pol3 (light green) to deliver Ca^2+^ sensitivity and reduced activity in the presence of Ca^2+^.

Here, as with our TdT engineering, we focus on sensing and responding to Ca^2+^, a metal ion that is biologically ubiquitous and important in several physiological contexts^60^, including some that are difficult to measure by traditional means^61^. Importantly, no template-dependent DNAP is known to be modulated by Ca^2+^ concentration in a manner conducive to enabling recording, though some have been tested for this capacity^62^. The trait, accordingly, must be engineered. To accomplish this, we have employed two protein engineering approaches, domain insertion^17,45,63–65^ and binding site grafting^66^. Domain insertion, as the name suggests, involves the insertion of an existing functional domain possessing a trait of interest, such as an analyte-binding motif, into the target protein sequence, thereby imbuing new sensitivity to the target protein while potentially maintaining its original function. Binding site grafting, instead, involves the sequential mutation of a section of the target protein, without the introduction of additional amino acids to the total sequence length, until it resembles a known functional domain and can perform the expected task (i.e. analyte binding)^67^.

For both approaches, the model calcium sensor calmodulin (CaM)^68^ was utilized, with the entire protein being used for domain insertion and CaM’s EF-hand Ca^2+^-binding site subdomain used as a template for binding site grafting. Calmodulin has been a popular choice for analogous applications because of its pronounced conformational change upon Ca^2+^ binding^69^, which is anticipated to disrupt typical enzyme function^70^. The DNAP chosen for engineering in this work is the Pol δ complex (Pol3, Pol31, Pol32, Proliferating Cell Nuclear Antigen [PCNA]) of *Saccharomyces cerevisiae*, as it is well-studied, possesses a crystal structure for the catalytic domain Pol3 (PDB ID: 3IAY), is active at physiologically relevant temperatures, has natively high fidelity and processivity, and is highly replicative^71^. Using these approaches, we demonstrate Ca^2+^ functional sensitivity within the Pol δ complex through (1) domain insertion into PCNA, (2) domain insertion into the catalytic subunit Pol3, and (3) binding site grafting onto Pol3 (Figure 1b). While additional work is required to optimize polymerase interactions to implement two polymerase recording, this work represents an important and fundamental step towards engineering DNAP-based recording systems.

## Results and Discussion

### PCNA domain insertion

DNAP structure is highly conserved^72^, suggesting that structural disruption through domain insertion or other engineering approaches is likely to inhibit typical enzyme function. Accordingly, we began with a more cautious approach to generate DNAP Ca^2+^ sensitivity by first targeting a DNAP accessory protein for engineering. Proliferating cell nuclear antigen (PCNA) is a DNAP-associated protein that assists in polymerase processivity by forming a trimeric “clamp” around the DNA being replicated while binding a partner DNAP (Figure 2a)^73^. When bound and interacting with a DNAP, PCNA is known to improve processivity while negatively impacting fidelity between 2 and 7-fold^74^. Therefore, we hypothesized that by engineering Pol δ-associated yeast PCNA to be Ca^2+^ sensitive, we might indirectly achieve DNAP functional sensitivity.

**Figure 2.**
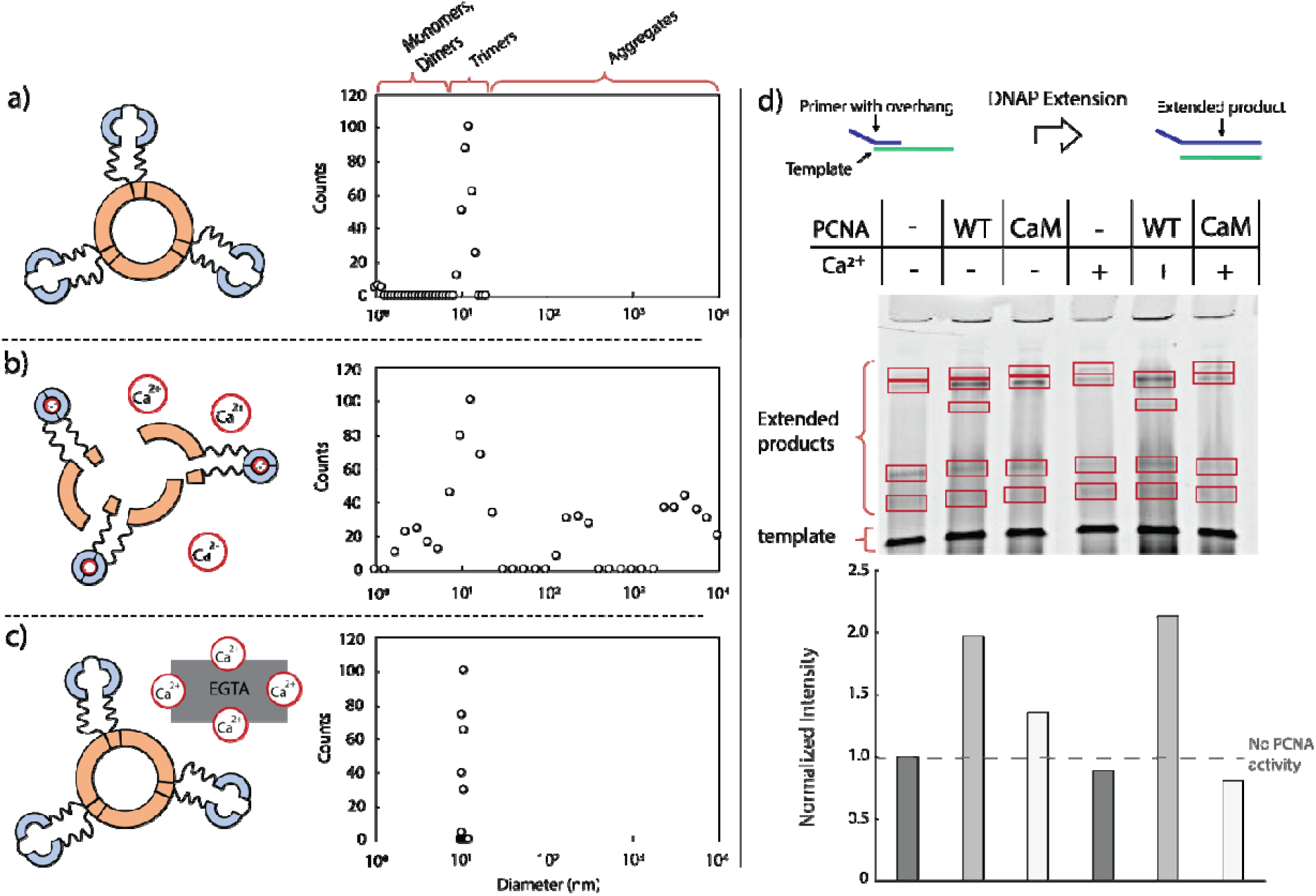
PCNA domain insertion. Panels **a-c** show the reversible impact of Ca^2+^ addition to F5-PCNA-CaM (where PCNA is depicted in orange and CaM domain insertion is in light blue). In **a)**, where Ca^2+^ has not yet been added, PCNA is primarily seen in its trimeric form as indicated by particle count at the predicted diameter using dynamic light scattering (DLS). In **b)**, upon addition of Ca^2+^, the dominant trimeric form is disrupted and both lower diameter species (monomers) and higher diameter species (aggregates) are observed. In **c)**, upon addition of EGTA to chelate the Ca^2+^, the trimeric form is restored and a tight distribution around the expected trimeric diameter is observed. In **d)**, summed intensities gathered from gel analysis of extension assays show a decrease in the amount of DNAP extension upon addition of Ca^2+^ for the “CaM” condition (where “CaM” refers to F5-PCNA-CaM), with wild-type PCNA being unaffected by Ca^2+^ addition.

We began by identifying sites within PCNA that could be tolerant to CaM domain insertion. Because of a lack of high-throughput methods for easily screening mesophilic DNAP activity, we sought to reduce the cloning load by first applying computational screening methods. Specifically, we utilized an approach that has previously been successful for us^59^ and others^75,76^ based on the program SCHEMA^77^, which considers structural coordinates of homologs to identify pairs of interacting amino acid residues to minimize the number of disrupted residue-to-residue contacts upon recombination^77–79^. Previously, a residue proximity between TEM-1 β lactamase split sites and SCHEMA crossover sites had been observed, and this information was used to generate functional TEM-1 β lactamase chimeras^75,76^. Accordingly, we applied SCHEMA to predict likely successful PCNA-CaM chimeras as well, with the SCHEMA-predicted crossover sites functioning as general protein split points that would be tolerant to domain insertion. Using this method, an ideal location was identified (K107). Table S1 shows the multi-sequence alignment used for PCNA. K107 is part of a loop that connects the β sheet at the C-terminus of each monomer. These β-sheets form hydrogen bonds between monomers, resulting in the final homotrimeric version of the protein. We speculated that inserting CaM in this loop would cause the trimerization of PCNA to become dependent on the conformation of CaM. This would in turn make the PCNA association with the Pol δ complex dependent on the conformation of CaM and thus Ca^2+^. We tested both inserting the CaM domain after the K107 residue (PCNA-CaM fusions 1, 3, 5) and replacing the K107 residue (PCNA-CaM fusions 2, 4, 6) with the inserted CaM domain. For each approach, we tested three linkers: a short flexible glycine-serine “GS” linker (F1-PCNA-CaM, F2-PCNA-CaM), a longer flexible “GSGGG” linker (F3-PCNA-CaM, F4-PCNA-CaM), and a short rigid three-amino acid linker composed of aspartate-lysine-serine “DKS” (F5-PCNA-CaM, F6-PCNA-CaM).

To quickly examine the ability of PCNA-CaM fusions to bind Ca^2+^, we utilized an electrophoretic shift assay^80–83^. Previous work had shown that when CaM binds Ca^2+^, its apparent molecular weight reduces by approximately 4.5 kDa^84^, likely due to exposure of a hydrophobic surface resulting from the conformation change. For all fusions generated, the addition of Ca^2+^ lowered the apparent molecular weight by ∼3 kDa compared to the reference state of only including the chelator EGTA (Figure S1). Next, we examined whether the PCNA-CaM fusion maintained native β-sheet interactions to form wild-type like trimers, as trimeric formation is necessary for downstream functionality. To capture this structure, we used ethylene glycol bis(succinimidyl succinate) (EGS) crosslinking to convert the β-sheet hydrogen bonds into covalent bonds. Using one of the fusions as an example (F5-PCNA-CaM, CaM insertion after K107 using the longer flexible “GSGGG” linker), we analyzed the crosslinking with a Western blot stain of an SDS-PAGE gel. Gel analysis showed that indeed the fusion was able to form trimers similarly to wild-type PCNA (Figure S2).

Following the establishment of Ca^2+^ binding and confirming the retained ability to form trimers, we next moved to testing the fusion’s dynamic response to Ca^2+^ to determine if the trimeric form is disrupted by Ca^2+^ binding and if this response is reversible. For this, we used a dynamic light scattering (DLS) assay, which measures the distribution of particle size for protein solutions. Using predicted sizes of particles for the monomeric and trimeric forms of PCNA, as determined by PyMol, we used the DLS assay to determine how much of the PCNA was in each form. First, we tested wild-type PCNA and found that the mean particle diameter distribution was around the expected 9 nm for the trimer (Figure S3). Upon inclusion of 4 mM CaCl_2_ to the solution, wild-type PCNA retained its mean diameter distribution (Figure S4), suggesting that wild-type PCNA is not affected by the addition of Ca^2+^. Next, F5-PCNA-CaM was examined for its Ca^2+^-responsiveness, chosen among the fusions for its small rigid linker and intact K107 residue. In a state with only EGTA, which chelates Ca^2+^, F5-PCNA-CaM has a mean diameter distribution of approximately 12 nm (Figure 2a). This corresponds to the modest expected increase compared to wild-type (9-15 nm expected), due to the inserted CaM domain. Upon addition of Ca^2+^, the distribution changed. Instead of having a tight mean around the expected trimeric size, polydispersion was observed, as both lower and higher molecular weight species appeared (Figure 2b). The lower molecular weight species corresponded with the expected size of monomeric protein, and the much higher molecular weight species likely indicated protein aggregates. This suggests that Ca^2+^ binding disrupted the typical trimeric structure leading to monomers and aggregates of PCNA-CaM. Upon addition of EGTA to chelate the free Ca^2+^, DLS showed that the mean particle diameter returned to approximately 10 nm (Figure 2c), indicating that trimeric formation had been restored and that Ca^2+^-based disruption is reversible. Time course data for the DLS experiments can be found in the supplement (Figures S5-S7).

Lastly, to determine if the change in PCNA oligomeric formation impacts DNAP function, a fluorescent primer extension assay was carried out with the Pol δ complex. This assay uses a DNA extension construct consisting of an 89-base, 5’ TAMRA-labeled template with an annealed 60-base 5’ FAM-labeled primer, creating a 3’ recessed strand that can be extended by replication using the PCNA and Pol δ complex (Figure S8). The TAMRA and FAM labeling allow distinction of the different DNA strands by fluorescent imaging. Successful extension generates a DNA species with higher molecular weight (124-base product) than either the template or primer. The extension product can be resolved via polyacrylamide gel electrophoresis (PAGE) under denaturing conditions and subsequently imaged based on FAM fluorescence. Using this assay, we were able to show that adding wild-type PCNA to the Pol δ complex does indeed lead to increased DNA synthesis activity and that wild-type PCNA is unaffected by the inclusion of Ca^2+^ (Figure 2d, columns 2 and 5). However, F5-PCNA-CaM showed reduced activity with the inclusion of Ca^2+^ (Figure 2d, columns 3 and 6). Gels for replicate experiments can be found in the supplement (Figures S9 and S10). Taken together, our data demonstrates that PCNA is reversibly affected by the presence of Ca^2+^, and that its natural function with the Pol δ complex is impacted by this sensitivity.

### Pol3 domain insertion

We next adopted a more direct approach for modulating DNAP activity by engineering the catalytic subunit of Pol δ, Pol3, to be Ca^2+^ sensitive. As before, we utilized SCHEMA to predict permissive sites for inserting CaM into Pol3. We identified 19 frequently occurring crossover residues in wild-type Pol3 as potential CaM insertion sites. These SCHEMA-predicted residues spanned all major subdomains of Pol3 (including the palm, thumb, fingers, exonuclease, and N-terminal domains) and were characterized by a range of average B-factors (∼20 to ∼37), which were calculated using PyMOL (PDB ID: 3IAY) to obtain a better sense of the flexibility at each particular position (Table S2). The B-factor represents the fluctuation of atoms about their average position, with larger values signifying higher flexibility at a particular region^85,86^. For each of the 19 identified sites, a Pol3-CaM fusion was constructed with flexible linkers (GSGGG)^87^ (Figure S11), using a variant of Pol3 with N- and C-terminal deletions matching those utilized for the crystal structure^71^ that had been codon optimized for *E. coli* expression. As with the PCNA-CaM approach, the expectation for the Pol3-CaM fusions was that the addition of Ca^2+^ would negatively impact Pol3’s catalytic activity via CaM’s conformation shift upon binding Ca^2+^ (Figure 3a-b).

**Figure 3.**
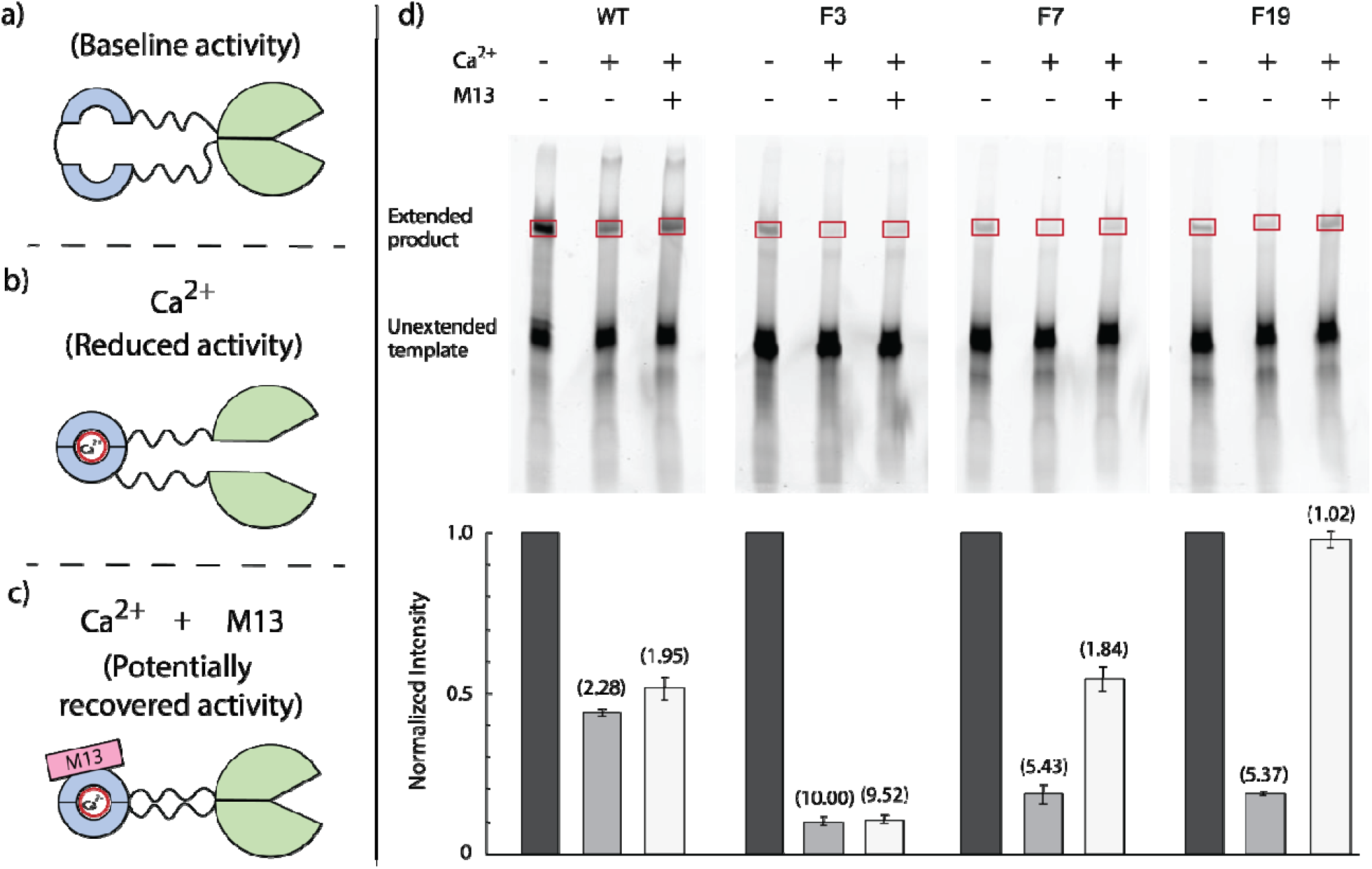
Measuring activity modulation potential of select Pol3-CaM fusions. Panels **a-c** depict the expected impact of **a)** Mg^2+^, **b)** Ca^2+^, and **c)** Ca^2+^ + M13 peptide conditions on Pol3-CaM fusion activity, where Ca^2+^- and M13-triggered conformational changes in CaM (light blue) can propagate to Pol3 (light green) and potentially modulate DNAP activity. Panel **d)** shows activity testing of wild-type Pol3 (WT), F3-Pol3-CaM (F3), F7-Pol3-CaM (F7), and F19-Pol3-CaM (F19) under Mg^2+^, Ca^2+^, and M13 peptide conditions. FAM-labeled extension products (124 bases, designated by red box) were resolved on 10% PAGE under denaturing conditions and FAM fluorescence was imaged (λ^ex^ = 488 nm, λ^em^ = 520 nm). Lanes were skipped to prevent cross-contamination of samples during gel loading. Mg^2+^, Ca^2+^, and M13 peptide conditions for a given Pol3 variant were run on the same gel to enable quantitative comparison. Fold change in activity of wild-type Pol3 (WT), F3-Pol3-CaM (F3), F7-Pol3-CaM (F7), and F19-Pol3-CaM (F19) under Ca^2+^ and M13 peptide conditions (compared to the baseline Mg^2+^ condition) was calculated (n=4, SEM plotted). Gel image analysis was performed to quantify intensities of extension products. Two independent extension reaction experiments were performed per condition and two technical replicates were imaged per experiment. Product intensities were background subtracted and used to calculate relative changes in activity for a given DNAP. Fold change calculations were confined to samples within the same gel.

All 19 variants were expressed *in vitro* using NEB PURExpress and were tested for activity using the previously implemented fluorescent primer extension assay (Figures S8, S12-S23). Standard conditions (where magnesium [Mg^2+^] was the only divalent cation present) were initially used to determine the impact of the CaM insertion on DNAP activity (Figure S12). Qualitative gel analysis revealed varying activity levels for the 19 Pol3-CaM fusions, with most variants demonstrating no catalytic activity. A subset of Pol3-CaM fusions, F2-Pol3-CaM (C-terminal), F3-Pol3-CaM (exonuclease), F7-Pol3-CaM (palm), and F19-Pol3-CaM (palm), revealed a stronger baseline activity signal, with F2-Pol3-CaM demonstrating near wild-type levels of activity. Tolerance of CaM insertion into the exonuclease domain of Pol3, as demonstrated by F3-Pol3-CaM, can potentially be explained by the fact that exonuclease activity is not essential to polymerase function^88^. Unexpectedly, functional activity of F7-Pol3-CaM and F19-Pol3-CaM, both palm domain fusions, indicate that insertions in this region, which contains the polymerase active site^71^, are possible without interrupting DNAP activity.

The next step was to test F2-Pol3-CaM, F3-Pol3-CaM, F7-Pol3-CaM, and F19-Pol3-CaM under Ca^2+^ conditions to determine ON/OFF switching potential (Figure 3d, Figures S14-23 for extension assay gels of all Pol3-CaM fusions). Calmodulin undergoes two pronounced conformational shifts that could potentially be exploited for activity modulation: 1) from “closed” to “open” upon binding Ca^2+^ (resulting in a ∼22 angstrom separation of N- and C-termini), and 2) from “open” to “bound” when a protein target binds Ca^2+^-loaded CaM^68,89,90^. To capture both conformational shifts in our testing, we tested a Ca^2+^ condition that included M13, a peptide derived from myosin light chain kinase that binds to CaM in the presence of Ca^2+91,92^ (Figure 3c). Activity levels of F2-Pol3-CaM, F3-Pol3-CaM, F7-Pol3-CaM, and F19-Pol3-CaM were quantified using image analysis and normalized to the base Mg^2+^ condition (no Ca^2+^) to calculate fold-change in activity for each variant (Figure 3d). Although wild-type Pol3 activity was slightly decreased in the presence of Ca^2+^, the change was only modest compared to the impact of Ca^2+^ on F3-Pol3-CaM, F7-Pol3-CaM, and F19-Pol3-CaM, which showed a 10-fold, 5-fold, and 5-fold reduction in activity, respectively, compared with a 2-fold activity decrease for wild-type.

Interestingly, the Ca^2+^ impact for the exonuclease domain fusion (F3-Pol3-CaM) was nearly two-fold stronger than for the two palm domain fusions (F7-Pol3-CaM and F19-Pol3-CaM). This could potentially be explained by insertion site secondary structure, as inserting CaM in the middle of an α-helix (F3-Pol3-CaM, exonuclease domain) is more likely to propagate Ca^2+^-triggered disruption than an insertion in an unstructured loop (F7-Pol3-CaM and F19-Pol3-CaM, palm domain)^93,94^. Perhaps unsurprisingly, F2-Pol3-CaM, a C-terminal fusion which posed the least potential disruption to Pol3^93^, showed a pattern similar to wild-type, with a slight decrease in signal in the presence of both Ca^2+^ and M13 (Figures S14 and S24).

Interestingly, the inclusion of M13 had different effects on fusion activity. While the addition of M13 had virtually no impact on F3-Pol3-CaM activity in the presence of Ca^2+^, F7-Pol3-CaM recovered around half of its pre-Ca^2+^ activity in the presence of M13 whereas F19-Pol3-CaM recovered nearly all of its pre-Ca^2+^ activity (Figure 3d). Given that F7-Pol3-CaM and F19-Pol3-CaM are only two residues apart (Y587 and G589, respectively), it’s not surprising that both variants were modulated similarly by Ca^2+^ and M13. Furthermore, while CaM binding to M13 triggers a return to a near-closed CaM conformation, it does not necessarily follow that the structural motifs split by CaM will also return to their intact forms. Specifically, the disrupted α-helix in F3-Pol3-CaM, along with the potentially disrupted nearby secondary structures, are unlikely to be restored to their original forms upon M13 binding, whereas the unstructured loop region in F7-Pol3-CaM and F19-Pol3-CaM is likely more forgiving of conformational changes^93,94^. Overall, we demonstrate Ca^2+^-driven activity modulation in Pol3 by inserting CaM in both the exonuclease and palm domains. Together, these results strongly implicate CaM as a key functional domain in mediating allosteric Ca^2+^-based changes to DNAP activity.

### Pol3 binding site grafting

The incorporation of a full CaM domain in engineered DNAPs may be variably affected by the presence of M13-like peptides in an *in vivo* application. Similarly, bearing a full CaM domain on the polymerase may interact with such peptides and impact overall cellular function. Our results themselves indicate that M13 peptide can variably impact the allosteric function of Pol3-CaM fusions (Figure 3d). Importantly, different physiological contexts where it is desirable to measure Ca^2+^ could possess varying CaM-interacting peptides such as M13. Expressly, a study showed GCaMPs with full CaM impacted calcium channel function^95^. This motivated a more finessed engineering approach, and we next decided to introduce Ca^2+^ sensitivity to Pol3 using binding site grafting of an incomplete CaM domain^96^, instead of full domain insertion, to minimize potential unwanted interactions (Figure 4a). In this approach, amino acid residues are mutated to create a new Ca^2+^-binding site out of the existing protein structure. Binding site grafting has been used to successfully introduce Ca^2+^ binding into natively non-Ca^2+^ binding proteins^97^ and even to introduce Ca^2+^-dependent functionality^98^. This approach has predominantly used the highly conserved, twelve residue EF-hand motif from CaM as a template binding site^99^. Successful binding site grafting is primarily dependent upon choosing a location in the protein with ideal Ca^2+^-binding characteristics. Previous work has identified important criteria to include 1) being solvent exposed to allow for water to coordinate negatively charged residues when Ca^2+^ is not bound, 2) being flexible to allow for conformation changes, and 3) for the site to be in a region that is unlikely to disrupt typical enzyme function (e.g. not the active site)^100^. Examining Pol3 for a suitable site for grafting, a region within the exonuclease domain fit the desired characteristics well (K416 to G427) (Figure 4b). In addition to being solvent exposed and flexible, it originates from an α-helix, which parallels the EF-hand motif of CaM (Figure S25).

**Figure 4.**
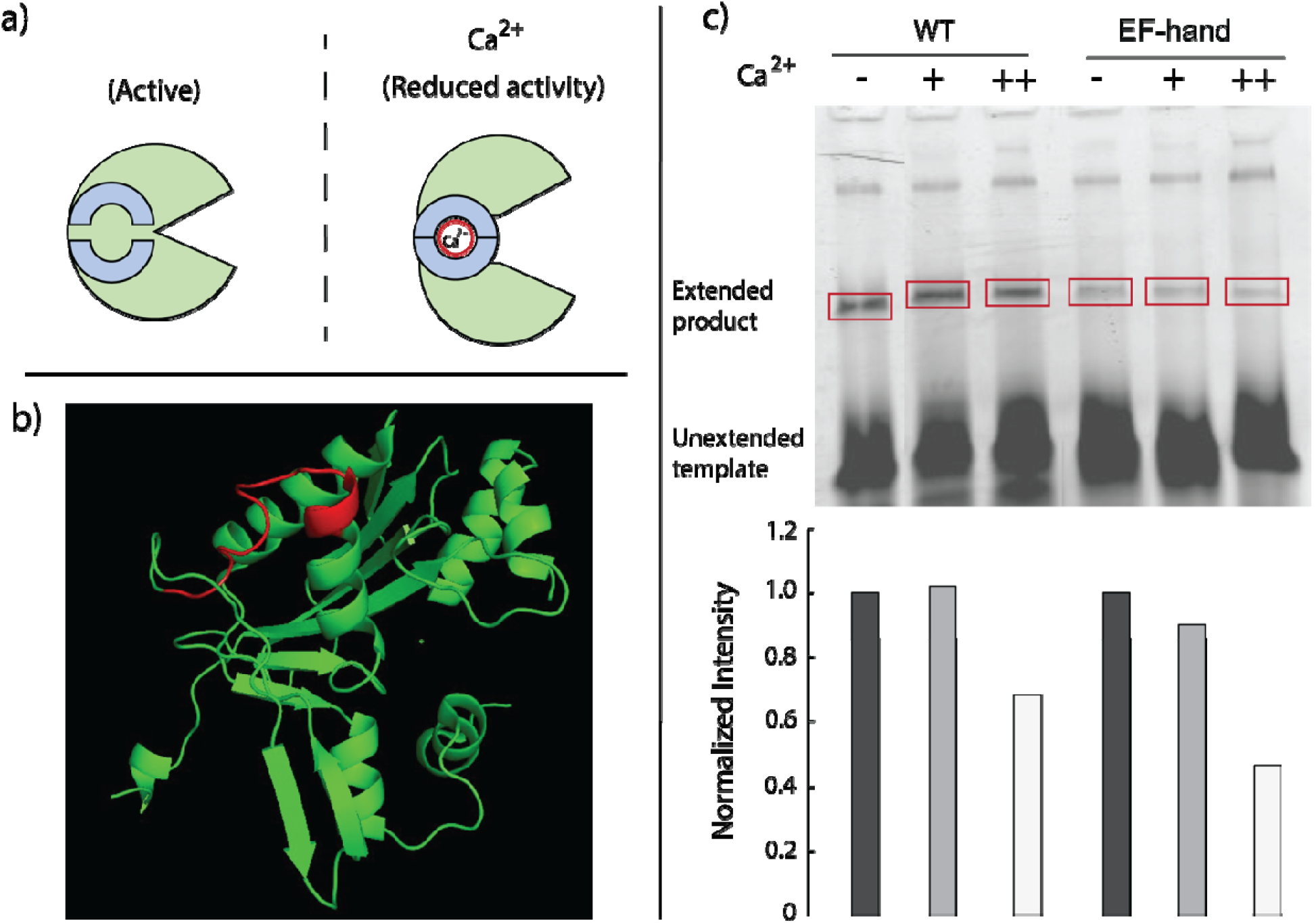
Pol3 binding site grafting approach. In **a)**, schematic demonstrating the expected function where the engineered Pol3 variant (light green) has reduced activity upon Ca^2+^ binding to the introduced EF-hand site (light blue). In **b)**, highlighted in red is the site within the exonuclease domain of Pol3 for binding site grafting because of its ideal characteristics. Image was created with PyMOL using the PDB 3IAY. In **c)**, fluorescent extension assay gel shows the greater impact of calcium upon the engineered EF-hand bearing Pol3 variant.

Accordingly, stepwise mutations were introduced at this location to create a Ca^2+^-binding site. To initially test if stepwise mutations disrupted Pol3 catalytic activity, variants were screened using a complementation assay in *Saccharomyces cerevisiae* (Figure S26). This assay utilizes a yeast strain where wild-type Pol3 has been removed from the chromosome and is instead contained on a plasmid that bears the *URA3* marker. By transforming yeast with a plasmid bearing a Pol3 variant and *LEU2* marker, using yeast homologous recombination to assemble the plasmid *in vivo*, and plating on medium that lacks leucine but contains uracil and 5-fluoorotic acid (5-FOA), the plasmids can be exchanged. If yeast colonies form, it indicates that the variant can viably maintain yeast duplication. If no colonies form, then the Pol3 variant is likely either too slow or error prone to maintain replication. Using this assay, we found that the yeast strain formed colonies with six of the seven key mutations made, but upon adding the 7^th^ mutation (K416E), which forms the anchor point for the EF-hand and completes the primary consensus sequence, colonies no longer form (Table S3). By moving the anchor point one residue (A415E), colonies do form. In addition, the single mutation K416E by itself allowed for cell growth. Interestingly, removing part of the consensus EF-hand Ca^2+^-coordinating residues (D425A or D427A), while maintaining the rest of the motif, does not recover growth, perhaps due to an ability to still bind Ca^2+^ albeit to a lesser extent. This data suggests that the EF-hand variant of Pol3 may be responding to intracellular Ca^2+^ concentrations, but is not definitive. Unfortunately, because the cells could not grow while depending upon this Pol3 variant in the complementation assay, we could not further test this hypothesis, compelling a move to other expression approaches.

We first attempted *E. coli*-based expression of the necessary proteins. Though *E. coli* expression of wild-type Pol3, an exonuclease deactivated Pol3, and a catalytically inactive Pol3 all were successful, attempts to express functional EF-hand Pol3 variant in *E. coli* failed multiple attempts (Figure S27 for one example), perhaps due to host interactions. Recent work showed that the yeast genome could be replicated replacing Pol3 with bacteriophage polymerase RB69^101^. If Pol3 possesses an interchangeable function with a native *E. coli* polymerase, the EF-hand variant’s probable higher error rate because of impaired exonuclease function or its likely slower replication speed could be causing cell stress. Therefore, we moved to cell-free *in vitro* expression with NEB PURExpress, using the same *E. coli* codon optimized and N- and C-terminally truncated variant described earlier. With PURExpress we were able to successfully express all desired variants (Figure S28) and were able to use them in the fluorescent primer extension assay (Figure S29). Using the fluorescent extension assay, we found that while wild-type Pol3 is minimally affected by lower Ca^2+^ concentrations (400 μM), the EF-hand variant appears to be impacted (Figure 4c). In addition, at high Ca^2+^ concentrations (4 mM), though the wild-type Pol3 is impacted, the EF-hand variant is impacted to a greater extent (Figure 4c, Figure S30 for replicate data). Overall, this data indicates that we generated a Ca^2+^ sensitive Pol3 variant, with the functional impact likely involving the region of mutation (exonuclease domain).

## Conclusion

In this work we demonstrate multiple approaches to implementing Ca^2+^ sensitivity to DNA replication, targeting elements of the Pol δ DNA polymerase complex from *Saccharomyces cerevisiae*. We achieved Ca^2+^ sensitivity by multiple engineering approaches, including utilizing the accessory protein PCNA, even demonstrating reversibility in this case. Moreover, through both domain insertion of CaM and binding site grafting of the EF-hand motif, we show direct modulation of the catalytic subunit Pol3’s function. This work not only serves as a template for engineering other ligand responsive protein domains, but represents an important step forward toward more direct signal transduction in DNA-based biosensing and recording. Our findings complement our previous work engineering TdT^59^ and provide a template-based alternative that would produce double-stranded DNA as an output.

Future work with these constructs should involve identifying an appropriate polymerase to pair with the engineered variants, one that has a disparate error rate and is especially insensitive to Ca^2+^. In addition, future work should involve improving Pol3’s replication speed and processivity. Increasing nucleotide incorporation rate will help improve recording resolution^102^. Improvements of the fundamental kinetics of the engineered DNAP and pairing with an appropriate partner DNAP will enable new biosensing applications where present technologies are limited. One example is in measurement of neuronal activity. The mapping of the brain could enable new treatments for neurological pathologies and insight into emergent behavior, but current neuronal measurement technologies are limited in their space-time resolution^103^ and prevent realization of this goal. Whereas technologies such as MRI can cover the entire brain, they are too slow to capture activity as it happens^104^. Meanwhile, fluorescent microscopy and patch clamp technologies allow for real-time measurement, but they cannot cover the entire brain at once^105^, in part due to the density of brain tissue. Nanoscale molecular devices, such as our engineered DNAP, have been proposed as a solution^61^. Such approaches could leverage the spike in intracellular Ca^2+^ concentration concurrent with neuronal firing as a proxy measurement for neuronal activity, enabling the collection of real-time neuronal activity data across the entire brain. Although much work remains to achieve such a goal, here we establish an engineering foundation for such a technology, avoiding the pitfalls of approaches that involve transcription and translation, as a first step toward enabling applications such as neural recording.

## Supporting information

Cloning files

Table of Primers

## Abbreviations

DNAP: DNA polymerase
PCNA: proliferating cell nuclear antigen
TdT: Terminal deoxynucleotidyl transferase
CaM: calmodulin
F#: engineered fusion protein, with subsequent description describing if the fusion was with PCNA or Pol3.

## Author contributions

BWB, ADP, NJB and KEJT wrote the manuscript. BWB, ADP and NJB conducted experiments. BWB, ADP, NJB, TC, GMC, KEJT analyzed the experiments.

## Acknowledgements

We would like to thank Prof. Ed Boyden, Prof. Konrad Körding, Bradley Zamft, and Adam Marblestone for their feedback over the course of this work. We would like to thank Andrea Guerrero for her assistance in construct cloning. Sanger sequencing was supported by the Northwestern University NUSeq Core Facility. Gel imaging was supported by the Northwestern University Keck Biophysics Facility and a Cancer Center Support Grant (NCI CA060553). The Keck Biophysics Facility’s Azure Sapphire Imager was funded by a 1S10OD026963-01 NIH grant. Protein purification was supported by the Northwestern University Recombinant Protein Production Core. This work was funded by the National Institutes of Health grants R01MH103910 (to K.E.J.T. and G.C.), and UF1NS107697 (to K.E.J.T.) and a National Institutes of Health Training Grant (T32GM008449) through Northwestern University’s Biotechnology Training Program (to B.W.B).

