## Supplemental Materials for "Engineering Ca^2+^-dependent DNA polymerase activity"

#### Table of Contents

Methods and Materials ....................................................................................................3-8

Supplemental Figures......................................................................................................9-33

Supplemental Tables .......................................................................................................34-47

References.......................................................................................................................48

#### Methods and Materials

##### General cloning

Cloning was carried out in *Escherichia coli* DH5α unless otherwise noted. PCR was carried out primarily with New England Biolabs (NEB) Pfusion, Takara PrimeSTAR Max, NEB Q5, and Bio-Rad iProof High-Fidelity according to manufacturer protocols, unless otherwise indicated. Cloning of the Pol δ complex and its subcomponents was primarily carried on the pET-28a(+) vector bearing kanamycin resistance with T7 expression. For the yeast complementation assay, the plasmid YcpLac33 housed the wild-type Pol3 and URA3 marker and YcpLac111 held the engineered Pol3 variant and the LEU2 marker. Transformation of the Pol3 variants for the yeast complementation assay was carried out in the *Saccharomyces cerevisiae* strain BY4741 (*pol3Δ0::kanMX*), following previously established lithium acetate protocols[1]. Both plasmids and the strain were generous gifts of Peter Burgers. Protein expression was carried out in BL21 *E. coli* derivatives as described below, including Rosetta(DE3), Arctic, and NEB T7 Express. Unless provided as gifts, all other genes were synthesized as gBlocks by Integrated DNA Technologies (IDT) or GeneArt from Thermo Fisher Scientific and were codon optimized for *E. coli* expression. Primers, including fluorescently labeled primers, were ordered from IDT. Point mutations and simple deletions or insertions were created by PCR and the NEB KLD kit according to manufacturer protocols. Gibson assembly was carried out with 2x NEB master mix according to manufacturer protocols.

##### Using SCHEMA to predict CaM insertion sites in PCNA

The computational algorithm SCHEMA was used as previously described[2]. The amino acid sequence of *S. cerevisiae* PCNA was extracted from RCSB Protein Data Bank (PDB) structure file 1PLR. A protein-protein BLAST was performed to identify five protein homologs with sequence identity greater than 55%. Clustal Omega[3] was used to perform a multiple sequence alignment between PCNA and the five selected homologs (Table S1)**.** The RASPP script was run four times with 5, 6, 7 and 8 selected crossover points for library generation. Each independent run yielded a list of 2-26 different libraries consisting of 5, 6, 7, or 8 crossover residues (Tables S4-S7). The unique set of crossover residues were examined for a high number of occurrences in the libraries. Next a short-listed number of residues were examined for high flexibility based on B-factor and the root mean square difference (RMSD) between a DNA-bound vs DNA-free protein structure from PDB. The insertion point (AA 107) that occurred the highest number of times from SCHEMA predictions and had the highest RMSD and B-factor was identified as the ideal calmodulin (CaM) insertion point.

##### Creating PCNA-CaM fusions

Expression plasmids harboring wild-type and fusion PCNA sequences were constructed using a pET-28a(+) vector backbone such that each fusion had an N-terminus 6x His tag. All coding sequences of interest were placed under T7 expression. *S. cerevisiae* PCNA (Thermo Fisher Scientific) and *Rattus norvegicus* CaM (Integrated DNA Technologies) gene sequences were commercially synthesized and codon optimized for *E. coli* expression. PCR was used to amplify parts of PCNA based on the CaM insertion sites chosen from the full-length synthetic PCNA gene. Additionally, several different linkers were added to both ends of CaM via PCR resulting in a LINKER-CaM-LINKER design. Overlap PCR was optimized for the three parts by using equimolar concentrations of each part of the final fusion. The protocol was as follows: initial denaturation at 98°C for 30 seconds, then 15 cycles of denaturation at 98°C for 10 seconds, annealing at 60°C for 30 seconds, extension at 72°C for 60 seconds and final extension at 72°C for 10 minutes. Following primer addition, a second stage of PCR was conducted as follows: initial denaturation at 98°C for 30 seconds, then 20 cycles of denaturation at 98°C for 10 seconds, annealing at 72°C for 10 seconds, extension at 72°C for 60 seconds, with a final extension at 72°C for 10 minutes. For 250 femtomoles of PCNA fragments the PCR was set up as follows: 10 µL 5X iProof HF Buffer, 1 µL 10 mM dNTPs, 2.5 µL 10 µM NB45 Forward Primer, 2.5 µL 10 µM NB46 Reverse Primer, 2.70 µL PCNA-Fragment 1, 5.52 µL of PCNA-Fragment 2, 2.5 µL of 100 µM GS-CaM-GS, 0.5 µL iProof DNA Polymerase, 22.78 µL water to a final volume of 50 µL.

All constructs (resulting from overlap PCR) were transformed into competent DH5α *E. coli* cells and plated on Luria-Bertani (LB) agar with 25 µg/mL kanamycin. Colonies were screened via Sanger sequencing to determine correct assembly and all final constructs were fully sequence verified. Later each clone was transformed into BL21(DE3) for expression and purification of the wild-type and fusion proteins.

**Purification of PCNA-CaM fusions with Pol δ (conducted with Northwestern Recombinant Protein Production Core)**
Protein over expressions were carried out in *E. coli* BL21 (DE3). For PCNA-CaM fusions and wild-type PCNA purification, N-terminally His-tagged (6x His) proteins were expressed in 50 mL cultures for small scale production. Overnight inoculates were used to start 50 mL cultures at 0.1 $OD_{600}$. After $OD_{600}$ reached 0.6-0.8, protein expression was induced by addition of 1 mM isopropyl β-D-1-thiogalactopyranoside (IPTG). Purification of His-tagged proteins was carried out with a Qiagen Ni-NTA spin kit. For large scale protein isolation, the same procedure was used for 1-liter cultures.

For the Pol δ complex, expression followed a workflow established by Finkelstein et al.[4], with minor changes. A two-plasmid system was used, wherein Pol3 and Pol31 were expressed from a single plasmid, while Pol32 was expressed from a separate plasmid and carried an N-terminal 6x His tag. After overnight growth of individual transformants in liquid medium, cultures were diluted into 4 liters of fresh selective medium and grown at 37°C to an $OD_{600}$ of 0.7, then placed on ice for 30 minutes, induced with 1 mM IPTG, and grown overnight at 15°C. Induced cultures were harvested by centrifugation, resuspended in 3 mL of Tris-sucrose (50 mM Tris, pH 7.5, 10% sucrose), and stored at -80°C. Frozen cell pellets were thawed, lysed then subjected to Ni-NTA purification. Purified proteins were stored in Tris-HCl buffer at pH 7.5 containing 5-10% glycerol based on the downstream application.

**Procedure for Western blots, including crosslinking**
Various concentrations of purified proteins stored in 50 mM HEPES Buffer (150 mM NaCl, pH 7.5) were treated at a final concentration of 0.2 mM ethylene glycol bis(succinimidyl succinate) (EGS) for 30 minutes on ice to allow for crosslinking. The crosslinking reaction was immediately quenched with 50 mM Tris at pH 7.5 for 15 minutes at room temperature. After another 15 minutes, the samples were boiled in the presence of loading buffer and run on an SDS-polyacrylamide gel electrophoresis (PAGE) gel. After fixing the gel overnight, western blot analysis was carried out and the samples were treated with anti-Histidine antibody and imaged using a film.

**Procedure for $Ca^{2+}$-induced gel shift**
The $Ca^{2+}$-dependent electrophoretic mobility shift was demonstrated by SDS-polyacrylamide gel electrophoresis essentially as described by Burgess et al.[5] with exceptions. First, the $Ca^{2+}$-free samples were boiled for 3 min in sample buffer (Laemmli, Bio-Rad) containing 5 mM EGTA before being loaded onto the gel. Second, the electrophoresis buffer for $Ca^{2+}$-free conditions contained 8 mM EDTA. Third, the gels in the presence of $Ca^{2+}$ contained 0.1 mM $CaC1_2$.

**Dynamic light scattering for measuring change in PCNA trimer diameter upon treatment with $Ca^{2+}$**

Purified wild-type PCNA or F5-PCNA-CaM (DKS) were diluted at 1 mg/mL in 10 mM Tris-HCL, 200 mM NaCl, pH 7.5 buffer containing 2.5% glycerol to a final volume of 200 μL. Initial controls were performed using the same buffer with 1 mg/mL of pure BSA. This helped establish the viscosity of the buffer as 0.97 and to determine the deviation of the measured diameter (6.8 nm) from the predicted diameter (7.5 nm). Wild-type PCNA diameter was measured at ~8.5 nm which is very close to the predicted diameter of ~9 nm (Figure S3). F5-PCNA-CaM (DKS) diameter was predicted to be ~9-11 nm with the expected diameter of ~9 nm. After treating with 12.5 μM $CaCl_2$, the diameter was measured at 1, 2, 5, 10, 20, 40 minutes after treatment. Following this, we treated the same sample with 125 μM EGTA and measured the diameter after 1, 5, 20 and 25 minutes of treatment. All measurements were taken at room temperature.

**Procedure for thermal shift assay**

We largely followed a previously described workflow[6]. Briefly, we suspended 2 μg of either wild-type or F5-PCNA-CaM (DKS) in 15 μl of 10 mM Tris-HCL, 200 mM NaCl, pH 7.5 buffer in triplicates. In addition, we treated the same amount of each protein with 0, 0.0625, 0.125, 0.25, 0.5, 1, 2 or 4 mM $CaCl_2$ or 0, 0.03125, 0.0625, 0.125, 0.25, 0.5, 1 or 2 mM EGTA, all in triplicate. We then measured the melting curve for each protein under the different conditions (Figure S31 for an example set of melting curves).

**Fluorescent DNA primer extension assay for PCNA-CaM fusion experiments**

A gel-based, fluorescent primer extension assay was developed to measure DNA polymerase activity. The primer extension assay consisted of a 5' FAM-labeled extension primer (with four 5' phosphorothioate bonds) and a 5' TAMRA-labeled DNA template (with four 5' phosphorothioate bonds and a 3' dideoxy-C modification) that were commercially synthesized by Integrated DNA Technologies (Table S8). The 5' FAM-labeled extension primer was annealed to the 5' TAMRA-labeled DNA template by incubation at 95°C for 2 minutes, followed by a -0.1°C/sec ramp until reaching 4°C. The primer and template were annealed in a 1.5:1 ratio of 375 nM primer: 250 nM template. Annealed primer/template was kept on ice and protected from light until use. Extension reactions were prepared in 50 μL volumes and consisted of 1X Pol δ buffer (40 mM Tris-HCl at pH 7.5 and 1 mM Dithiothreitol [DTT]), 1 mM $MgCl_2$, 200 μM dNTPs, 300 nM of Pol δ and 300 nM wild-type PCNA or F5-PCNA-CaM. For $Ca^{2+}$ conditions, $CaCl_2$ was added to the extension reaction at a final concentration of 4 mM. All reactions were prepared on ice. To initiate extension, 3.1 μL of the annealed primer/template was added to each reaction. Reactions were incubated at 30°C for 3 hours while being protected from light. After extension, reactions were covered in aluminum foil and stored at -20°C until gel electrophoresis.

**Using SCHEMA to predict CaM insertion sites in Pol3**

The computational algorithm SCHEMA was used as previously described[2]. The amino acid sequence of the catalytic subunit of *S. cerevisiae* DNA polymerase δ (Pol3) was extracted from RCSB Protein Data Bank (PDB) structure file 3IAY[7]. Consistent with the crystal structure, the Pol3 sequence consisted of N- and C-terminal truncations (amino acids 1-66 and 986-1097 were deleted) to produce a minimal catalytic core, $Pol3_{67\text{-}985}$, since full-length Pol3 is prone to aggregation. In this study, we refer to $Pol3_{67\text{-}985}$ as Pol3, since the truncated and full-length versions were previously found to be catalytically equivalent[7]. Next, a protein-protein BLAST was performed to identify five protein homologs with sequence identity greater than 60% (Table S9). Clustal Omega[3] was used to perform a multiple sequence alignment between Pol3 and the five selected homologs (Table S10). The N- and C-termini were also trimmed from the five selected homologs to improve alignment with the PDB-derived Pol3 sequence. The RASPP script was run three times with 4, 5, and 7 selected crossover points for library generation. Each independent run yielded a list of 35-37 different libraries consisting of 4, 5, or 7 crossover residues (Tables

S11-S13). The unique set of crossover residues was compiled for each run and ranked based on frequency. The crossover residues appearing at least five times in a given run were compiled to yield a unique set of 19 residues across all three *n*-crossover runs (Table S2 and S14). These 19 SCHEMA-predicted Pol3 residues served as potential CaM insertion sites.

**Pol3 expression plasmid construction**

As mentioned, expression plasmids harboring wild-type and variant Pol3 sequences were constructed using a pET-28a(+) vector backbone as the starting point for cloning. All coding sequences of interest were placed under T7 expression. *S. cerevisiae* Pol3 (Thermo Fisher Scientific) and *Rattus norvegicus* CaM (Integrated DNA Technologies) gene sequences were commercially synthesized and codon optimized for *E. coli* expression (Table S15). PCR was used to amplify N- and C- terminally truncated Pol3 (amino acids 1-66 and 986-1097 were deleted) from the full-length synthetic Pol3 gene as well as add a stop codon after residue 985. These truncations were consistent with a previously published Pol3 crystal structure (PDB ID: 3IAY)[7]. Additionally, glycine-serine-glycine-glycine-glycine (GSGGG) linkers were added to both ends of CaM via PCR resulting in GSGGG-CaM-GSGGG (Table S15).

Gibson assembly was used to create an expression vector, pET-GST-Pol3 (Figure S32), consisting of a glutathione S-transferase (GST) tag fused to the N-terminus of Pol3. A catalytically inactive variant of Pol3 ($Pol3_{Inact}$) was created using the Q5 Site-Directed Mutagenesis Kit (New England Biolabs) to introduce four separate point mutations (D608A, D764A, E800A, E802A) successively into the pET-GST-Pol3 construct[7,8] Pol3 EF-hand variants were created similarly, utilizing PCR with PrimeSTAR Max from Takara and NEB's KLD kit. To construct Pol3-CaM fusions, PCR was used to prepare GSGGG-CaM-GSGGG inserts and linearized pET-GST-Pol3 vector backbones for two-part Gibson assembly. Using standard methods[9], CaM inserts were assembled at the carboxyl end (just after the residue) of 19 selected Pol3 insertion residues in pET-GST-Pol3 to create 19 Pol3-CaM fusion constructs. Primer sequences used for Gibson assembly were commercially synthesized (Integrated DNA Technologies) and are listed in Table S16.

All constructs (resulting from Gibson assembly and site-directed mutagenesis) were transformed into competent DH5α *E. coli* cells and plated on LB agar with 25 μg/mL kanamycin. Colonies were screened via Sanger sequencing to determine correct assembly and all final constructs were fully sequence verified.

***In vitro* protein expression with NEB PURExpress**

Protein expression of Pol3 constructs was performed using the PURExpress *In Vitro* Protein Synthesis Kit (New England Biolabs). *In vitro* transcription/translation (IVTT) reactions were prepared according to manufacturer's guidelines. For each 25 μL reaction, 500 ng of ethanol precipitated plasmid was used as template and 20 units of Murine RNAse inhibitor (New England Biolabs) were added as a supplement. Reactions were incubated at 30°C for 3 hours and kept on ice. Fresh DNA polymerase was prepared for each downstream experiment.

**Fluorescent DNA primer extension assay for Pol3 variant experiments**

Using the same fluorescent primer extension assay described earlier for PCNA-CaM fusion experiments, activity of Pol3 variants (referring to Pol3-CaM fusions and Pol3 EF-hand variants) was tested. Reactions were set up identically, except only Pol3 (no PCNA or other accessory proteins) was included and the sourcing for Pol3 was IVTT expression. For $Ca^{2+}$ conditions of Pol3-CaM fusion experiments, $CaCl_2$ was added to the extension reaction at a final concentration of 4 mM. For experiments with Pol3 EF-hand variants, $CaCl_2$ was added to the extension reaction at a final concentration of either 400 μM or 4 mM. For Pol3-CaM fusion experiments, a synthetic CaM-binding peptide (ARRKWQKTGHAVRAIGRLSS) from smooth muscle myosin light chain kinase (smMLCK)[10], which we refer to as M13 peptide, was also included as a modulating condition. M13 peptide was prepared by dissolving 1 mg of lyophilized M13 peptide (synthesized and pre-aliquoted by GenScript) into 1 mL of sterile, nuclease-free water. For M13 peptide conditions, M13 peptide solution was added to the extension reaction at a final concentration of 400 nM. All M13 peptide conditions also included 4 mM $CaCl_2$ in the reaction. All reactions were prepared on ice. To initiate extension, 3.1 µL of the annealed primer/template was added to each reaction. Reactions were incubated at 30°C for 3 hours while being protected from light. After extension, reactions were covered in aluminum foil and stored at -20°C until gel electrophoresis.

**Molecular weight analysis of extension products**

Primer extension products were prepared for polyacrylamide gel electrophoresis (PAGE) by combining 8 µL of extension product with 12 µL of TBE-Urea Sample Buffer (Bio-Rad) and boiling for 15 minutes at 100°C. Samples were immediately transferred to ice to cool down after boiling and covered with aluminum foil. Next, 13 µL of each prepared sample was loaded onto a precast 15-well, 10% polyacrylamide TBE-Urea gel (Mini-PROTEAN, Bio-Rad). A maximum of 8 samples were run per gel, with skipped lanes to reduce cross-contamination during gel loading. Samples were run in 1X TBE buffer at 200V for 40 minutes under low-light conditions.

Samples were imaged with a Typhoon 9400 Variable Mode Imager (GE Healthcare) or Azure Sapphire Biomolecular Imager (Azure Biosystems) under fluorescent mode. Samples from PCNA-CaM and Pol3-CaM fusion experiments were imaged with a Typhoon 9400 Variable Mode Imager and the following settings were applied: For FAM imaging, an excitation wavelength of 488 nm and an emission filter at 520 nm with a 40 nm bandpass was used. For TAMRA imaging, an excitation wavelength of 532 nm and an emission filter at 580 nm with a 30 nm bandpass was used. Normal sensitivity and a PMT voltage of 600 volts was maintained for all scans. ImageQuant TL 1D gel analysis was used to quantify the intensity of fluorescence for FAM-labeled extension products. Using 1D gel analysis software, lanes were automatically created from the whole gel and adjusted as necessary. Background fluorescence was calculated and subtracted using the rolling ball method (radius set to 20). For band detection, the following parameters were used: minimum slope of 102, noise reduction of 0, percentage maximum peak of 1%, and automatic edge detection. Background-subtracted volumes at the band of interest were used to calculate relative changes in activity for a given DNA polymerase. Fold change calculations were limited to samples within the same gel to control for gel-to-gel variability.

Samples from Pol3 EF-hand variant experiments were imaged with an Azure Sapphire Biomolecular Imager and the following settings were applied: For FAM imaging an excitation wavelength of 488 nm was used. Standard sensitivity and a voltage of 600 was maintained for all scans. For intensity measurement, the rolling ball method was applied at a setting of 100. Automatic band detection was utilized, with a minimum band slope of 100. The band noise factor was set to 10, the percentage maximum peak was set to 10, and automatic edge detection was used. Band positions were not edited. and band positions were not edited.

**Yeast complementation assay**

The yeast complementation assay for screening the viability of Pol3 EF-hand mutants was carried out as described previously[1], with adjustments to the specific needs of this Pol3 assay. Briefly, the standard lithium acetate yeast transformation protocol was employed to transform yeast with Pol3 mutants. The *S. cerevisiae* strain BY4741 (*pol3Δ0::kanMX*) was used for exchanging the wild-type Pol3 for the engineered variant to test variant viability. As this strain lacks a chromosomal copy of Pol3 it requires the Pol3 on the plasmid to grow. In the initial state, wild-type Pol3 is provided by the plasmid YcpLac33, which contains the URA3 marker and can be grown on uracil dropout media (CSM – Ura), but this is not necessary as the strain requires the Pol3 enzyme to grow and so precultures were grown in YPAD yeast medium. To force the exchange, transformed yeast was plated on CSM – Leu + 5-FOA to both require the retention of the plasmid bearing the variant Pol3 (YcpLac111, which contains the LEU2 marker) and the ejection of the wild-type plasmid as the URA3 gene utilizes 5-FOA to create a toxic byproduct.

As our initial cloning attempts of Pol3 variants on the YcpLac111 shuttle vector failed due to multiple gene deletions in Pol3 upon transforming into DH5α, we created a second vector with the fragment of interest (the Pol3 Exo domain, a 210 amino acid region from A305 to A514) cloned into the pET28 vector to perform mutations on just this fragment (pET28-Pol3-Exo). (Under the assumption that previous cloning of the *S. cerevisiae* Pol3 failed as the YcpLac111 vector gives leaky expression in *E. coli* and the Pol3 gene is toxic.) Once this second vector was established, all EF-hand variants and mutations could be created in the Pol3 Exo region. To incorporate these variants into the yeast complementation assay, we utilized yeast homologous recombination and separately prepared the "backbone" of YcpLac111 by "around the world" PCR, removing just the Exo region, and the insert. General Gibson assembly design was used to create the homologous recombination parts and the homology for both ends was ~30 nucleotides. For these transformations we added 1 μg of the backbone YcpLac111 and a 20:1 molar ratio of the insert fragment. After transformation, transformants were plated with spreaders and not glass beads. "Hits" were determined by colony formation within a one-week period with growth at 30°C.

#### Supplemental Figures

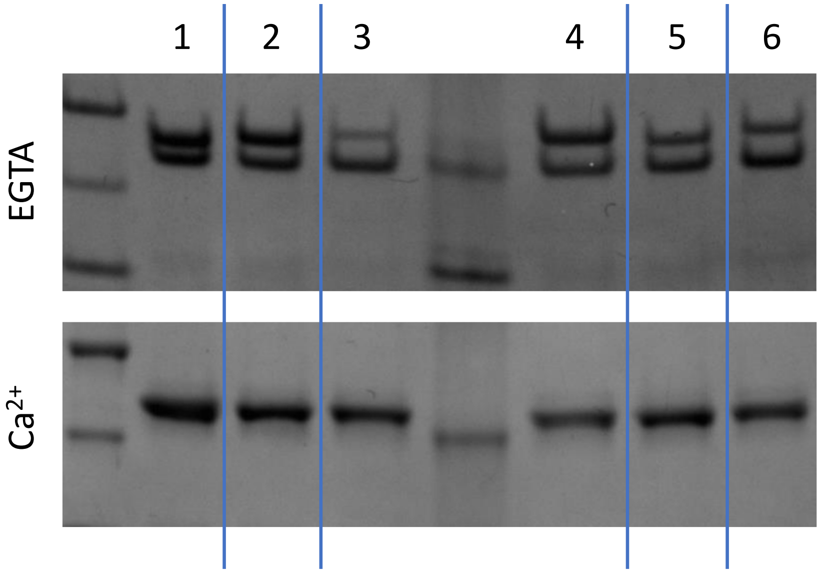


**Figure S1.** *Shifts of F1-F6 PCNA-CaM fusion variants upon EGTA or $Ca^{2+}$ addition*. Fusions all appear to show multiple bands in conditions with EGTA, but show a shift to a single band with $Ca^{2+}$ addition, possibly indicating a conformational change in the CaM domain upon binding to $Ca^{2+}$. #1 (F1-PCNA-CaM) represents the insertion of CaM after the K107 residue of PCNA using a flexible short “GS” linker. #2 (F2-PCNA-CaM) represents replacing the K107 residue with CaM using a “GS” linker. #3 (F3-PCNA-CaM) represents insertion of CaM after the K107 residue with a longer, more flexible “GSGGG” linker. #4 (F4-PCNA-CaM) represents insertion of CaM in the place of the K107 residue with a “GSGGG” linker. #5 (F5-PCNA-CaM) represents insertion of CaM after the K107 residue with short, inflexible “DKS” linker. #6 (F6-PCNA-CaM) represents replacement of the K107 residue with CaM using “DKS” linker.

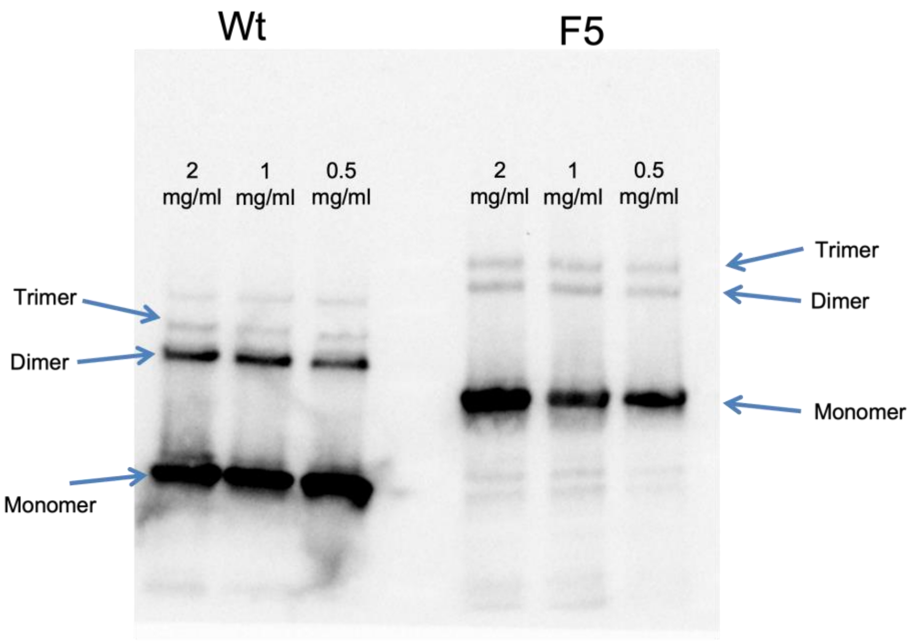


**Figure S2.** *Gel demonstrating that wild-type and engineered PCNA both form trimers*. As can be seen, both wild-type PCNA (Wt) and F5-PCNA-CaM (F5) are able to form dimers and trimers.

| | |
|---|---|
| Sample ID | Wt_2mMEGTA0.4mg/ml |
| Operator ID | Unknown Operator |
| Elapsed Time | 00:00:45 |
| Mean Diam. | 8.5 (nm) |
| Rel. Var. | 0.000 |
| Skew | 0.010 |
| RmsError | 1.7156e-02 |

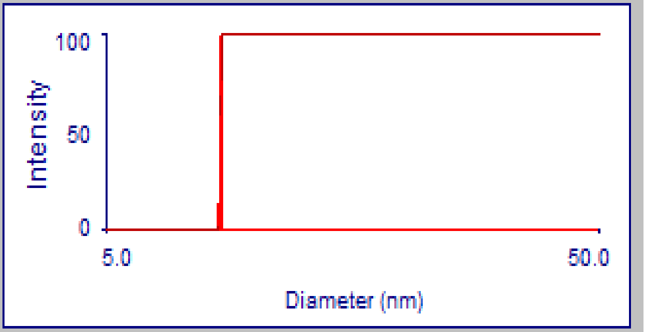


| d | G(d) | C(d) | d | G(d) | C(d) | d | G(d) | C(d) |
|---|---|---|---|---|---|---|---|---|
| 8.26 | 0 | 0 | 8.47 | 14 | 4 | 8.65 | 0 | 100 |
| 8.28 | 0 | 0 | 8.49 | 33 | 10 | 8.68 | 0 | 100 |
| 8.30 | 0 | 0 | 8.50 | 61 | 22 | 8.70 | 0 | 100 |
| 8.31 | 0 | 0 | 8.52 | 88 | 40 | 8.71 | 0 | 100 |
| 8.33 | 0 | 0 | 8.54 | 100 | 60 | 8.73 | 0 | 100 |
| 8.35 | 0 | 0 | 8.55 | 88 | 78 | 8.75 | 0 | 100 |
| 8.39 | 0 | 0 | 8.57 | 61 | 90 | 8.77 | 0 | 100 |
| 8.41 | 0 | 0 | 8.58 | 33 | 96 | 8.79 | 0 | 100 |
| 8.43 | 0 | 0 | 8.60 | 14 | 99 | 8.81 | 0 | 100 |
| 8.44 | 0 | 0 | 8.62 | 5 | 100 | 8.83 | 0 | 100 |
| 8.46 | 5 | 1 | 8.63 | 0 | 100 | 8.85 | 0 | 100 |

Print Window
Copy For Spreadsheet
Copy to Clipboard
Close

**Figure S3.** *DLS particle size distribution for wild-type PCNA in EGTA conditions*. A mean diameter of 8.5 nm is observed for PCNA in conditions with 0.4 mg/mL protein (~4 μM) with 2 mM EGTA added for 1 minute, which is very close to the expected 9 nm size for the PCNA trimer.

| | |
|---|---|
| Sample ID | Wt_4mMCaCl20.4mg/ml |
| Operator ID | Unknown Operator |
| Elapsed Time | 00:00:45 |
| Mean Diam. | 8.2 (nm) |
| Rel. Var. | 0.033 |
| Skew | -3.042 |
| RmsError | 2.3083e-02 |

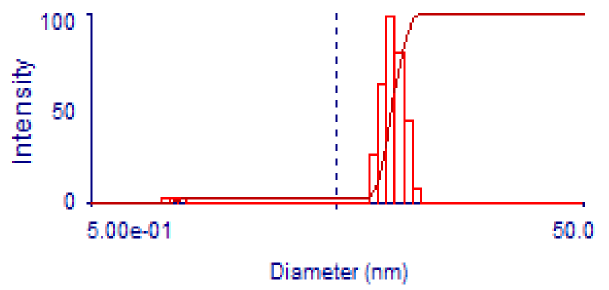


| d | G(d) | C(d) | d | G(d) | C(d) | d | G(d) | C(d) |
|---|---|---|---|---|---|---|---|---|
| 1.00 | 3 | 1 | 2.44 | 0 | 3 | 5.93 | 0 | 3 |
| 1.08 | 4 | 2 | 2.64 | 0 | 3 | 6.43 | 0 | 3 |
| 1.18 | 3 | 3 | 2.86 | 0 | 3 | 6.98 | 27 | 11 |
| 1.27 | 0 | 3 | 3.11 | 0 | 3 | 7.57 | 64 | 30 |
| 1.38 | 0 | 3 | 3.37 | 0 | 3 | 8.20 | 100 | 60 |
| 1.50 | 0 | 3 | 3.65 | 0 | 3 | 8.89 | 81 | 84 |
| 1.63 | 0 | 3 | 3.96 | 0 | 3 | 9.64 | 44 | 98 |
| 1.76 | 0 | 3 | 4.29 | 0 | 3 | 10.46 | 8 | 100 |
| 1.91 | 0 | 3 | 4.65 | 0 | 3 | 11.34 | 0 | 100 |
| 2.07 | 0 | 3 | 5.05 | 0 | 3 | 12.30 | 0 | 100 |
| 2.25 | 0 | 3 | 5.47 | 0 | 3 | 13.33 | 0 | 100 |

Print Window
Copy For Spreadsheet
Copy to Clipboard
Close

**Figure S4.** *DLS particle size for wild-type PCNA upon $Ca^{2+}$ addition*. A mean diameter of 8.2 nm is observed, which closely matched the mean diameter in the EGTA conditions (8.5 nm) and is close to the predicted size 9 nm. This indicates that the wild-type PCNA is unaffected by $Ca^{2+}$ addition. The condition shown is 0.4 mg/mL protein (~4 μM) with 4 mM $CaCl_2$ added for 1 minute.

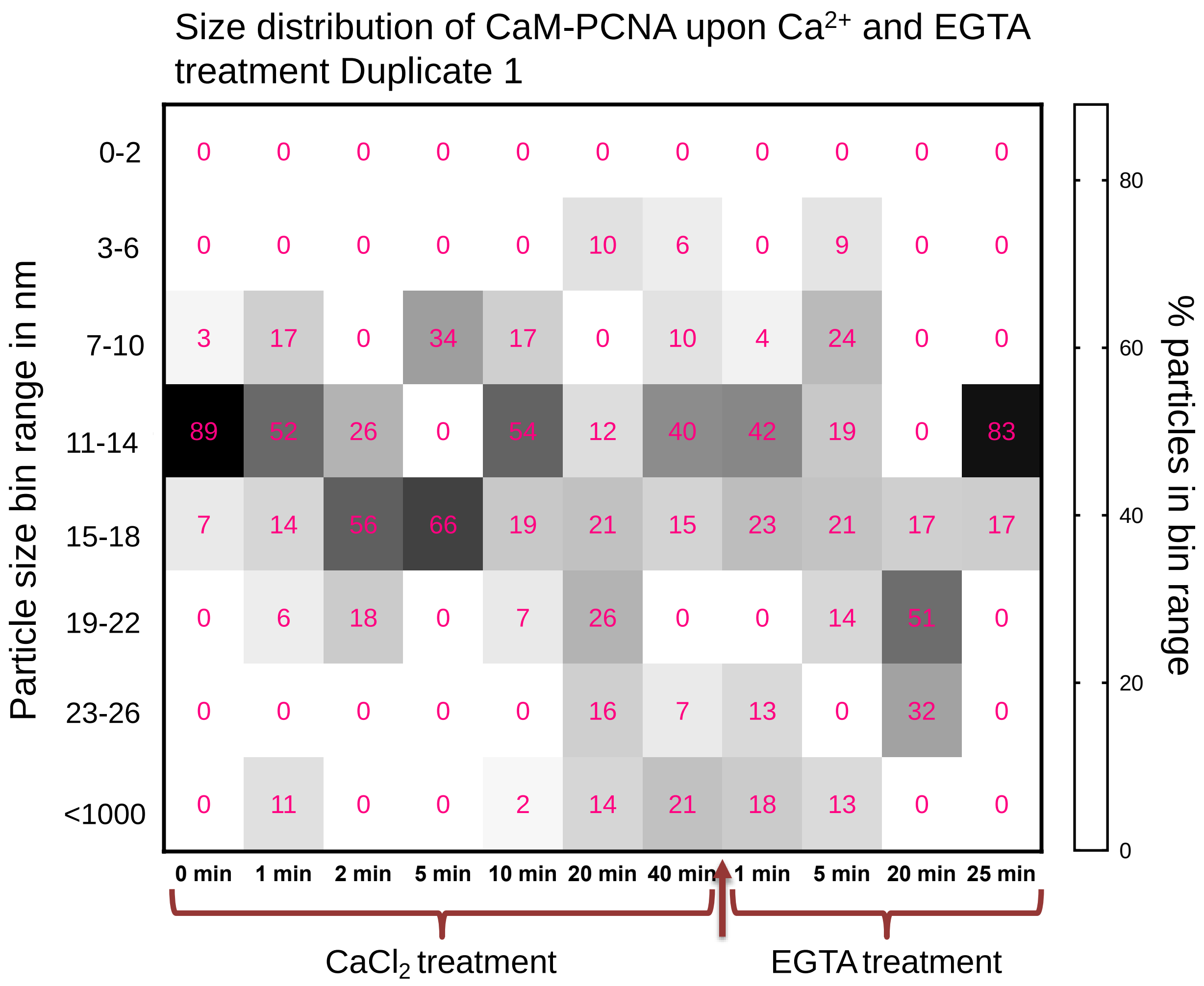


**Figure S5.** *Time course dynamic light scattering (DLS) data for run 1*.

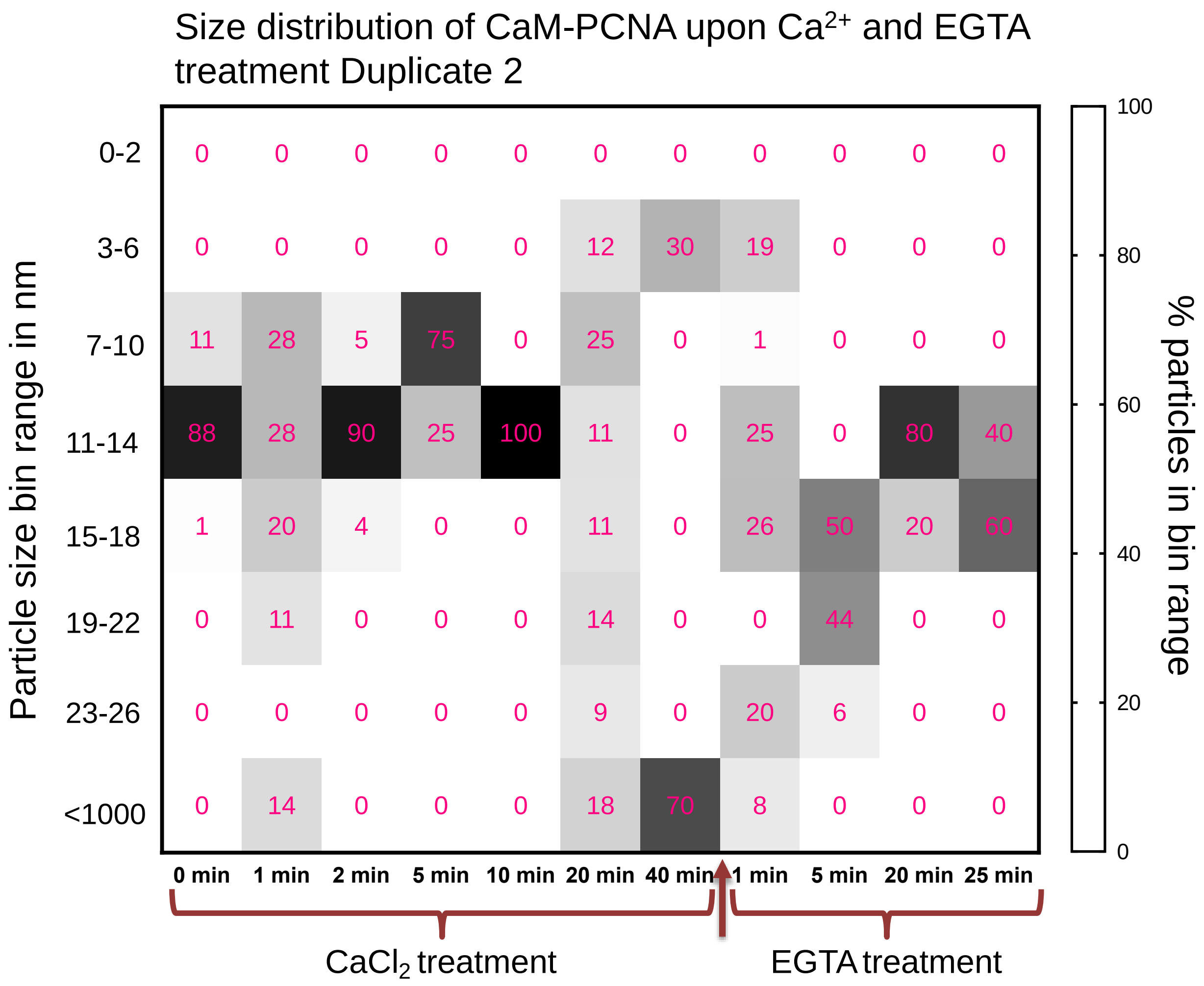


**Figure S6.** *Time course dynamic light scattering (DLS) data for run 2*.

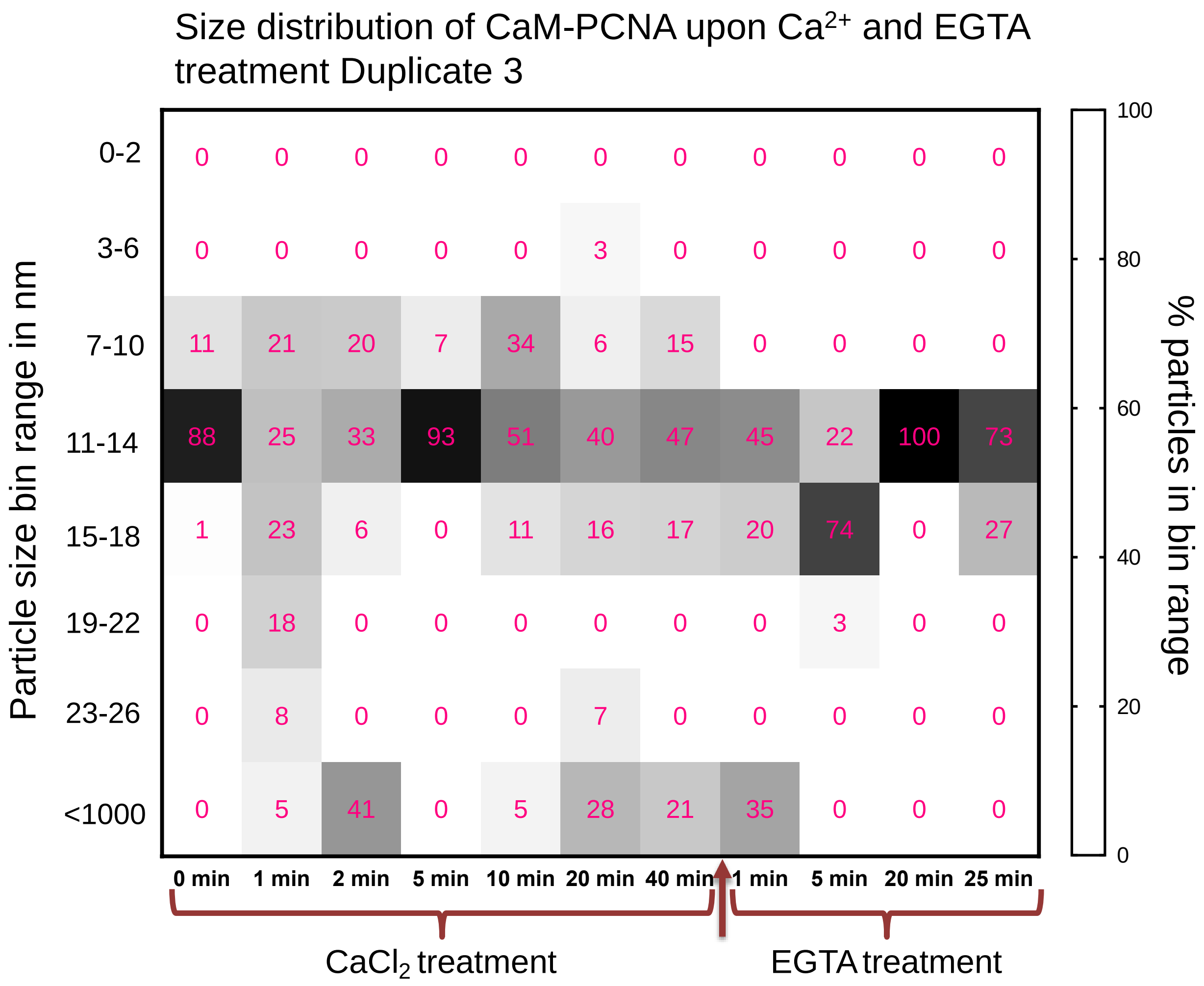


**Figure S7.** *Time course dynamic light scattering (DLS) data for run 3*.

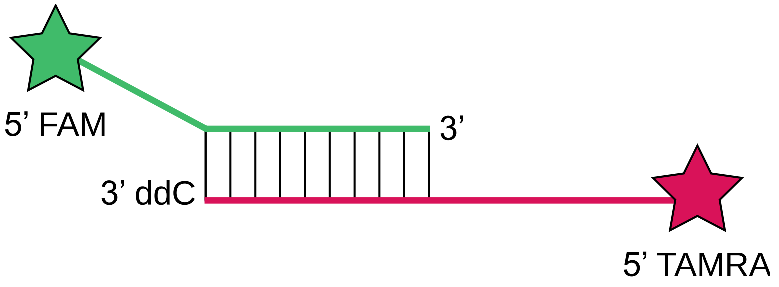


**Figure S8.** *Schematic for fluorescent primer extension assay*. FAM-labeled primer (green) is 60 bases, TAMRA-labeled template (dark pink) is 89 bases, overlap between primer and template is 25 bases, and extension products are 124 bases.

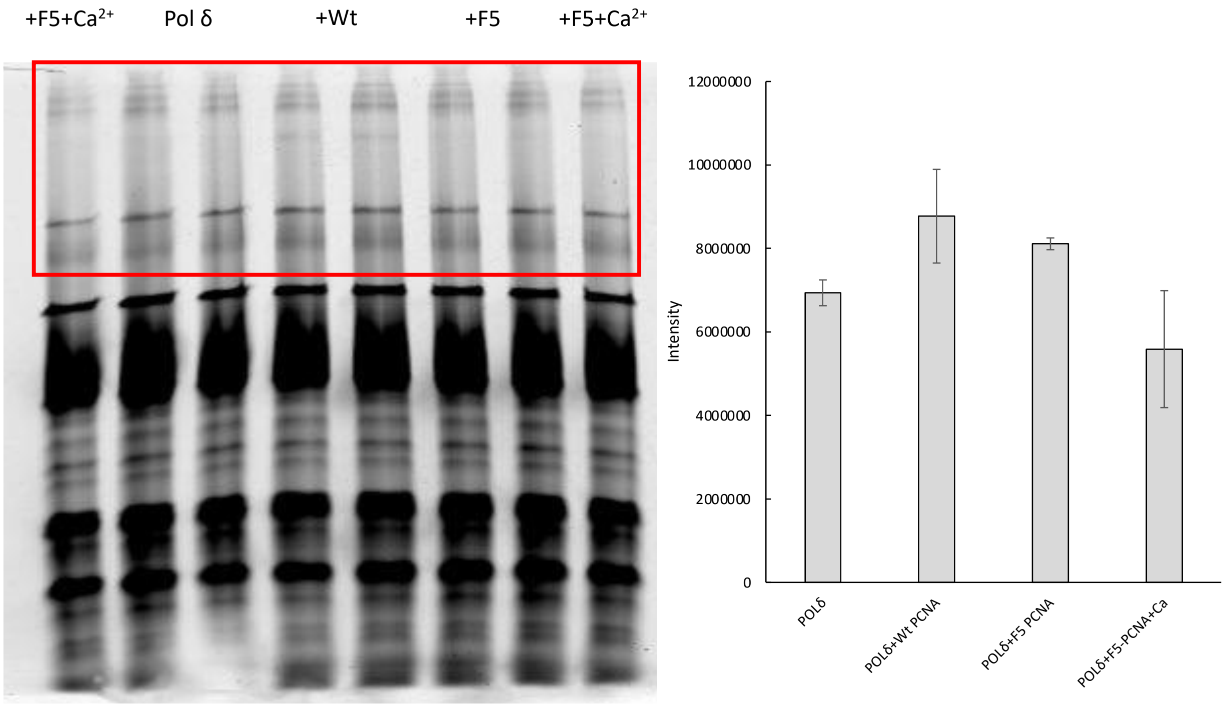


**Figure S9.** *PCNA fluorescent extension assay replicate* (1). $Ca^{2+}$ condition contains 1 μM $CaCl_2$. Wt = wild-type PCNA, F5 = F5-PCNA-CaM.

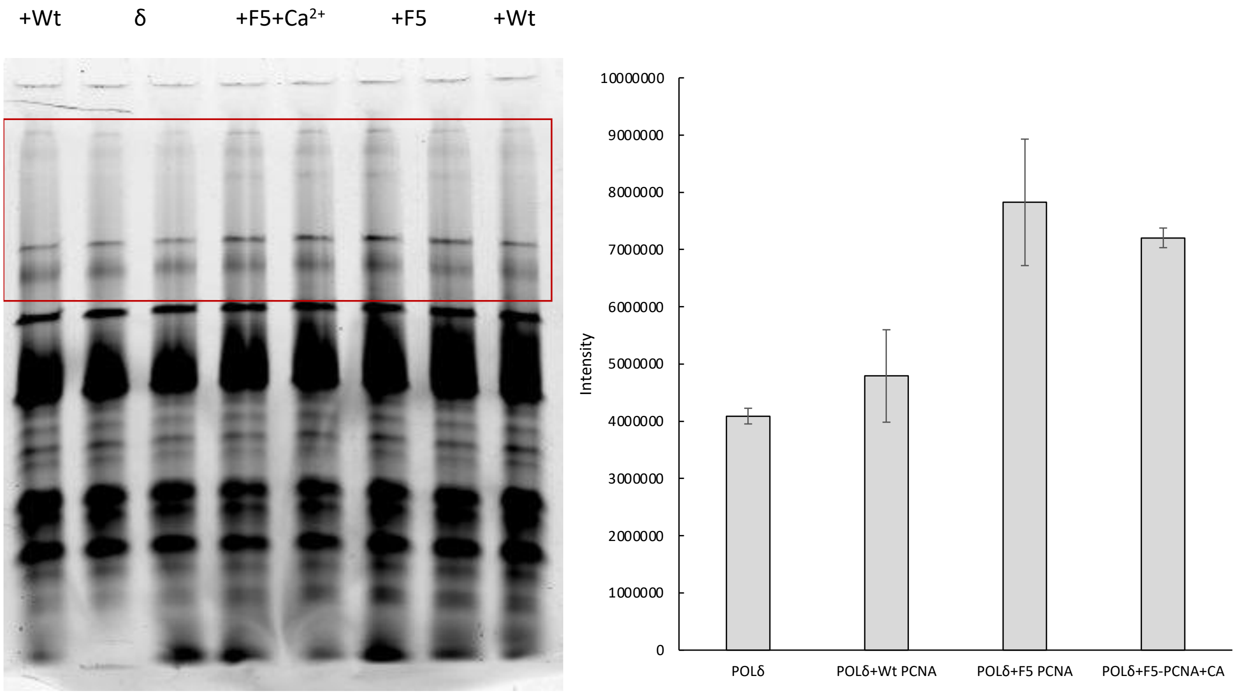


**Figure S10.** *PCNA fluorescent extension assay replicate* (2). $Ca^{2+}$ condition contains 1 μM $CaCl_2$. Wt = wild-type PCNA, F5 = F5-PCNA-CaM.

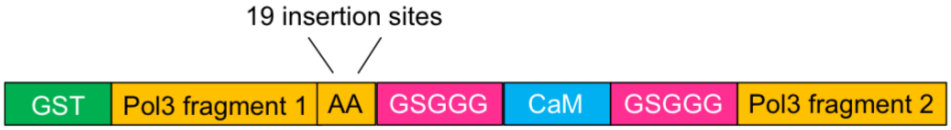


**Figure S11.** *Validation of SCHEMA-predicted Pol3 split sites through CaM insertion*. Pol3-CaM engineering strategy, where GSGGG-CaM-GSGGG is inserted directly at the carboxyl end of 19 different SCHEMA-predicted Pol3 amino acids (AA) to produce 19 distinct Pol3-CaM fusions.

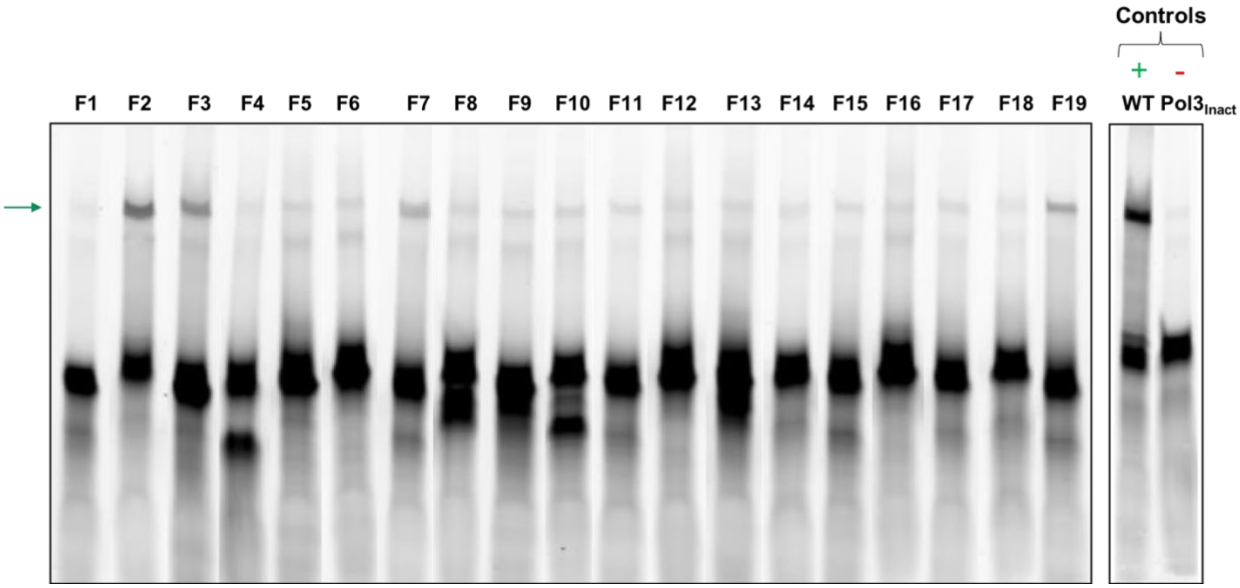


**Figure S12.** *Activity screening of F1-F19 Pol3-CaM fusions under standard $Mg^{2+}$ buffer conditions*. FAM-labeled extension products (124 bases, designated by green arrow) were resolved on 10% PAGE under denaturing conditions and FAM fluorescence was imaged ($\lambda_{ex}$ = 488 nm, $\lambda_{em}$ = 520 nm). Controls consisted of wild-type Pol3 (WT) and a catalytically inactive Pol3 variant ($Pol3_{Inact}$). Images of sample lanes from separate gels (Figures S13-S23) were stitched together to enable qualitative Pol3-CaM fusion activity comparison. A separate experiment (data not shown) where samples F1–F9 (referring to F1-Pol3-CaM–F9-Pol3-CaM) were run on the same gel revealed the same banding patterns shown above.

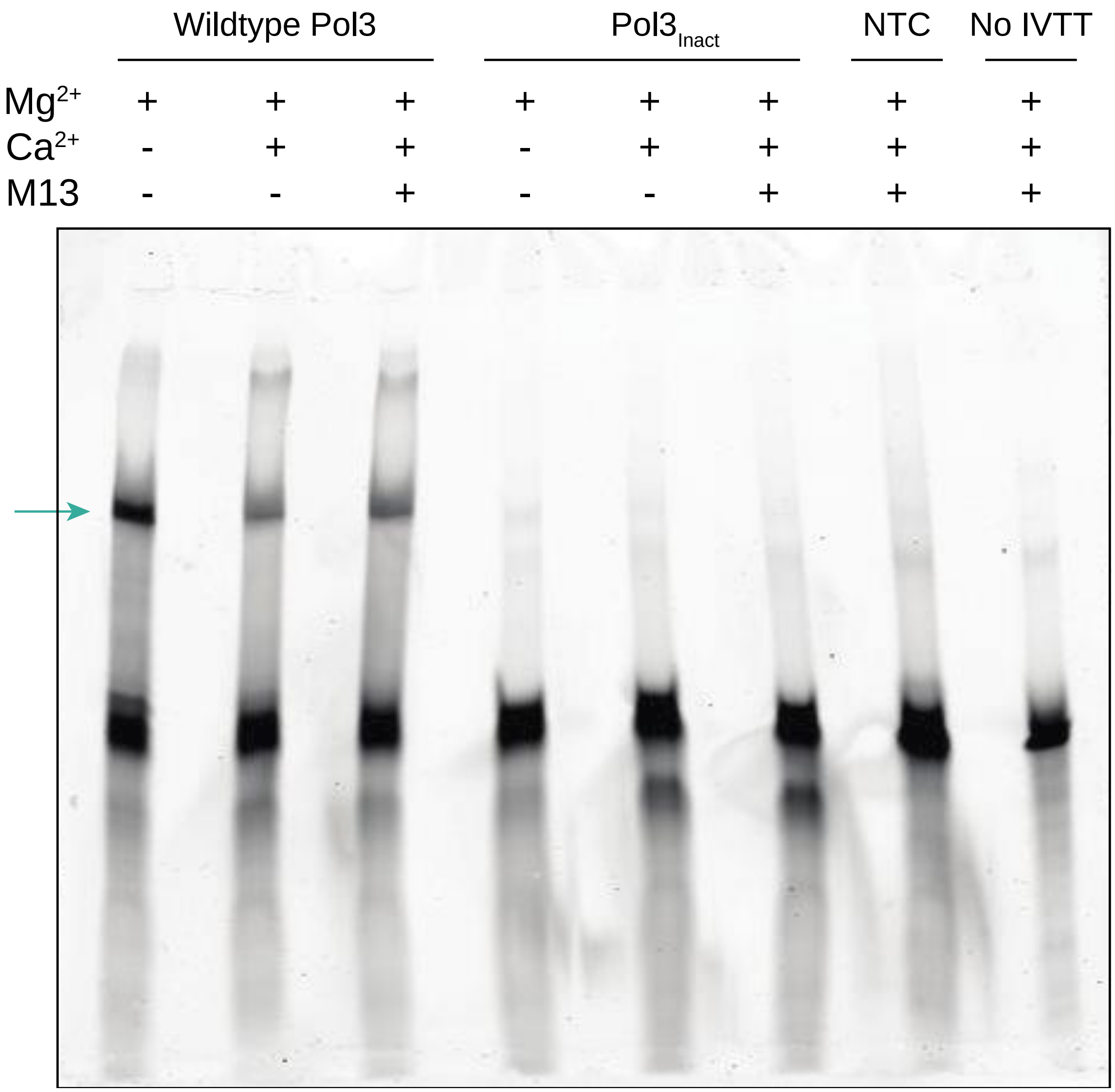


**Figure S13.** *Control samples for Pol3 activity testing.* Controls consisted of wild-type Pol3 (WT) and a catalytically inactive Pol3 variant ($Pol3_{Inact}$). Reactions were performed under $Mg^{2+}$, $Ca^{2+}$, and M13 peptide conditions. "NTC" is a no template control that refers to the absence of a Pol3 DNA template during *in vitro* transcription/translation (IVTT). "No IVTT" refers to a condition where the IVTT reagent (NEB PURExpress) was excluded from the primer extension reaction, to determine if there were any autofluorescence contributed from the kit components. FAM-labeled extension products (124 bases, designated by green arrow) were resolved on 10% PAGE under denaturing conditions. FAM fluorescence was imaged ($\lambda_{ex}$ = 488 nm, $\lambda_{em}$ = 520 nm). Lanes were skipped to prevent cross-contamination of samples during gel loading.

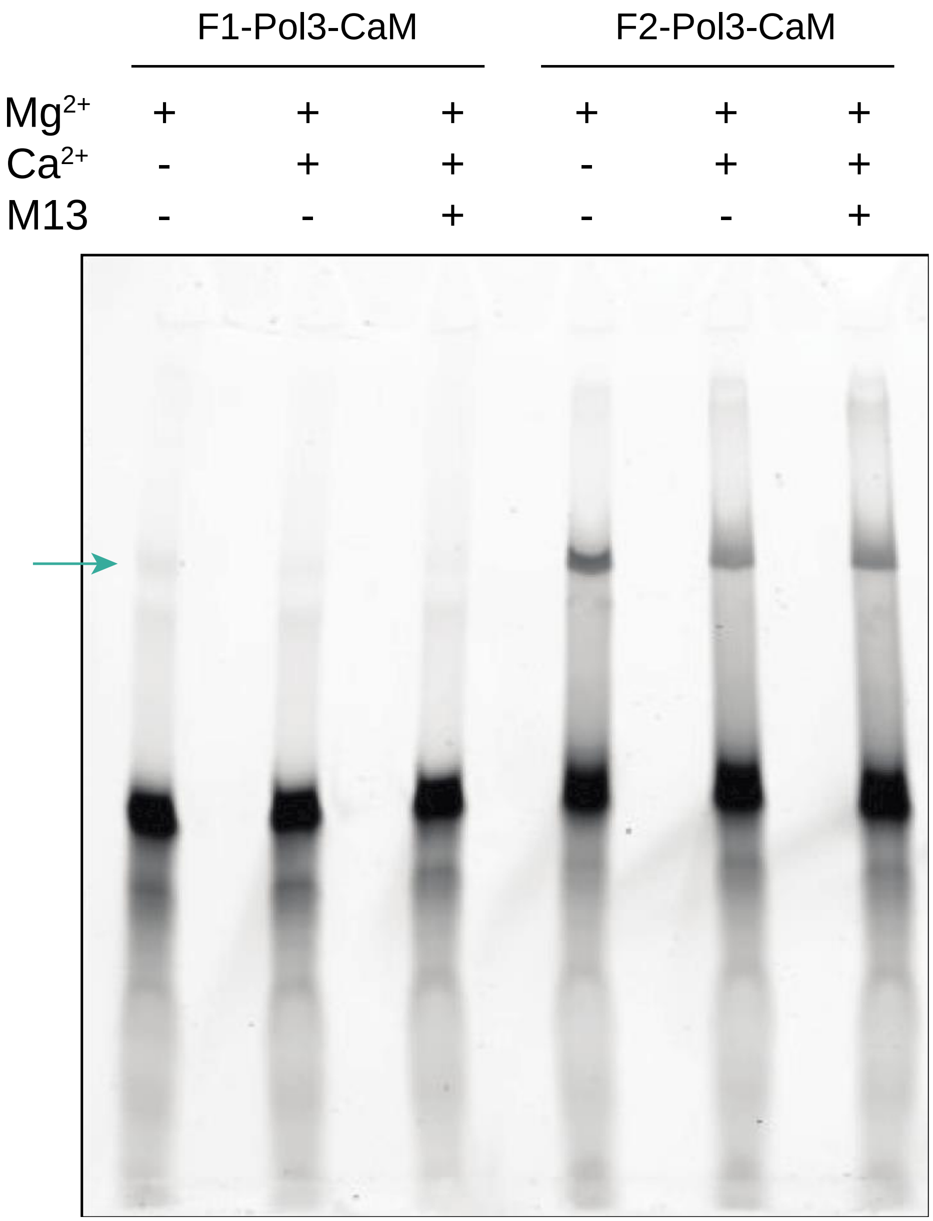


**Figure S14.** Activity testing of F1-Pol3-CaM and F2-Pol3-CaM. Reactions were performed under $Mg^{2+}$, $Ca^{2+}$, and M13 peptide conditions. FAM-labeled extension products (124 bases, designated by green arrow) were resolved on 10% PAGE under denaturing conditions. FAM fluorescence was imaged ($\lambda_{ex}$ = 488 nm, $\lambda_{em}$ = 520 nm). Lanes were skipped to prevent cross-contamination of samples during gel loading.

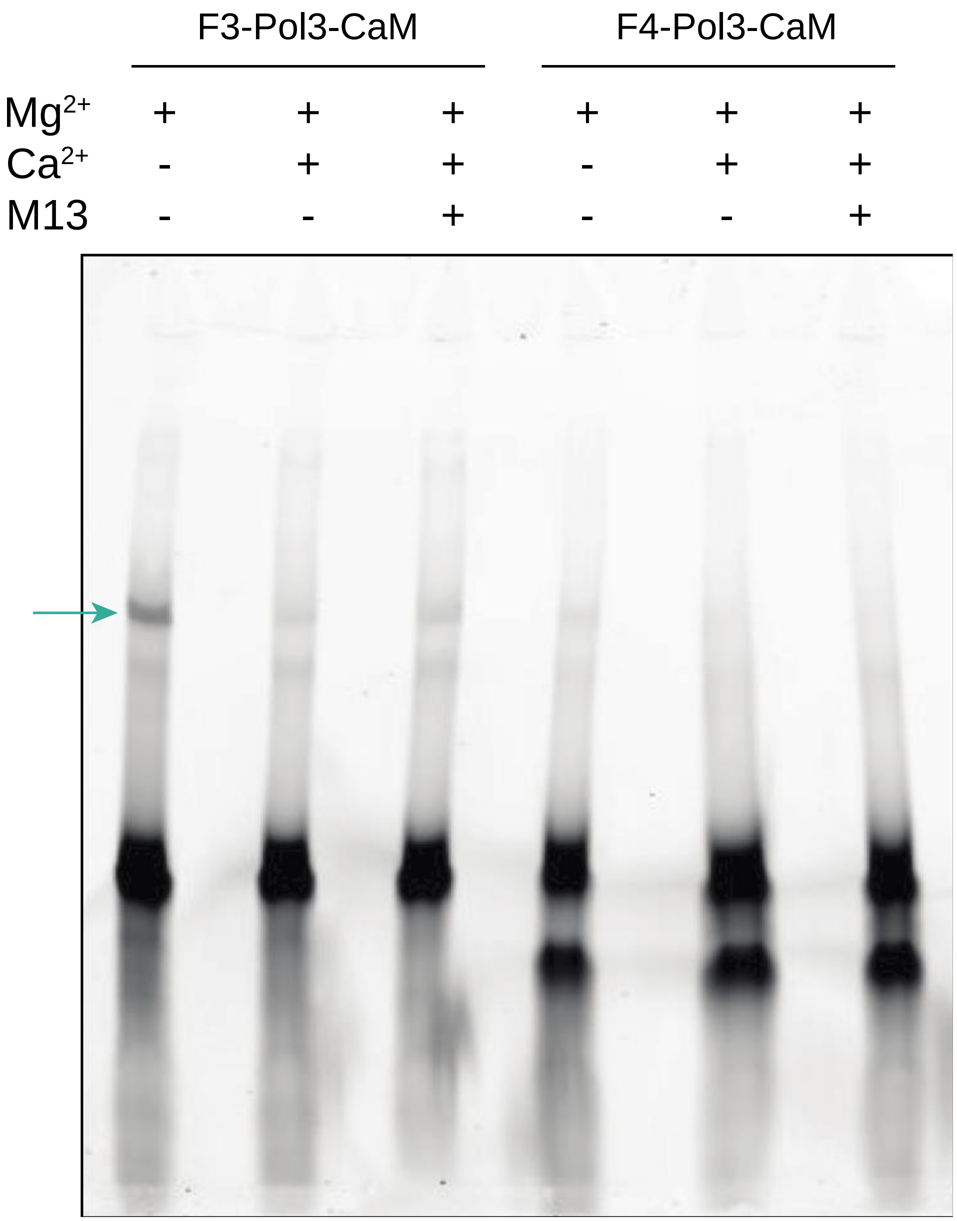


**Figure S15.** Activity testing of F3-Pol3-CaM and F4-Pol3-CaM. Reactions were performed under $Mg^{2+}$, $Ca^{2+}$, and M13 peptide conditions. FAM-labeled extension products (124 bases, designated by green arrow) were resolved on 10% PAGE under denaturing conditions. FAM fluorescence was imaged ($\lambda_{ex}$ = 488 nm, $\lambda_{em}$ = 520 nm). Lanes were skipped to prevent cross-contamination of samples during gel loading.

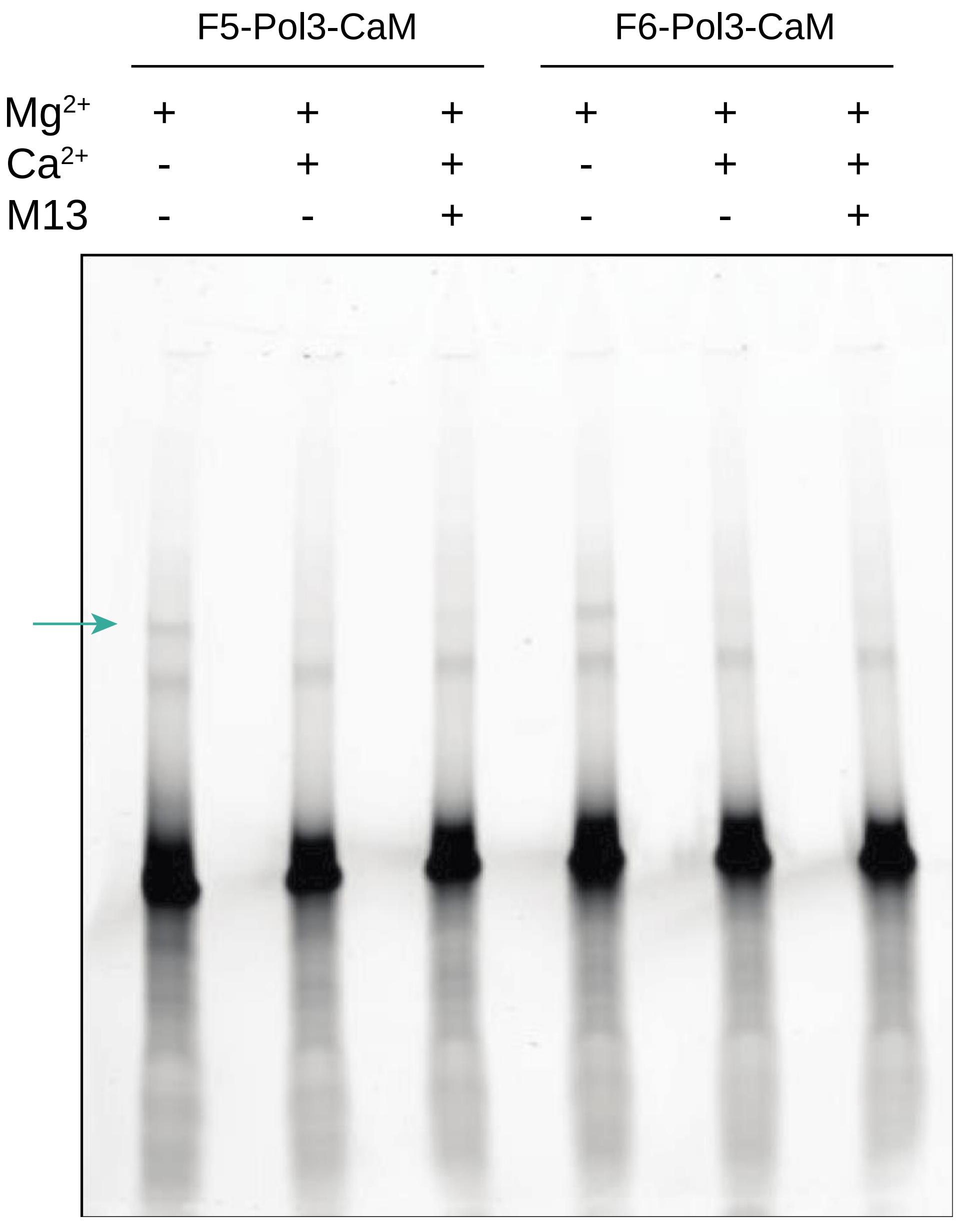


**Figure S16.** *Activity testing of F5-Pol3-CaM and F6-Pol3-CaM.* Reactions were performed under $Mg^{2+}$, $Ca^{2+}$, and M13 peptide conditions. FAM-labeled extension products (124 bases, designated by green arrow) were resolved on 10% PAGE under denaturing conditions. FAM fluorescence was imaged ($\lambda_{ex}$ = 488 nm, $\lambda_{em}$ = 520 nm). Lanes were skipped to prevent cross-contamination of samples during gel loading.

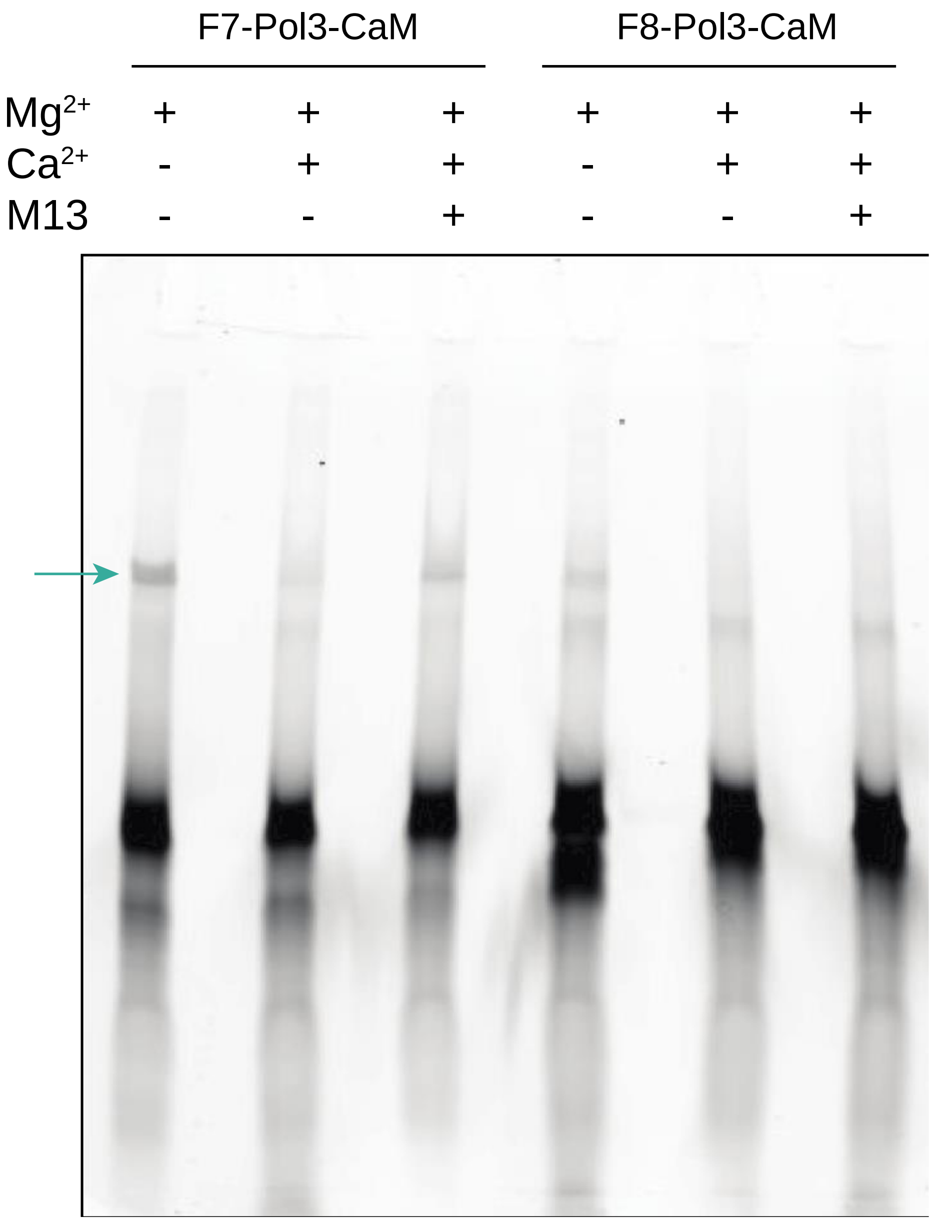


**Figure S17.** *Activity testing of F7-Pol3-CaM and F8-Pol3-CaM*. Reactions were performed under $Mg^{2+}$, $Ca^{2+}$, and M13 peptide conditions. FAM-labeled extension products (124 bases, designated by green arrow) were resolved on 10% PAGE under denaturing conditions. FAM fluorescence was imaged ($\lambda_{ex}$ = 488 nm, $\lambda_{em}$ = 520 nm). Lanes were skipped to prevent cross-contamination of samples during gel loading.

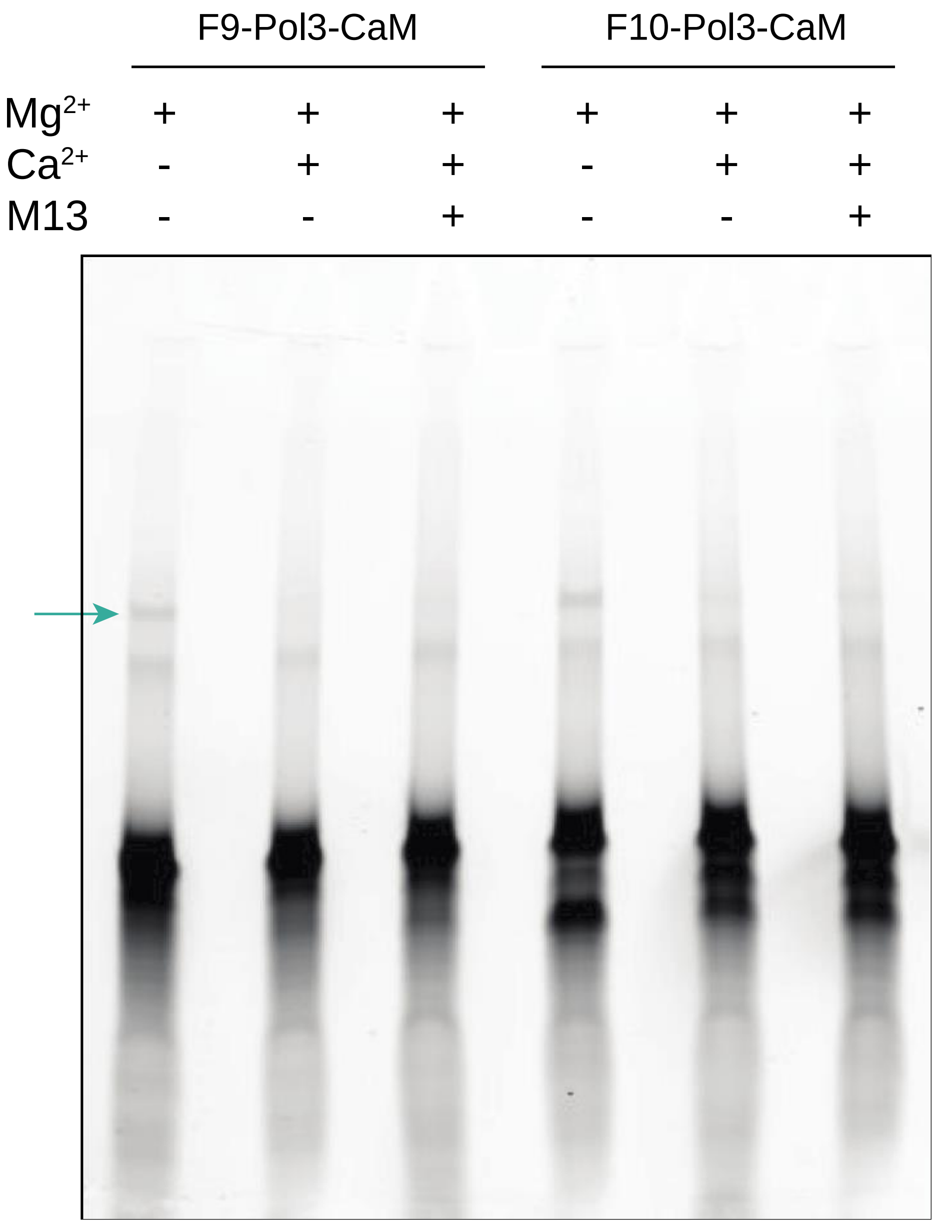


**Figure S18.** *Activity testing of F9-Pol3-CaM and F10-Pol3-CaM.* Reactions were performed under $Mg^{2+}$, $Ca^{2+}$, and M13 peptide conditions. FAM-labeled extension products (124 bases, designated by green arrow) were resolved on 10% PAGE under denaturing conditions. FAM fluorescence was imaged ($\lambda_{ex}$ = 488 nm, $\lambda_{em}$ = 520 nm). Lanes were skipped to prevent cross-contamination of samples during gel loading.

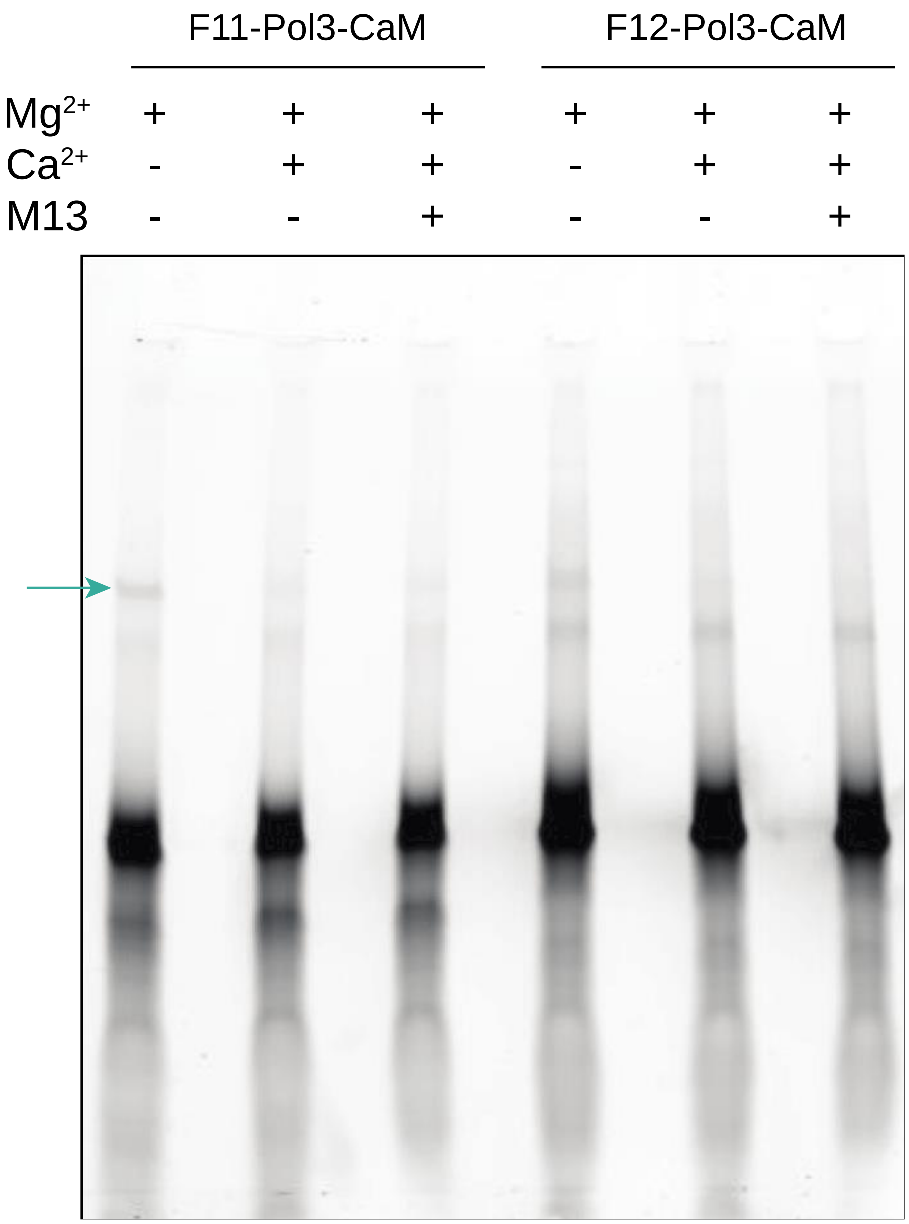


**Figure S19.** Activity testing of F11-Pol3-CaM and F12-Pol3-CaM. Reactions were performed under $Mg^{2+}$, $Ca^{2+}$, and M13 peptide conditions. FAM-labeled extension products (124 bases, designated by green arrow) were resolved on 10% PAGE under denaturing conditions. FAM fluorescence was imaged ($\lambda_{ex}$ = 488 nm, $\lambda_{em}$ = 520 nm). Lanes were skipped to prevent cross-contamination of samples during gel loading.

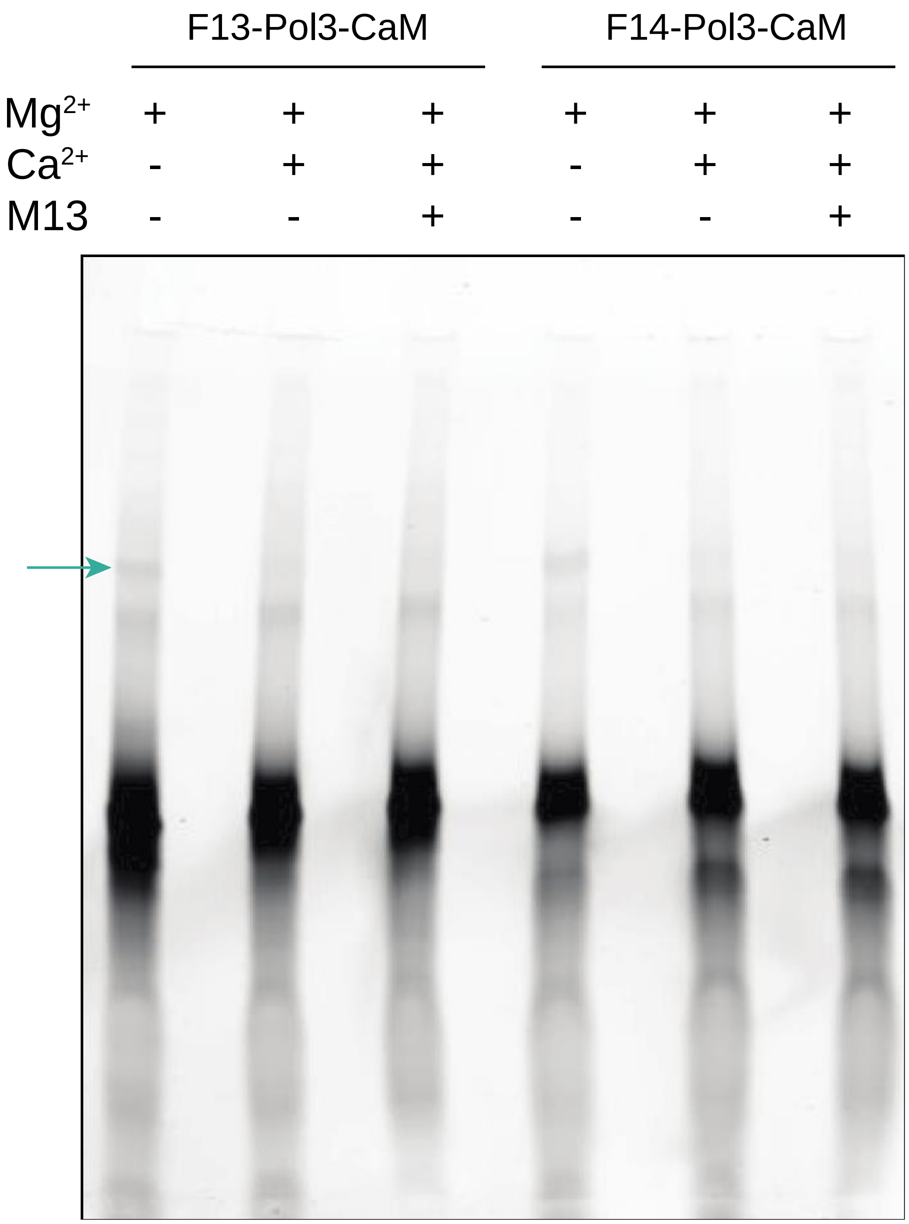


**Figure S20.** *Activity testing of F13-Pol3-CaM and F14-Pol3-CaM.* Reactions were performed under $Mg^{2+}$, $Ca^{2+}$, and M13 peptide conditions. FAM-labeled extension products (124 bases, designated by green arrow) were resolved on 10% PAGE under denaturing conditions. FAM fluorescence was imaged ($\lambda_{ex}$ = 488 nm, $\lambda_{em}$ = 520 nm). Lanes were skipped to prevent cross-contamination of samples during gel loading.

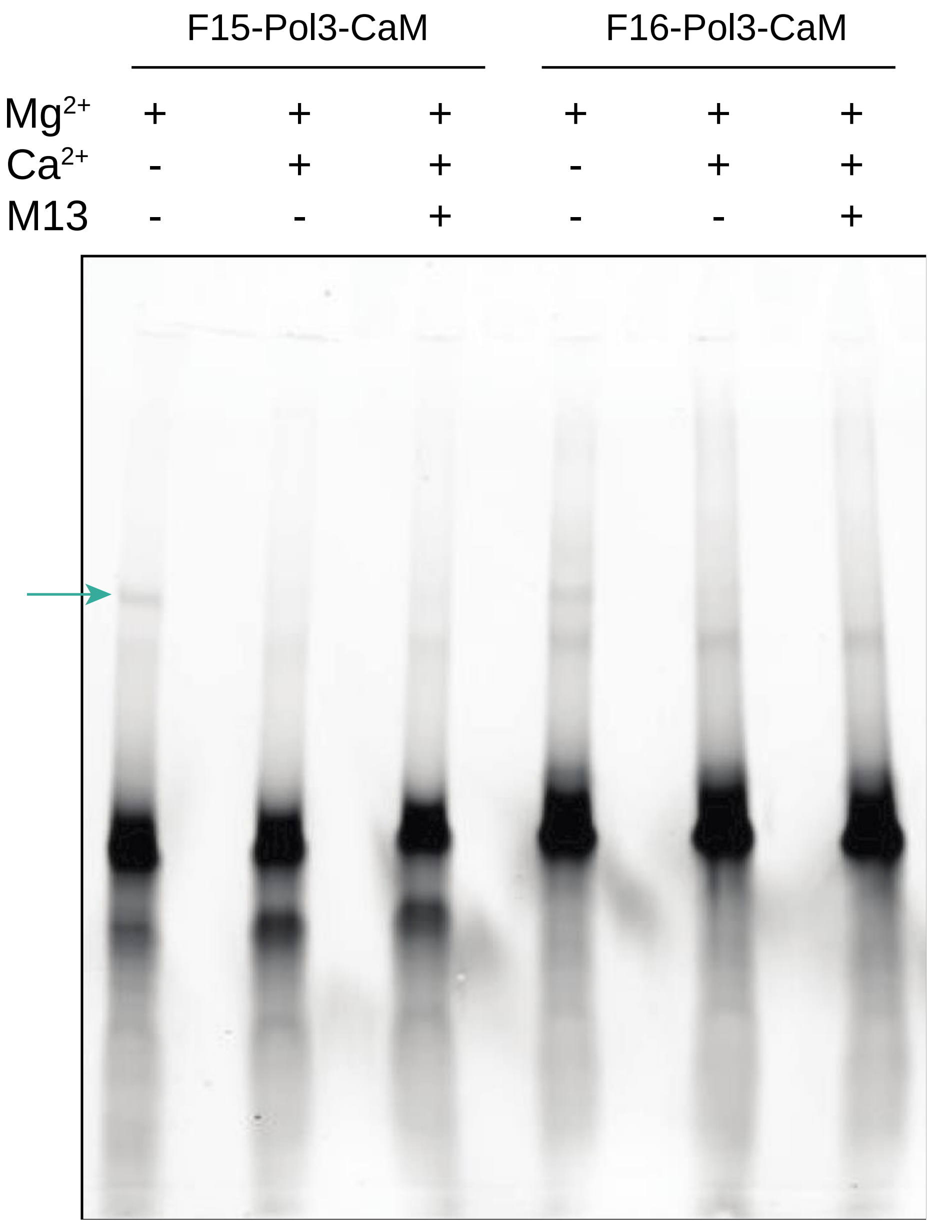


**Figure S21.** Activity testing of F15-Pol3-CaM and F16-Pol3-CaM. Reactions were performed under $Mg^{2+}$, $Ca^{2+}$, and M13 peptide conditions. FAM-labeled extension products (124 bases, designated by green arrow) were resolved on 10% PAGE under denaturing conditions. FAM fluorescence was imaged ($\lambda_{ex}$ = 488 nm, $\lambda_{em}$ = 520 nm). Lanes were skipped to prevent cross-contamination of samples during gel loading.

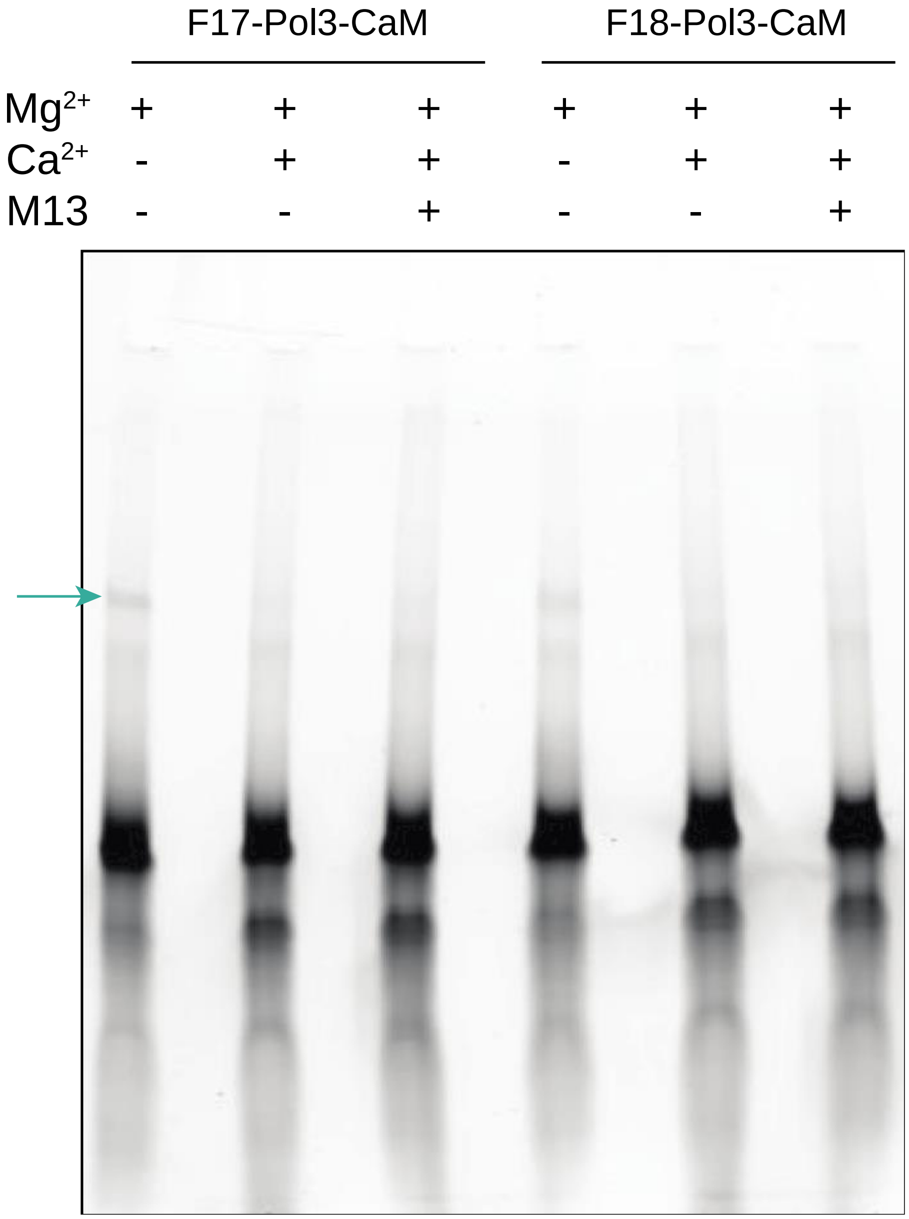


**Figure S22.** Activity testing of F17-Pol3-CaM and F18-Pol3-CaM. Reactions were performed under $Mg^{2+}$, $Ca^{2+}$, and M13 peptide conditions. FAM-labeled extension products (124 bases, designated by green arrow) were resolved on 10% PAGE under denaturing conditions. FAM fluorescence was imaged ($\lambda_{ex}$ = 488 nm, $\lambda_{em}$ = 520 nm). Lanes were skipped to prevent cross-contamination of samples during gel loading.

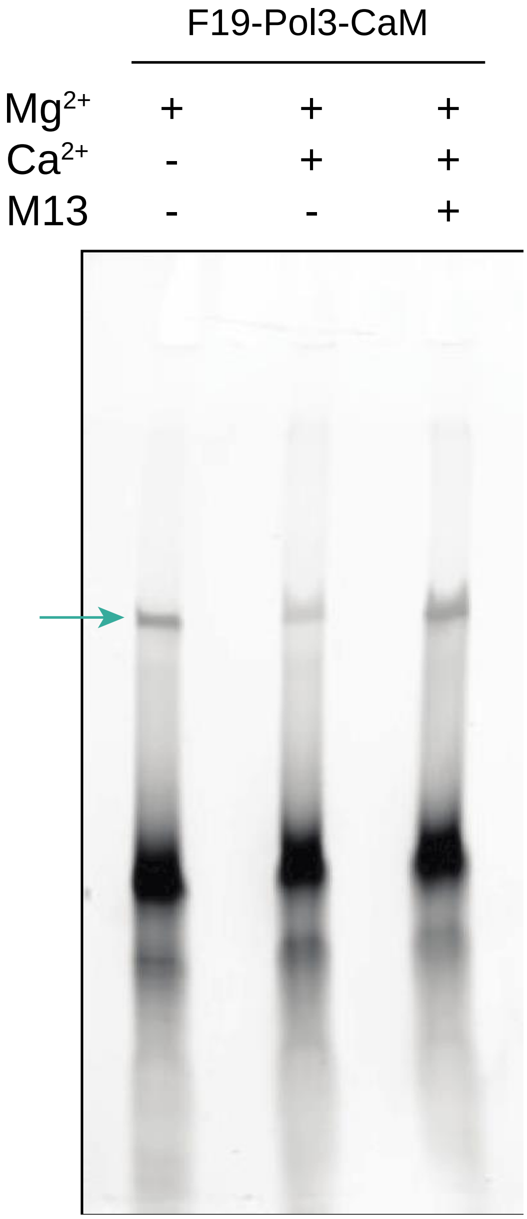


**Figure S23.** Activity testing of F19-Pol3-CaM. Reactions were performed under $Mg^{2+}$, $Ca^{2+}$, and M13 peptide conditions. FAM-labeled extension products (124 bases, designated by green arrow) were resolved on 10% PAGE under denaturing conditions. FAM fluorescence was imaged ($\lambda_{ex}$ = 488 nm, $\lambda_{em}$ = 520 nm). Lanes were skipped to prevent cross-contamination of samples during gel loading.

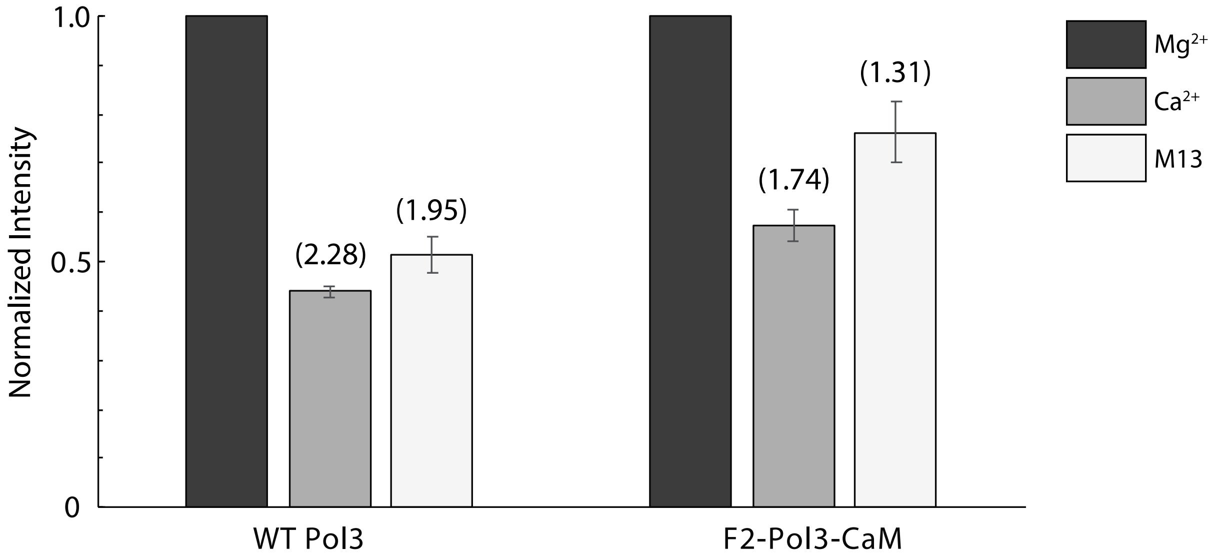


**Figure S24.** *Measuring activity modulation potential of F2-Pol3-CaM*. Fold change in activity of wild-type (WT) Pol3 and F2-Pol3-CaM under $Ca^{2+}$ and M13 peptide conditions (compared to the baseline $Mg^{2+}$ condition) was calculated (n=4, SEM plotted). Gel image analysis was performed to quantify intensities of extension products. Two independent extension reaction experiments were performed per condition and two technical replicates were imaged per experiment. Product intensities were background subtracted and used to calculate relative changes in activity for a given DNA polymerase. Fold change calculations were confined to samples within the same gel.

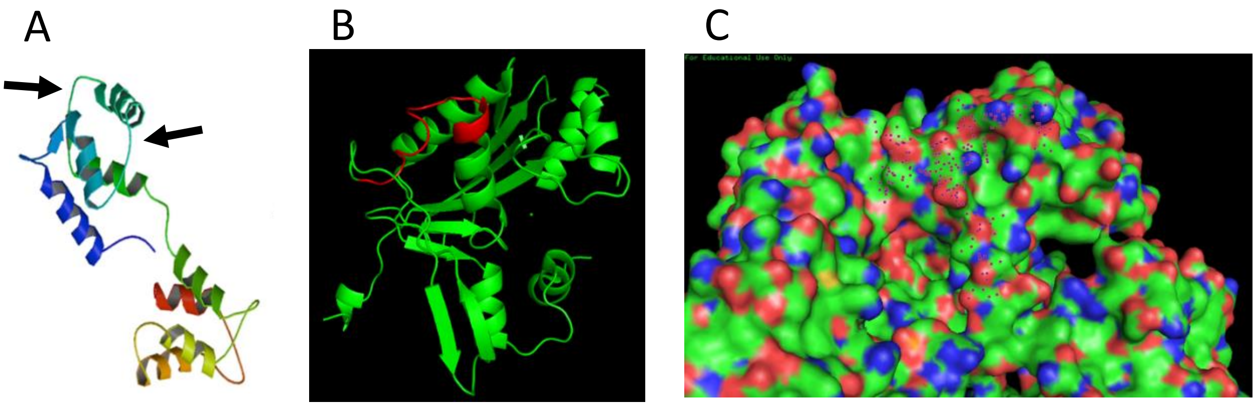


**Figure S25.** *Choosing a specific polymerase site to engineer for binding site grafting*. A) Arrows indicate on CaM the EF-hand region ($Ca^{2+}$ binding region) as seen in apo form[11]. B) Pol3's exonuclease domain, region highlighted in red, from 415-430, bears resemblance to the EF-hand $Ca^{2+}$ binding region (flexible loop exiting an α-helix). C) Demonstration that the region is solvent exposed.

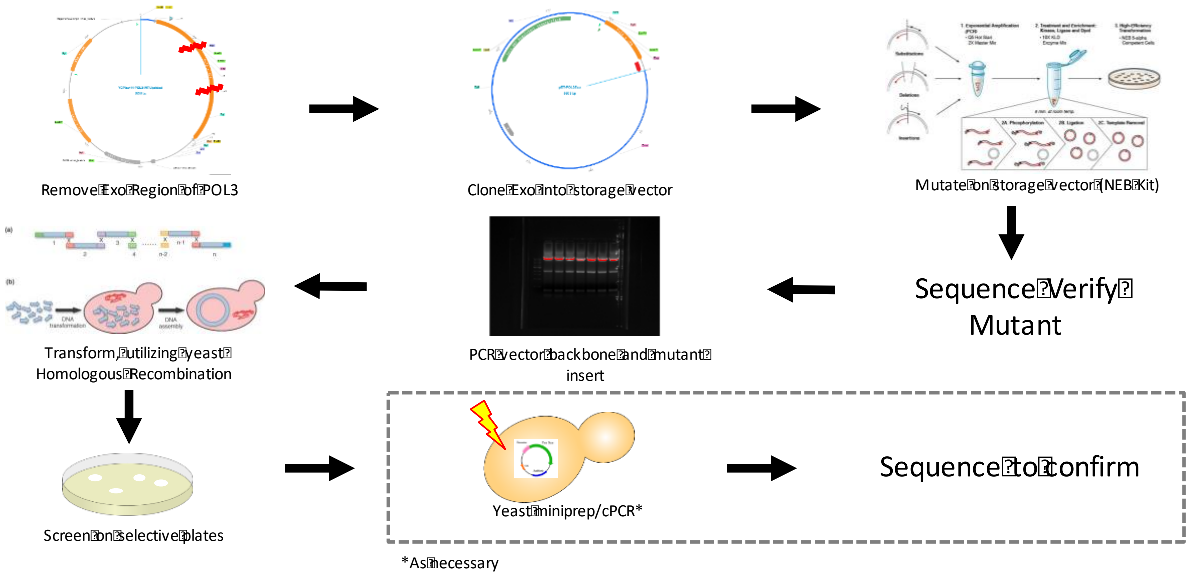


**Figure S26.** Scheme for yeast polymerase screening assay. Variants are created on an excised section of the Pol3 exonuclease domain, verified by sequencing and prepared along with the backbone for yeast transformation. Homologous recombination within yeast allows for creation of the full Pol3 vector. Plating of the yeast reveals whether the variant produces a viable polymerase.

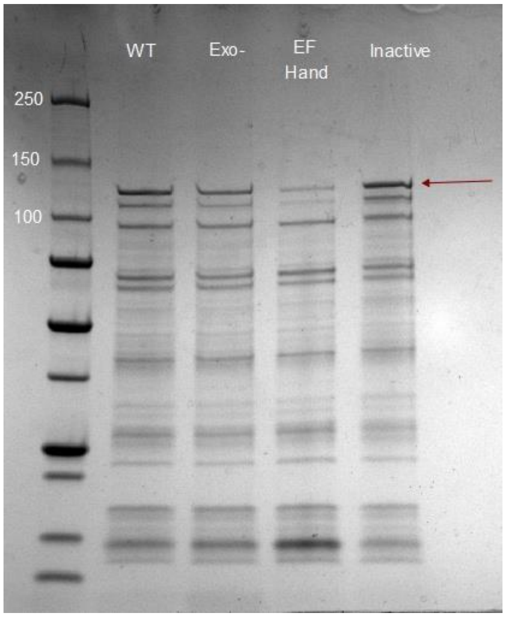


**Figure S27.** *Protein gel for Pol3 NEB T7 Express E. coli expression*. Gel shows strong expression for wild-type (WT) and inactive Pol3 (inactive), modest expression for the variant with the exonuclease deactivated by mutation (Exo$^-$), and poor to no expression for the EF-hand variant (EF-Hand).

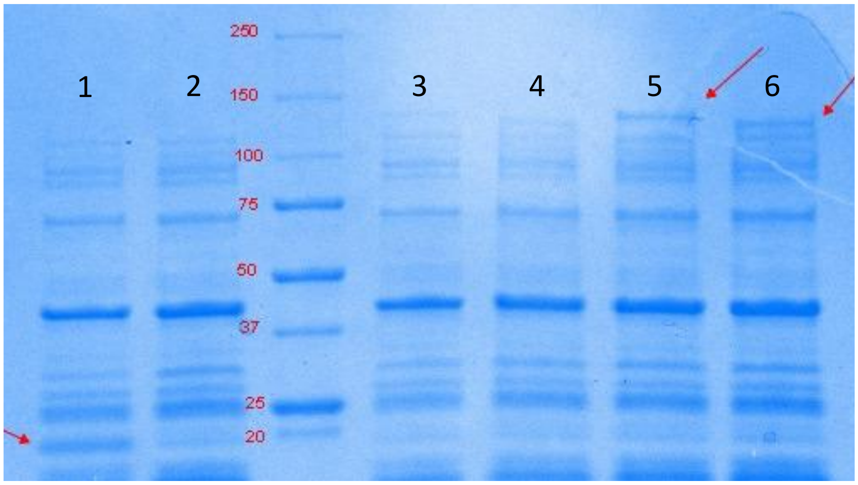


1 – Positive Control DHFR
2 – Negative Control
3 – Yeast codon ΔN-terminal
4 – Yeast codon ΔN,C-terminal
5 – *E. coli* codon ΔN-terminal
6 – *E. coli* codon ΔN,C-terminal

**Expected Bands:**
DHFR - ~20 kDa
ΔN terminus – 144.6 kDa
ΔN,C terminus – 131 kDA

**Figure S28.** *NEB PURExpress expression of Pol3*. Gel shows expression of Pol3 variants with *in vitro* transcription/translation.

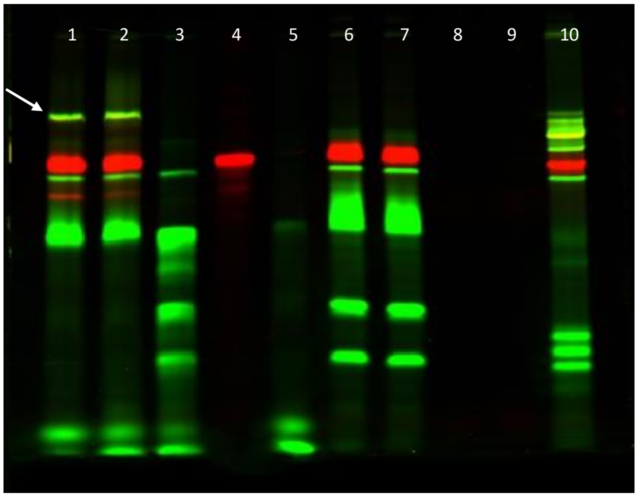


| Samples: |
|---|
| 1 - Test |
| 2 - Test Duplicate |
| 3 - No template |
| 4 - No primer |
| 5 - No dNTPs |
| 6 - No IVTT |
| 7 - Yes POL3 DNA, no IVTT |
| 8 - No primer, template |
| 9 - Only IVTT |
| 10 - Klenow |

Green – Primer
Red – Template

**Figure S29.** *NEB PURExpress Pol3 control gel for fluorescent primer extension assay*. Gel demonstrates the type of extension products observed in this assay and contains controls lacking single elements of the extension assay along with a positive control (lane 10) utilizing the commercial polymerase "Klenow" (the Klenow fragment of DNA polymerase I of *E. coli*).

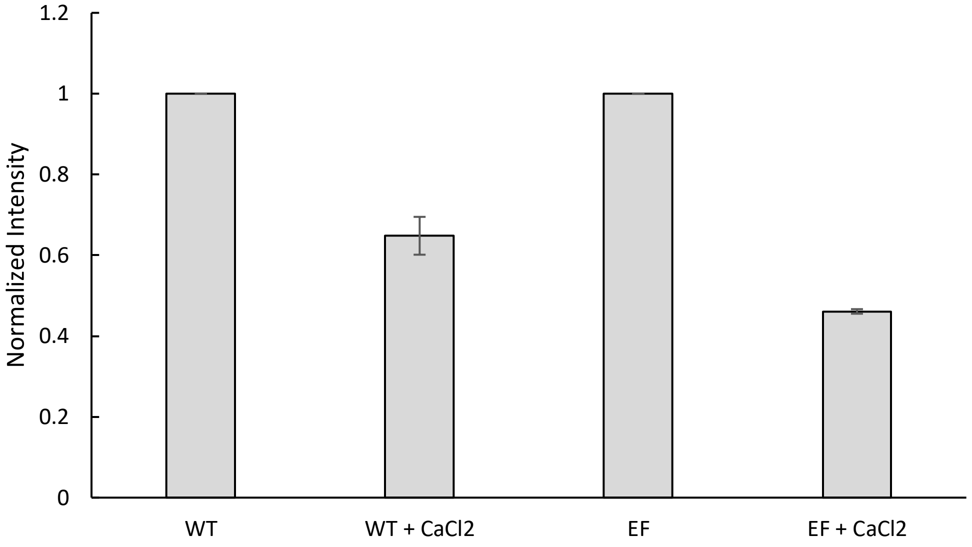


**Figure S30.** *Replicate extension assay data plotted for Pol3 EF-hand grafting approach*. WT = wild-type Pol3, EF = Pol3 EF-hand variant.

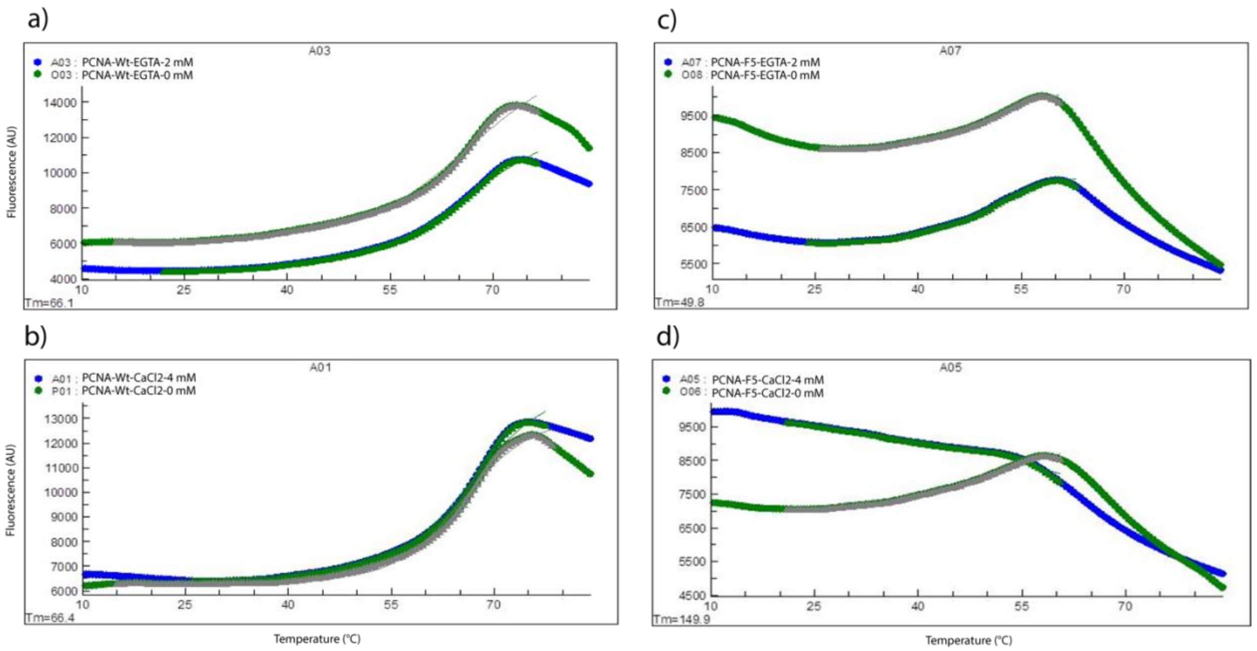


**Figure S31.** *Example melting curves for wild-type PCNA and F5-PCNA-CaM (DKS) in the presence of EGTA and $CaCl_2$.* a) Melting curves for wild-type PCNA with 0 and 2 mM EGTA. b) Melting curves for wild-type PCNA with 0 and 4 mM $CaCl_2$. c) Melting curves for F5-PCNA-CaM (DKS) with 0 and 2 mM EGTA. d) Melting curves for F5-PCNA-CaM with 0 and 4 mM $CaCl_2$. Curves demonstrate a stabilizing effect for both wild-type and F5-PCNA-CaM in the presence of EGTA, possibly due to solvent effect. Wild-type PCNA is minimally impacted by the presence of $CaCl_2$. However, F5 is destabilized in the 4 mM $CaCl_2$ condition. Y-axis is fluorescence (AU) and x-axis is time (min).

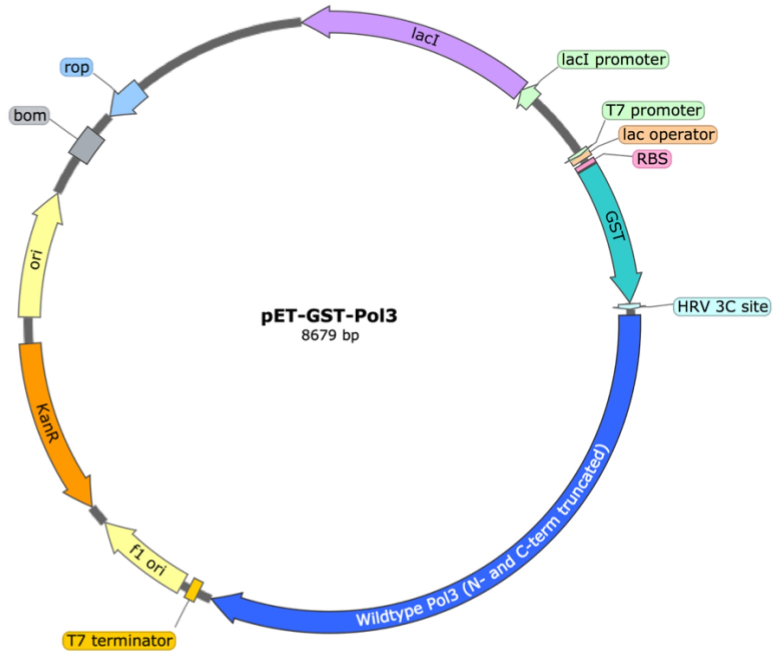


**Figure S32.** *Plasmid map of pET-GST-Pol3 construct*. This vector was used for Gibson assembly of Pol3-CaM fusions. SnapGene® software was used to generate the plasmid map.

### Supplemental Tables

**Table S1.** Multiple Sequence Alignment between *S. cerevisiae* PCNA[a] and Three PCNA Homologs[b].

```
gi|83767836|dbj|BAE57975.1|         MTNILNEEDMLEARLEQASLLKRVVDAIKDLVQDCNFDCNDSGIALQAMDNSHVALVSML
gi|452836388|gb|EME38332.1|         ---------MLEARLEQANLLKKVVDAIKDLVQDCNFDCNDSGIALQAMDNSHVALVSMM
Scer                                ---------MLEAKFEEASLFKRIIDGFKDCVQLVNFQCKEDGIIAQAVDDSRVLLVSLE
gi|363755758|ref|XP_003648094.1|    ---------MLEAKFQQSSFFKKIIDGLKDCVQLVNFNCSEKGIAAQAVDDSRVLLVSLV
                                             ****:::::.::*:::*.:** **  **:*.:.**  **:*:*:* ***:

gi|83767836|dbj|BAE57975.1|         LKAEGFSPYRCDRNIALGINLVSLTKVLRAAQNEDILTLKADDSPDAVNLMFESAETDRL
gi|452836388|gb|EME38332.1|         LKSESFSPFRCDRNIALGINLTSLTKVLRCAQNEDILTMKAEDAPDVVNFTFESAESDRL
Scer                                IGVEAFQEYRCDHPVTLGMDLTSLSKILRCGNNTDTLTLIADNTPDSIILLFEDTKKDRI
gi|363755758|ref|XP_003648094.1|    ITPDAFEEYRSDRAVTLGVDLNSLGKILRCGANEDSLTLVAEDSPDNMLVLFEDTKRDRI
                                     :  :.*. :*.*: ::**::* ** *:**.. * * **: *:::** : . **.:: **:

gi|83767836|dbj|BAE57975.1|         SEYDIKLMDIDQEHLAIPETEYAATVEMPSAEFQRICRDLNALSESVVIEATKEGVKFSC
gi|452836388|gb|EME38332.1|         SEYDIKLMDIDQEHLGIPDTEYAATISLPSSEFQRITRDLSALSESVSIECTKDGVSFKC
Scer                                AEYSLKLMDIDADFLKIEELQYDSTLSLPSSEFSKIVRDLSQLSDSINIMITKETIKFVA
gi|363755758|ref|XP_003648094.1|    SEYSLKLMEIDADFLEIDQIDYDTTIHMPSTEFAKIIRDLNQLSDSLNIVVTKETIKFIS
                                     :**.:***:** :.* * : :* :*: :**:** :* ***. **:*: *  **: :.* .

gi|83767836|dbj|BAE57975.1|         QGDIGSGSVTIRQHTNVDKPEQNVSIALSEPVALTFSLKYLVNFCKATTLSSKVSLCLSQ
gi|452836388|gb|EME38332.1|         NGDIGNGSVTLRSHTNVEKPEQNIEINLSEPVALTFSLKYLMNFCKASGLSGSVKLCLSN
Scer                                DGDIGSGSVIIKPFVDMEHPETSIKLEMDQPVDLTFGAKYLLDIIKGSSLSDRVGIRLSS
gi|363755758|ref|XP_003648094.1|    EGDFGSGSVVVKPRTDVEKPEDSVKVELEKPVDLTFGSKYLLDIIKATSLSTSITIKLSA
                                     :**:*.*** ::  .::::** .:.: :.:** ***. ***::: *.: **  : : **

gi|83767836|dbj|BAE57975.1|         EVPLLVEYGLGS-GHLRFYLAPKIGDEE
gi|452836388|gb|EME38332.1|         EVPLLVEYGLANNSYLRFYLAPKIGDEE
Scer                                EAPALFQFDLK-SGFLQFFLAPKFNDEE
gi|363755758|ref|XP_003648094.1|    ETPALFQFNLDGAGQLQYFLAPKFNEEE
                                    *.* *.:: *    . *:::****: :**
```

[a]The protein sequence of *S. cerevisiae* PCNA was extracted from PDB structure file 1PLR.
[b]Three PCNA homologs from *A. oryzae, D. septosporum NZE10, E. cymbalariae* were aligned with *S. cerevisiae* PCNA.

**Table S2.** Key Features of SCHEMA-Predicted Pol3 Split Sites**.**

| Residue Site | Residue Identity | Residue Number | Domain | Average B-factor[a] |
|---|---|---|---|---|
| 1 | Leucine | 830 | Palm | 27.96 |
| 2 | Valine | 985 | Thumb | 37.12 |
| 3 | Valine | 515 | Exo | 29.75 |
| 4 | Glutamic Acid | 570 | N-term | 33.49 |
| 5 | Aspartic Acid | 321 | Exo | 30.46 |
| 6 | Leucine | 256 | N-term | 27.75 |
| 7 | Tyrosine | 587 | Palm | 21.29 |
| 8 | Tyrosine | 550 | N-term | 26.48 |
| 9 | Serine | 439 | Exo | 30.11 |
| 10 | Glutamine | 579 | Palm | 37.31 |
| 11 | Lysine | 977 | Thumb | 27.55 |
| 12 | Aspartic Acid | 463 | Exo | 26.24 |
| 13 | Lysine | 157 | N-term | 24.49 |
| 14 | Glycine | 782 | Palm | 21.09 |
| 15 | Isoleucine | 662 | Fingers | 23.83 |
| 16 | Leucine | 534 | N-term | 27.96 |
| 17 | Threonine | 606 | Palm | 20.83 |
| 18 | Leucine | 700 | Fingers | 23.74 |
| 19 | Glycine | 589 | Palm | 20.21 |

[a]The average B-factor for each amino acid residue was calculated using PyMOL (PDB ID: 3IAY).

**Table S3.** Table of Yeast Complementation Assay Variants Screened.

| Variant | Sequence | Alive |
|---|---|---|
| Pol3 Wild-type | GFYPFDNVKLAKA | Yes |
| K416E | GFYPFDNVKLA**E**A | Yes |
| K419D | GFYPFDNV**D**LAKA | Yes |
| V420D | GFYPFDN**D**KLAKA | Yes |
| N421D | GFYPFD**D**VKLAKA | Not tested |
| F423D | GFYP**D**ANVKLAKA | Yes |
| P424D | GFY**D**FDNVKLAKA | Yes |
| Y425D | GF**D**PFDNVKLAKA | Yes |
| F426D | G**D**YPFDNVKLAKA | Not Tested |
| G427D, "1/7" | **D**FYPFDNVKLAKA | Yes |
| "2/7", [Y425D] | **D**F**D**PFDNVKLAKA | Yes |
| "3/7", [F426I] | **DID**PFDNVKLAKA | Yes |
| "4/7", [P424G] | **DIDG**FDNVKLAKA | Yes |
| "6/7", [F423N, D422G] | **DIDGNG**NVKLAKA | Yes |
| "7/7" EF Hand Mutant, [K426E] | **DIDGNG**NVKLA**E**A | No |
| "6/7", A415E | **DIDGNG**NVKLAK**E** | Yes |
| "7/7", N421T, K433A | **DIDGNGT**VKLA**E**A | No |
| "7/7", D427A | **AIDGNG**NVKLA**E**A | No |
| "7/7", D425A | **DIAGNG**NVKLA**E**A | No |
| Exo- (D321A, E323A) | | Yes |

**Table S4.** RASPP Script Library Outputs for Five Crossover Points[a].

```
# No bin width specified; using bin width=1.0.
# RASPP took 26.62 secs
# RASPP found 442 results
# RASPP found 16 unique (<E>,<m>) points
# RASPP curve took 23.76 secs
# <E>    <m>      crossover points
19.5000  38.3750  21 49 73 100 124
19.5000  39.3750  21 49 73 100 129
22.2500  40.7500  21 49 73 100 134
23.1250  41.9976  23 59 85 110 138
23.5625  47.5547  23 71 99 124 242
23.5625  48.4280  23 71 99 129 242
24.3125  51.3164  23 71 99 124 232
24.3125  52.0447  23 71 99 129 232
24.6250  52.3999  23 67 100 132 232
25.5000  53.4294  23 67 100 136 232
27.8750  54.4614  23 67 100 136 227
28.8750  55.6035  23 71 99 124 204
29.0625  56.4116  23 67 100 132 204
30.4375  57.7744  23 67 124 151 205
34.3750  58.4150  27 67 117 154 207
39.3125  59.8816  61 99 138 184 232
```

[a]Crossover points reflect residue numbering in PDB structure file 1PLR (*S. cerevisiae* PCNA).

**Table S5.** RASPP Script Library Outputs for Six Crossover Points[a].

```
# No bin width specified; using bin width=1.0.
# RASPP took 24.39 secs
# RASPP found 261 results
# RASPP found 10 unique (<E>,<m>) points
# RASPP curve took 55.28 secs
# <E>    <m>      crossover points
28.5625  48.5903  21 49 73 100 129 242
28.9375  48.9420  21 49 73 100 132 242
31.5000  50.2838  23 53 76 107 136 242
29.3125  51.6830  21 49 73 100 124 232
29.5625  52.8279  21 49 73 100 132 232
30.4375  53.9600  21 49 73 100 136 232
32.8125  59.0157  23 71 99 124 151 204
35.7500  59.8953  23 67 120 151 184 207
37.3125  60.7367  23 67 120 151 204 232
42.7500  61.7132  28 71 115 151 196 232
```

[a]Crossover points reflect residue numbering in PDB structure file 1PLR (*S. cerevisiae* PCNA).

**Table S6.** RASPP Script Library Outputs for Seven Crossover Points[a].

```
# No minimum fragment length specified; using L=4.
# No bin width specified; using bin width=1.0.
# RASPP took 63.61 secs
# RASPP found 1054 results
# RASPP found 26 unique (<E>,<m>) points
# RASPP curve took 1236.26 secs
# <E>    <m>      crossover points
15.0625  38.3750  14 23 71 85 102 114 124
14.8125  39.3750  6 15 23 71 85 117 129
12.0625  41.2492  5 14 19 24 120 129 259
12.9375  41.6229  5 14 19 24 120 132 259
13.2500  43.0953  100 107 114 120 129 136 259
14.5000  43.8271  100 107 114 120 129 138 259
15.5625  45.0051  102 114 120 129 136 254 259
16.3125  45.7112  102 114 120 129 136 253 259
16.9375  47.0737  102 114 120 129 136 251 259
18.0625  48.1628  102 114 120 129 242 251 259
18.4375  48.4557  102 114 120 132 242 251 259
19.3125  49.3798  100 120 129 136 242 251 259
19.9375  51.2932  102 114 120 129 136 232 259
20.5625  51.4972  100 114 120 129 136 232 259
22.4375  52.5544  91 99 120 129 138 232 259
24.2500  53.5806  14 23 71 85 120 136 232
26.1250  54.6131  85 99 120 129 136 204 259
26.1875  55.9905  71 85 120 129 136 204 259
28.1875  56.6409  14 67 120 129 138 202 259
29.0625  57.4140  14 24 71 87 120 138 204
30.5000  58.4397  14 23 67 120 136 151 205
34.1875  59.4444  23 69 87 116 136 199 251
35.0625  60.4525  23 71 99 120 151 184 204
37.5625  61.4747  23 71 117 139 184 205 251
39.5625  62.4294  23 68 120 151 184 204 232
46.2500  63.4037  27 67 100 133 167 199 233
```

[a]Crossover points reflect residue numbering in PDB structure file 1PLR (*S. cerevisiae* PCNA).

**Table S7.** RASPP Script Library Outputs for Eight Crossover Points[a].

```
# No bin width specified; using bin width=1.0.
# RASPP took 20.66 secs
# RASPP found 64 results
# RASPP found 2 unique (<E>,<m>) points
# RASPP curve took 252.55 secs
# <E>    <m>      crossover points
40.5625  61.8715  21 49 73 100 124 151 184 205
42.3750  63.9526  23 71 99 124 151 184 205 232
```

[a]Crossover points reflect residue numbering in PDB structure file 1PLR (*S. cerevisiae* PCNA).

**Table S8.** DNA Sequences used for Fluorescent Primer Extension Activity Assays.

| DNA Name | DNA Sequence (5' - 3') |
|---|---|
| 5' FAM-labeled primer | /56-FAM/A*C*A*C*TGACGACATGGTTCTACAACCGGTATGTACGGCGGTCGGTTATCGTA |
| 5' TAMRA-labeled template | /55-TAMK/CAGGCACGACTAGCAGGCAACGTAGGCACGGAACTCGGAGCACACCTCGAGCACGAACTACGCAACGGCTCGCTACGATAACCGA*C*C*G*C/3ddC/ |

**Table S9.** Selected Pol3 Homologs from NCBI Protein-Protein BLAST.

| DNA Polymerase | Organism | % Protein Identity | Accession Number |
|---|---|---|---|
| DNA polymerase delta catalytic subunit | *Kluyveromyces marxianus* | 78.38% | XP_022674962.1 |
| DNA polymerase delta catalytic subunit | *Wickerhamomyces ciferrii* | 65.14% | XP_011276882.1 |
| DNA polymerase delta catalytic subunit | *Spathaspora passalidarum* | 64.80% | XP_007377768.1 |
| DNA polymerase delta catalytic subunit | *Scheffersomyces stipitis* | 64.72% | XP_001386065.2 |
| DNA polymerase delta catalytic subunit | *Candida albicans* | 64.57% | CAA61282.1 |

**Table S10.** Multiple Sequence Alignment between *S. cerevisiae* Pol3[a] and Five Pol3 Homologs[b].

```
Pol3                                  SSFERKKLPTDFDPSLYDISFQQIDAEQSVLNGIKDENTSTVVRFFGVTSEGHSVLCNVT
gi|574141310|dbj|BAO39104.1|          NLWCRKPIPDDFDPNLSDISFQQLDAEQAILPGQYDSNTNVVTRFFGVTDHGNSILCNVT
gi|754418003|ref|XP_011276882.1|      QIWERPTLPVDWKPETHDITFQQLDAEEALT------EGSSAVRFFGVTQEGYSVLCNVT
gi|951001|emb|CAA61282.1|             QTWDRPPLPSSF----EDISFQQLDAEEYHD------RGNTYARFFGITQEGHSVLCNVT
gi|598073879|ref|XP_007377768.1|      QNWSRPELSADFNSELEDVSFQQLDAEEFNE------RDNTFARFFGITEAGHSVLCNVV
gi|150866455|ref|XP_001386065.2|      QKWGRPPLPETFDPTIDDVSFQQLDAEEFQI------GDYTIARYFGITEEGHSVLCNVT
                                      . : *  :   :     *::***:***:              .*:**:*. * *:****.

Pol3                                  GFKNYLYVPAPNSSDANDQEQINKFVHYLNETFDHAIDSIEVVSKQSIWGYSGDTKLPFW
gi|574141310|dbj|BAO39104.1|          GFKHYLYVPAPLGFQQ---TDVATLVQYLNEHFENNVDSIKIVSKQSIWGFSGDAKIPFL
gi|754418003|ref|XP_011276882.1|      GFLHYFYVPAPLNFRKE---HLLSFKNYLQQSFE-GVLDIELCFKESIWGFNGNQKVCFL
gi|951001|emb|CAA61282.1|             GFIHYFYCPVPKGFEE----NLTEFTNYLKATFD-GIERVEITSKESIWGYSNNIKTPFF
gi|598073879|ref|XP_007377768.1|      GFIHYFYVPVPKGFVKD--QHLQQFTNYLHANYE-GIDKVEITLKETIWGYNRNIKTPFF
gi|150866455|ref|XP_001386065.2|      GFVHYFYVPVPKGFYKD--QHLDQFSSYLRANYE-GVENIELALKESIWGYNKSVKTPFF
                                      ** .*:* *.*         .:  :  **.  ::  :  :::  *::***:. . *  *

Pol3                                  KIYVTYPHMVNKLRTAFERGHLSF-NSWFSN--GTTTYDNIAYTLRLMVDCGIVGMSWIT
gi|574141310|dbj|BAO39104.1|          QIFVKNPNMLNKIRTGFEKGYIQPNDKWFVG--GCTTYDNIAYTLRLMIDCGIVGMSWIT
gi|754418003|ref|XP_011276882.1|      KIIGVNAKDIPKIRSGFEKGMVSWNGMFNSNGDGTMTYDNIQYLLRLMIDCKITGMSWIT
gi|951001|emb|CAA61282.1|             KIFAKN--NISKIRSAFQNGQVPNI-------DPCITYDNINYLLRLMIDCKITGMSWIT
gi|598073879|ref|XP_007377768.1|      KVFVNNTRNITKLRTAFERGDIRFENLFS----ETVSYDNINYLMRLMIDCKITGMSWIT
gi|150866455|ref|XP_001386065.2|      KFFINNTKNITKLRSAFERGEIRFENLFP---PQNVSYDNINYLLRLMIDCKVTGMSWVT
                                      :.       : *:*:.*:.* :              :**** * :***:** :.****:*

Pol3                                  LPKGKYSMIEPNNRVSSCQLEVSINYRNLIAHPAEGDWSHTAPLRIMSFDIECAGRIGVF
gi|574141310|dbj|BAO39104.1|          LPASKYIMVPQDQRVSTCQFEVNINYKDLISHPAEGDWSHNAPLRIMSFDIECAGRPGIF
gi|754418003|ref|XP_011276882.1|      LPKGTYTPVETGLKTSRCQLEVNIDYKSLISHPPEGEWLKMAPLRILSFDIECAGRKGIF
gi|951001|emb|CAA61282.1|             LPRDKYKIVN--NKISTCQIECSIDYRDLISHPPEGEWLKMAPLRILSFDIECAGRKGVF
gi|598073879|ref|XP_007377768.1|      LPKSKYKMVPDNLKISSCQIECSIDYRDLITHPSEGEWLKMAPLRILSFDIECAGRKGIF
gi|150866455|ref|XP_001386065.2|      LPKGKFSLVHSNDKVSTCQIECSINYKDLISHPSEGEWLKMAPLRILSFDIECAGRKGIF
                                      **  .:  :    : * **:* .*:*:.**:** **:* : *****:********* *:*

Pol3                                  PEPEYDPVIQIANVVSIAGAKKPFIRNVFTLNTCSPITGSMIFSHATEEEMLSNWRNFII
gi|574141310|dbj|BAO39104.1|          PEPEHDAVIQIANVVSIAGAPKPFIRNVFTVNTCSPITGSQIFEHEQESDMLKHWRDFIV
gi|754418003|ref|XP_011276882.1|      PEAEHDSIIQIANVVSRSGESRPFVRNVFTVNSCSPITGSEIFAHEDERDMLLEWKDFVN
gi|951001|emb|CAA61282.1|             PEAEHDPVIQIANVVQKSGESKPFVRNVFTVNTCSSIIGSQIFEHQREEDMLMHWKEFIT
gi|598073879|ref|XP_007377768.1|      PEAQHDPVIQIANVVSKSGESKPFVRNVFTVNTCSPIIGSQIFEHAKEEEMLMHWRQFIT
gi|150866455|ref|XP_001386065.2|      PEAEHDPVIQIANVVSKHGESRPFVRNVFTVNTCSPIIGSQIFEHEKEEDMLMNWKDFVV
                                      ** ::* :*******.  *  :**:*****:*:** * ** ** *  * :** .*::*:
```

```
Pol3                                KVDPDVIIGYNTTNFDIPYLLNRAKALKVNDFPYFGRLKTVKQEIKESVFSSKAYGTRET
gi|574141310|dbj|BAO39104.1|        EVDPDVIIGYNTTNFDFPYLLDRAAALGVHSFPYFGRLSNVKQEIKSSTFSSKAYGTRES
gi|754418003|ref|XP_011276882.1|    KVDPDVIIGYNTTNFDLPYLLDRAKALGLRDFPYFGRLINIKQEAKDSVFSSKAYGTRES
gi|951001|emb|CAA61282.1|           KVDPDVIIGYNTANFDIPYVLNRAKALGLNDFPFFGRLKRVKQEIKDAVFSSRAYGTREN
gi|598073879|ref|XP_007377768.1|    EVDPDVIIGYNTSNFDIPYLLDRAKALGLKDFPFFGRLKRIKQEVKDAVFSSRAYGTREN
gi|150866455|ref|XP_001386065.2|    SVDADVIIGYNTANFDIPYLLDRAKALGLRNFPFFSRLKNSKQEAKDSVFSSRAYGTREN
                                    .** ********:***:**:*:** ** :..**:*.**   *** *.:.***:******.

Pol3                                KNVNIDGRLQLDLLQFIQREYKLRSYTLNAVSAHFLGEQ------SIISDLQNGDSETRR
gi|574141310|dbj|BAO39104.1|        KNVNIDGRLQLDLLQFVQREYKLRSYTLNSVSAHFLGEQKEDVHHSIITSLQNGDSETRR
gi|754418003|ref|XP_011276882.1|    KSINIDGRMQLDLLQFIQREYKLRSYTLNSVSAHFLNEQKEEVQHSIITDLQNGNSETRR
gi|951001|emb|CAA61282.1|           KVVNIDGRMQLDLLQFIQREYKLRSYTLNSVSAHFLGEQKEDVQHSIITDLQNGTKETRR
gi|598073879|ref|XP_007377768.1|    KVVNIDGRMQLDLLQFVQREYKLRSYTLNSVSAHFLGEQKEDVQHSIITDLQNGTKETRR
gi|150866455|ref|XP_001386065.2|    KVVNIDGRMQLDLLQFIQREYKLRSYTLNSVSAHFLGEQKEDVQHSIITDLQNGTSETRR
                                    * :*****:*******:************:****** **      ***:.**** .****

Pol3                                RLAVYCLKDAYLPLRLMEKLMALVNYTEMARVTGVPFSYLLARGQQIKVVSQLFRKCLEI
gi|574141310|dbj|BAO39104.1|        RLAVYCLKDAYLPLRLMEKLMALVNYTEMSRVTGVPFMYLLSRGQQIKVISQLFRKCLQI
gi|754418003|ref|XP_011276882.1|    RLAVYCLKDAYLPMRLLEKLMCLVNYTEMARVTGVPFSFLLTRGQQIKVISQLFRKCLEI
gi|951001|emb|CAA61282.1|           RLAVYCLKDAFLPLRLLDKLMCLVNYTEMARVTGVPFSYLLSRGQQIKVISQLFRKCLQE
gi|598073879|ref|XP_007377768.1|    RLAVYCLKDAFLPLRLLDKLMCLVNYTEMARVTGVPFSYLLSRGQQIKVISQLFRKCLEE
gi|150866455|ref|XP_001386065.2|    RLAVYCLKDAYLPLRLLDKLMCLVNYTEMARVTGVPFSYLLARGQQIKVISQLFRKCLEE
                                    **********:**:**::***.*******:******* :**:*******:********:

Pol3                                DTVIPNMQSQAS-DDQYEGATVIEPIRGYYDVPIATLDFNSLYPSIMMAHNLCYTTLCNK
gi|574141310|dbj|BAO39104.1|        ETVIPNMGSQGS-EEQYEGATVIEPKRGYYDVPIATLDFSSLYPSIMMAHNLCYTTLCDR
gi|754418003|ref|XP_011276882.1|    DLVVPNLQQGGSSDVQYEGATVIEPHRGYYDVPIATLDFSSLYPSIMMAHNLCYTTLLNK
gi|951001|emb|CAA61282.1|           DIVIPNLKSEGS-NEEYEGATVIEPERGYYDVPIATLDFSSLYPSIMMAHNLCYTTLLNK
gi|598073879|ref|XP_007377768.1|    DIIIPNMKSEGT-NEEYEGATVIEPIRGYYDVPIATLDFSSLYPSIMMAHNLCYTTLLSK
gi|150866455|ref|XP_001386065.2|    DIVIPNMKSEGT-NEEYEGATVIEPSRGYYDVPIATLDFSSLYPSIMMAHNLCYTTLLNI
                                    : ::**: . .: : :********* *************.***************** .

Pol3                                ATVERLNLKIDEDYVITPNGDYFVTTKRRRGILPIILDELISARKRAKKDLRDEKDPFKR
gi|574141310|dbj|BAO39104.1|        STVKRLNLKENDDYIVTPNNDIFVTQKVRKGVLPEILDELLSARKKAKRDLKNETDPFKR
gi|754418003|ref|XP_011276882.1|    PTIERLKLEKDKDYTVSPSGDFFVKTSQRKGILPIILEELLSARKKAKKDMKTETDPFKR
gi|951001|emb|CAA61282.1|           NSIKAFGLT-EDDYTKTPNGDYFVHSNLRKGILPTILDELLTARKKAKADLKKETDPFKK
gi|598073879|ref|XP_007377768.1|    SAIQAYGLT-EEDYTRTPNGDFFVKSHKRQGILPTILNELLTARKKAKADLKKETDPFKK
gi|150866455|ref|XP_001386065.2|    QTIKAFNLT-EDDYTRTPNGDYFVKDHKKQGILPTILNELLTARKKAKADLKKETDPFKR
                                     :::   *  :.**  :*. * **    ::*:** **:**::***:** *:: *.****:

Pol3                                DVLNGRQLALKISANSVYGFTGATVGKLPCLAISSSVTAYGRTMILKTKTAVQEKYCIKN
gi|574141310|dbj|BAO39104.1|        DVLNGRQLALKISANSVYGFTGATVGKLPCLAISSSVTAFGRTMIEATKNAVEEKYSIKN
gi|754418003|ref|XP_011276882.1|    DVLNGRQLALKISANSVYGFTGATIGKLPCLAISSSVTAFGRGMIEKTKQEVQDKYTIAN
gi|951001|emb|CAA61282.1|           DVLNGRQLALKISANSVYGFTGATVGKLPCLAISSSVTAFGREMIEKTKNEVQEYYSKKN
gi|598073879|ref|XP_007377768.1|    DVLNGRQLALKISANSVYGFTGATIGKLPCLAISSSVTAFGREMIEKTKNEVQEYYSKKN
gi|150866455|ref|XP_001386065.2|    DVLNGRQLALKISANSVYGFTGATIGKLPCLAISSSVTAFGREMIEKTKNEVQEFYSKKN
                                    ************************:**************:** **  **  *:: *   *

Pol3                                GYKHDAVVVYGDTDSVMVKFGTTDLKEAMDLGTEAAKYVSTLFKHPINLEFEKAYFPYLL
gi|574141310|dbj|BAO39104.1|        GATHDAVVVYGDTDSVMVKFGTTNLEESMKLGAEAADYVSGLFKNPIKLEFEKVYFPYLL
gi|754418003|ref|XP_011276882.1|    GYDYDSEVIYGDTDSVMVKFGYKDLETCMKLGEEAADFVSTKFLNPIKLEFEKVYFPYLL
gi|951001|emb|CAA61282.1|           GHPYDAKVIYGDTDSVMVKFGYQDLETCMKLGEEAANYVSTKFKNPIKLEFEKVYFPYLL
gi|598073879|ref|XP_007377768.1|    GHPFDAQVIYGDTDSVMVKFGYQDLETCMKLGEEAANYVSTKFKSPIKLEFEKVYYPYLL
gi|150866455|ref|XP_001386065.2|    GYPYDAQVIYGDTDSVMVKFGYQDLETCMKLGEEAANYVSTKFKKPIKLEFEKVYFPYLL
                                    *  .*: *:************  :*: .*.** ***.:**  *  **:*****.*:****

Pol3                                INKKRYAGLFWTNPDKFDKLDQKGLASVRRDSCSLVSIVMNKVLKKILIERNVDGALAFV
gi|574141310|dbj|BAO39104.1|        INKKRYAGLYWTNPEKYDKLDQKGLASVRRDSCPLVSIVMNKVLRKILIDRNVEGALQFI
gi|754418003|ref|XP_011276882.1|    INKKRYAGLYWTNTEKFDKMDTKGIETVRRDNCRLVSNVITKILELILEKRDVKSAENFV
gi|951001|emb|CAA61282.1|           INKKRYAGLYWTRPEKFDKMDTKGIETVRRDNCQLVQNVITKVLEFILEERDVPKAQRFV
gi|598073879|ref|XP_007377768.1|    INKKRYAGLYWTRPDKFDKMDTKGIETVRRDNCRLVQNVITKVLQFILEERDVEKAQRFV
gi|150866455|ref|XP_001386065.2|    INKKRYAGLYWTRPDKFDKMDTKGIETVRRDNCRLVQNVITKVLEFILEDRDVEKAQRFV
                                    *********:**. :*:**:* **: :****.* **. *:.*:*. ** .*:*  *  *:

Pol3                                RETINDILHNRVDISKLIISKTLA-PNYTNPQPHAVLAERMKRRE-GVGPNVGDRVDYVI
gi|574141310|dbj|BAO39104.1|        RETIDDILQNRCDISKLIISKTLA-PNYTNPQPHAVLAERMKKRD-GVGPNVGDRVDYVI
gi|754418003|ref|XP_011276882.1|    KQTIADLLQDRVDMQQLVITKALSRSDYANKQPHVELADRMRKRDAGSAPTIGDRVAYVI
gi|951001|emb|CAA61282.1|           KQTIADLLQNRIDLSQLVITKAYSKHDYSAKQAHVELAERMRKRDPGSAPTLGDRVAYVI
gi|598073879|ref|XP_007377768.1|    KQTIADLLQNRVDLSQLVITKAYAKHDYSAKQAHVELAERMKKRDPGSAPTLGDRVAYVI
gi|150866455|ref|XP_001386065.2|    KQTIADLLQNRVDLSQLVITKAYAKHDYAAKQAHVELAERMKKRDAGSAPTLGDRVAYVI
                                    ::** *:*::* *:.:*:*:*: :  :*:  * *. **:**::*: * .*.:**** ***
```

```
Pol3                                 IG--GNDKLYNRAEDPLFVLENNIQVDSRYYLTNQLQNPIISIVAPIIGDKQANGMFVV
gi|574141310|dbj|BAO39104.1|         IG--GNDKLFNRAEDPLYVLEQNIQLDSMYYLTNQLQNPIISIIAPIIGEKQANSMFIV
gi|754418003|ref|XP_011276882.1|     IK-TAGNKSYEKSEDPLYVLEHNLPIDSKYYLENQLANPLTRIFEPILGEKKAKELLTG
gi|951001|emb|CAA61282.1|            IK-TGGDKNYEKSEDPLYVLENSLPIDVKYYLDQQLTKPLERIFIPILGETKTKELLTG
gi|598073879|ref|XP_007377768.1|     V-NTGSDKNYEKSEDPLYVLENSLPIDVKYYLDQQLTKPLERIFNPILGDGKTKELLAG
gi|150866455|ref|XP_001386065.2|     IKGSGSDKNYEKSEDPLYVLENSLPIDVKYYLDNQLTKPLERIFNPILGDAKTKELLAG
                                     :   . :* ::::****:***..: :*  *** :** :*:  *. **:*: ::: ::
```

[a]The protein sequence of *S. cerevisiae* Pol3 was extracted from the PDB structure file 3IAY (top sequence).

[b]Five Pol3 homologs from *K. marxianus*, *W. ciferrii*, *C. albicans*, *S. passalidarum*, and *S. stipitis*, (ordered consecutively from top to bottom) were aligned with *S. cerevisiae* Pol3.

**Table S11.** RASPP Script Library Outputs for Four Crossover Points[a].

### No bin width specified; using bin width=1.0.
### RASPP took 8850.04 secs
### RASPP found 13530 results
### RASPP found 35 unique (<E>,<m>) points
### RASPP curve took 642.89 secs

| # <E> | <m> | crossover points |
|---|---|---|
| 0.2500 | 47.9444 | 730 885 892 896 |
| 0.3889 | 91.1111 | 470 487 885 892 |
| 0.6944 | 92.0000 | 459 480 487 885 |
| 2.0278 | 95.7209 | 415 885 892 896 |
| 2.3333 | 99.3134 | 369 415 885 893 |
| 3.0833 | 99.9932 | 363 415 480 730 |
| 3.3889 | 101.5117 | 345 360 397 885 |
| 3.3889 | 102.1043 | 345 390 415 885 |
| 3.5556 | 103.2591 | 340 390 431 885 |
| 3.8611 | 104.4959 | 340 415 487 730 |
| 4.1667 | 104.9640 | 339 415 487 730 |
| 5.0000 | 111.2491 | 227 415 885 893 |
| 5.0000 | 112.1232 | 6 227 415 885 |
| 5.0556 | 113.4556 | 6 227 450 885 |
| 5.0556 | 113.9756 | 6 227 470 885 |
| 5.0556 | 114.9516 | 227 415 487 885 |
| 5.2500 | 120.4576 | 227 415 730 885 |
| 5.6667 | 120.9511 | 227 369 488 730 |
| 5.9722 | 122.2047 | 211 450 730 885 |
| 6.4167 | 123.0361 | 205 450 730 885 |
| 8.0278 | 123.9574 | 202 369 489 730 |
| 7.7222 | 125.2052 | 19 227 415 730 |
| 7.3333 | 126.8467 | 162 470 487 730 |
| 7.3333 | 127.4554 | 6 162 470 730 |
| 7.3333 | 128.2434 | 162 470 730 885 |
| 8.0833 | 129.0697 | 153 470 730 885 |
| 8.3889 | 130.6640 | 19 162 470 730 |
| 8.3889 | 130.9954 | 24 162 470 730 |
| 12.8611 | 131.9722 | 153 284 479 730 |
| 16.3889 | 133.3374 | 148 284 487 730 |
| 10.7500 | 134.8889 | 63 162 470 730 |
| 11.0556 | 134.9825 | 63 162 459 730 |
| 17.3889 | 135.9861 | 78 162 459 730 |
| 26.1111 | 137.0552 | 126 227 490 752 |
| 39.0833 | 137.9492 | 117 271 541 763 |

[a]Crossover points reflect residue numbering in PDB structure file 3IAY (*S. cerevisiae* Pol3).

**Table S12.** RASPP Script Library Outputs for Five Crossover Points[a].

```
# No bin width specified; using bin width=1.0.
# RASPP took 9155.05 secs
# RASPP found 11189 results
# RASPP found 37 unique (<E>,<m>) points
# RASPP curve took 3537.88 secs
```

| # <E> | <m> | crossover points |
|---|---|---|
| 0.6389 | 91.1111 | 470 487 730 885 894 |
| 0.8889 | 92.5556 | 450 479 487 885 893 |
| 2.0833 | 95.7222 | 415 450 479 487 885 |
| 2.3889 | 99.6039 | 369 415 450 480 885 |
| 3.3889 | 102.1823 | 345 360 390 415 885 |
| 3.5000 | 103.1334 | 345 400 450 488 885 |
| 3.6111 | 104.1715 | 340 390 431 487 885 |
| 4.8056 | 105.3138 | 336 377 431 487 730 |
| 5.3889 | 106.8991 | 330 369 434 487 730 |
| 5.0556 | 111.5842 | 227 415 450 479 487 |
| 5.0556 | 114.0728 | 227 415 450 480 885 |
| 5.0556 | 115.0386 | 227 415 450 487 885 |
| 5.1111 | 115.2258 | 227 415 450 488 885 |
| 5.2500 | 120.5130 | 227 415 730 885 894 |
| 5.3056 | 121.4068 | 227 415 479 730 885 |
| 5.8056 | 122.4267 | 7 227 415 730 885 |
| 6.3333 | 123.4521 | 207 450 487 730 885 |
| 7.0000 | 124.2147 | 202 450 487 730 885 |
| 10.0556 | 125.1277 | 202 286 369 489 730 |
| 7.7778 | 127.0098 | 19 227 415 487 730 |
| 7.9444 | 127.5030 | 24 227 415 487 730 |
| 7.3333 | 128.8433 | 162 470 487 730 885 |
| 7.6944 | 129.2121 | 156 470 487 730 885 |
| 8.5278 | 131.0988 | 19 169 450 479 730 |
| 8.3889 | 131.3188 | 19 162 470 487 730 |
| 8.3889 | 132.8623 | 19 162 470 730 885 |
| 8.5278 | 133.1612 | 24 169 450 730 885 |
| 9.2778 | 134.1945 | 24 162 470 730 877 |
| 15.6389 | 135.1393 | 152 247 415 562 730 |
| 18.9444 | 136.2306 | 148 245 415 562 730 |
| 12.7222 | 137.5248 | 63 162 415 490 730 |
| 12.7500 | 138.2965 | 63 162 369 487 730 |
| 13.8056 | 139.1411 | 63 162 345 487 730 |
| 15.8056 | 140.1904 | 64 162 340 490 745 |
| 17.5833 | 141.4355 | 70 152 247 487 742 |
| 21.6389 | 142.3056 | 63 162 345 562 770 |
| 26.5000 | 143.1611 | 74 169 345 575 789 |

[a]Crossover points reflect residue numbering in PDB structure file 3IAY (*S. cerevisiae* Pol3).

**Table S13.** RASPP Script Library Outputs for Seven Crossover Points[a].

```
# No bin width specified; using bin width=1.0.
# RASPP took 6671.32 secs
# RASPP found 5343 results
# RASPP found 36 unique (<E>,<m>) points
# RASPP curve took 117482.87 secs
```

| # <E> | <m> | crossover points |
|---|---|---|
| 5.8056 | 91.1111 | 470 506 562 600 682 730 877 |
| 6.3056 | 92.5556 | 450 506 562 600 682 730 877 |
| 6.7222 | 95.7222 | 415 482 538 562 600 682 730 |
| 6.9167 | 99.8310 | 369 450 520 562 600 682 730 |
| 7.9722 | 103.6451 | 345 415 487 562 600 682 730 |
| 8.1389 | 104.8574 | 340 415 487 562 600 682 730 |
| 7.3611 | 105.3273 | 339 400 450 491 682 730 877 |
| 7.8333 | 123.1341 | 227 271 369 434 487 730 877 |
| 16.0556 | 124.8283 | 227 369 506 640 714 789 849 |
| 8.6389 | 126.1033 | 205 247 369 434 487 730 877 |
| 8.8333 | 126.4227 | 203 245 369 434 487 730 877 |
| 11.7500 | 127.3910 | 202 260 345 415 489 730 873 |
| 13.4444 | 128.2828 | 203 273 369 487 730 789 858 |
| 8.9722 | 129.6788 | 19 227 369 434 487 730 877 |
| 9.1389 | 130.2191 | 24 227 369 434 487 730 877 |
| 10.4444 | 131.4049 | 19 211 369 437 489 730 877 |
| 11.2500 | 132.1458 | 28 211 363 437 490 730 877 |
| 11.5556 | 133.4757 | 24 202 247 415 487 730 877 |
| 14.4167 | 134.8605 | 153 211 271 340 415 490 730 |
| 13.3333 | 135.4109 | 29 169 450 506 562 640 730 |
| 10.9167 | 136.7914 | 19 169 369 434 487 730 877 |
| 10.9167 | 137.1882 | 24 169 369 434 487 730 877 |
| 11.8333 | 138.1474 | 28 156 369 437 489 730 877 |
| 13.0833 | 139.2109 | 24 153 345 415 487 730 877 |
| 13.6389 | 141.9089 | 63 169 369 434 487 730 877 |
| 13.8333 | 142.2297 | 63 162 369 437 489 730 877 |
| 14.8056 | 143.2796 | 63 162 345 415 489 730 877 |
| 14.9444 | 144.3205 | 63 162 211 450 491 730 877 |
| 16.3611 | 145.1708 | 63 162 214 450 506 735 874 |
| 17.1389 | 146.2211 | 63 162 227 415 490 730 863 |
| 18.7500 | 147.3547 | 63 156 227 415 496 730 863 |
| 19.5000 | 148.1860 | 63 153 227 415 506 730 862 |
| 21.4167 | 149.1879 | 64 152 227 369 489 682 789 |
| 26.1389 | 150.1323 | 70 151 243 345 493 682 792 |
| 30.4167 | 151.1669 | 73 150 247 369 557 693 800 |
| 33.1667 | 152.1898 | 63 138 227 377 562 730 828 |

[a]Crossover points reflect residue numbering in PDB structure file 3IAY (*S. cerevisiae* Pol3).

**Table S14.** Most Frequently-Occurring Crossover Residues from Three Independent SCHEMA Runs.

| # Crossover Points | 3IAY Residue Number[a] | Pol3 Residue Number[b] | Frequency[c] |
|---|---|---|---|
| 4 Crossover Points | 730 | 830 | 21 |
| | 885 | 985 | 18 |
| | 415 | 515 | 11 |
| | 470 | 570 | 9 |
| | 227 | 321 | 9 |
| | 162 | 256 | 8 |
| | 487 | 587 | 7 |
| 5 Crossover Points | 730 | 830 | 23 |
| | 885 | 985 | 19 |
| | 487 | 587 | 18 |
| | 415 | 515 | 15 |
| | 450 | 550 | 12 |
| | 227 | 321 | 9 |
| | 162 | 256 | 9 |
| | 470 | 570 | 6 |
| | 345 | 439 | 5 |
| | 479 | 579 | 5 |
| 7 Crossover Points | 730 | 830 | 31 |
| | 877 | 977 | 19 |
| | 369 | 463 | 16 |
| | 487 | 587 | 13 |
| | 415 | 515 | 11 |
| | 227 | 321 | 9 |
| | 63 | 157 | 9 |
| | 682 | 782 | 9 |
| | 562 | 662 | 8 |
| | 434 | 534 | 8 |
| | 450 | 550 | 6 |
| | 506 | 606 | 6 |
| | 600 | 700 | 6 |
| | 489 | 589 | 6 |
| | 345 | 439 | 5 |
| | 162 | 256 | 5 |

[a]Residue number from PDB structure file 3IAY.
[b]Equivalent residue number in wild-type Pol3 sequence (this is the numbering used in this study).
[c]Only crossover residues appearing ≥ 5 times in a given run were considered.

**Table S15.** DNA Sequences of GSGGG-CaM-GSGGG and $Pol3_{67–985}$.

| DNA Name | DNA Sequence (5' - 3') |
|---|---|
| **GSGGG-CaM-GSGGG** | GGAAGTGGTGGAGGTATGGCGGATCAGCTGACCGAAGAACAGATTGCGGAATTTAAAGAAGCGTTTAGCCTGTTTGATAAAGATGGCGATGGC<br>ACCATTACCACCAAAGAACTGGGCACCGTGATGCGCAGCCTGGGCCAGAACCCGACCGAAGCGGAACTGCAGGATATGATTAACGAAGTGGA<br>TGCGGATGGCAACGGCACCATTGATTTTCCGGAATTTCTGACCATGATGGCGCGCAAAATGAAAGATACCGATAGCGAAGAAGAAATTCGCGA<br>AGCGTTTCGCGTGTTTGATAAAGATGGCAACGGCTATATTAGCGCGGCGGAACTGCGCCATGTGATGACCAACCTGGGCGAAAAACTGACCGA<br>TGAAGAAGTGGATGAAATGATTCGCGAAGCGGATATTGATGGCGATGGCCAGGTGAACTATGAAGAATTTGTGCAGATGATGACCGCGAAAGG<br>ATCCGGTGGAGGT |
| **N- and C-term truncated Pol3 (residues 67-985)** | ATGGGCACCCAGCTGGAAAGCACCTTTGAACAAGAACTGAGCCAGATGGAACATGATATGGCCGATCAAGAAGAACACGATCTGAGCAGCTTT<br>GAACGTAAAAAACTGCCGACCGATTTTGATCCGAGCCTGTATGATATTAGCTTTCAGCAGATTGATGCAGAACAGAGCGTTCTGAATGGCATCAA<br>AGATGAAAATACCAGCACCGTGGTTCGTTTTTTTGGTGTTACCAGCGAAGGTCATAGCGTTCTGTGTAATGTTACCGGCTTTAAAAACTATCTGTA<br>TGTGCCTGCACCGAATAGCAGTGATGCAAATGATCAAGAGCAGATTAACAAATTCGTGCATTACCTGAACGAAACCTTTGATCATGCCATTGATA<br>GCATTGAAGTTGTGAGCAAACAGAGCATTTGGGGTTATAGCGGTGATACCAAACTGCCGTTTTGGAAAATCTATGTTACCTATCCGCACATGGTG<br>AATAAACTGCGTACCGCATTTGAACGTGGTCATCTGAGCTTTAATAGCTGGTTTAGCAATGGCACCACCACCTATGATAATATTGCATATACCCTG<br>CGTCTGATGGTTGATTGTGGTATTGTTGGTATGAGCTGGATTACCCTGCCGAAAGGTAAATATAGCATGATTGAACCGAATAATCGTGTTAGCAG<br>CTGTCAGCTGGAAGTGAGCATTAACTATCGTAATCTGATTGCACATCCGGCAGAAGGTGATTGGAGCCATACCGCACCGCTGCGTATTATGAGC<br>TTTGATATTGAATGTGCAGGTCGTATTGGTGTTTTTCCGGAACCGGAATATGATCCGGTTATTCAGATTGCAAATGTTGTTAGCATTGCCGGTGCA<br>AAAAAACCGTTTATTCGTAATGTGTTTACCCTGAACACCTGTAGCCCGATTACCGGTAGCATGATTTTTAGCCATGCAACCGAAGAAGAAATGCT<br>GAGCAATTGGCGCAACTTTATCATTAAAGTTGATCCGGATGTGATTATTGGCTACAACACCACCAATTTCGATATTCCGTATCTGCTGAATCGTGC<br>AAAAGCCCTGAAAGTTAATGATTTTCCGTATTTCGGTCGCCTGAAAACCGTGAAACAAGAAATTAAAGAAAGCGTGTTTAGCAGCAAAGCATATG<br>GCACCCGTGAAACCAAAAATGTGAATATTGATGGTCGTCTGCAGCTGGATCTGCTGCAGTTTATCCAGCGTGAATATAAACTGCGTTCCTATACC<br>CTGAATGCAGTTAGCGCACATTTTCTGGGTGAACAGAAAGAAGATGTGCACTATAGCATTATTAGCGATCTGCAGAATGGTGATAGCGAAACCC<br>GTCGTCGTCTGGCAGTTTATTGTCTGAAAGATGCATATCTGCCGCTGCGCCTGATGGAAAAACTGATGGCACTGGTGAACTATACCGAAATGGC<br>ACGTGTTACCGGTGTTCCGTTTAGCTATCTGCTGGCACGTGGTCAGCAGATTAAAGTTGTTAGCCAGCTGTTTCGTAAATGCCTGGAAATTGATA<br>CCGTGATTCCGAATATGCAGAGCCAGGCAAGTGATGATCAGTATGAAGGTGCAACCGTTATTGAACCGATTCGCGGTTATTATGATGTTCCGATT<br>GCCACCCTGGATTTTAATAGCCTGTATCCGAGCATTATGATGGCCCATAATCTGTGTTATACCACCCTGTGTAATAAAGCAACCGTTGAACGCCT<br>GAACCTGAAAATCGATGAAGATTATGTTATTACCCCGAACGGCGATTATTTTGTTACCACCAAACGTCGTCGTGGTATTCTGCCGATTATTCTGGA<br>TGAACTGATTAGCGCACGTAAACGTGCAAAAAAAGATCTGCGTGATGAGAAAGATCCGTTTAAACGTGATGTTCTGAATGGTCGTCAGCTGGCA<br>CTGAAAATTAGCGCAAATAGCGTTTATGGTTTTACCGGTGCCACCGTTGGTAAACTGCCGTGTCTGGCAATTAGCAGCAGCGTTACCGCATATG<br>GTCGTACCATGATTCTGAAAACCAAAACCGCAGTGCAAGAGAAATACTGCATTAAAAACGGCTATAAACATGATGCCGTTGTGGTTTATGGTGAT<br>ACCGATAGCGTTATGGTTAAATTTGGCACCACCGATCTGAAAGAAGCAATGGATCTGGGCACCGAAGCAGCAAAATATGTTAGCACCCTGTTTA<br>AACATCCGATTAATCTGGAATTCGAAAAAGCCTATTTTCCGTACCTGCTGATCAACAAAAAACGTTATGCAGGTCTGTTTTGGACCAACCCGGATA<br>AATTTGATAAACTGGATCAGAAAGGTCTGGCAAGCGTTCGTCGTGATAGCTGTAGCCTGGTTAGCATTGTTATGAACAAAGTGCTGAAAAAAATC<br>CTGATCGAACGCAATGTTGATGGTGCACTGGCATTTGTTCGTGAAACCATTAATGATATTCTGCACAACCGTGTGGATATTAGCAAACTGATTATT<br>AGCAAAACCCTGGCACCGAATTATACCAATCCGCAGCCGCATGCAGTTCTGGCAGAACGTATGAAACGTCGTGAAGGTGTTGGTCCGAATGTTG<br>GTGATCGTGTTGATTATGTTATCATCGGTGGCAACGATAAACTGTATAATCGTGCAGAAGATCCGCTGTTTGTTCTGGAAAACAATATTCAGGTT<br>GATAGCCGCTATTATCTGACCAATCAGCTGCAGAATCCGATTATTAGCATTGTGGCACCGATTATTGGTGATAAACAGGCCAATGGTATGTTCGT<br>GGTGTAA |

**Table S16.** Primer Sequences used for Gibson Assembly of Pol3-CaM Fusions.

| Primer Name | DNA Sequence (5' - 3') |
|---|---|
| Forward Gibson_Fragment_Primer_F1-Pol3-CaM | AAATTTGATAAACTGGGAAGTGGTGGAGGTATGGCG |
| Reverse Gibson_Fragment_Primer_F1-Pol3-CaM | CAGACCTTTCTGATCACCTCCACCGGATCCTTTCG |
| Reverse Gibson_Vector_Primer_F1-Pol3-CaM | ACCTCCACCACTTCCCAGTTTATCAAATTTATCCGGGTTGG |
| Forward Gibson_Vector_Primer_F1-Pol3-CaM | GGATCCGGTGGAGGTGATCAGAAAGGTCTGGCAAGCG |
| Forward Gibson_Fragment_Primer_F2-Pol3-CaM | GGTATGTTCGTGGTGGGAAGTGGTGGAGGTATGGCG |
| Reverse Gibson_Fragment_Primer_F2-Pol3-CaM | CAGCCGGATCTCTTAACCTCCACCGGATCCTTTCG |
| Reverse Gibson_Vector_Primer_F2-Pol3-CaM | ACCTCCACCACTTCCCACCACGAACATACCATTGGCC |
| Forward Gibson_Vector_Primer_F2-Pol3-CaM | GGATCCGGTGGAGGTTAAGAGATCCGGCTGCTAACAAAGCCC |
| Forward Gibson_Fragment_Primer_F3-Pol3-CaM | GATAGCGAAACCCGTCGTCGTCTGGCAGTTGGAAGTGGTGGAGGTATGGCG |
| Reverse Gibson_Fragment_Primer_F3-Pol3-CaM | CAGCGGCAGATATGCATCTTTCAGACAATAACCTCCACCGGATCCTTTCG |
| Reverse Gibson_Vector_Primer_F3-Pol3-CaM | AACTGCCAGACGACGACGGGTTTC |
| Forward Gibson_Vector_Primer_F3-Pol3-CaM | TATTGTCTGAAAGATGCATATCTGCC |
| Forward Gibson_Fragment_Primer_F4-Pol3-CaM | CGTAAATGCCTGGAAGGAAGTGGTGGAGGTATGGCG |
| Reverse Gibson_Fragment_Primer_F4-Pol3-CaM | AATCACGGTATCAATACCTCCACCGGATCCTTTCG |
| Reverse Gibson_Vector_Primer_F4-Pol3-CaM | ACCTCCACCACTTCCTTCCAGGCATTTACGAAACAGCTG |
| Forward Gibson_Vector_Primer_F4-Pol3-CaM | GGATCCGGTGGAGGTATTGATACCGTGATTCCGAATATGCAG |
| Forward Gibson_Fragment_Primer_F5-Pol3-CaM | ATTATGAGCTTTGATGGAAGTGGTGGAGGTATGGCG |
| Reverse Gibson_Fragment_Primer_F5-Pol3-CaM | ACCTGCACATTCAATACCTCCACCGGATCCTTTCG |
| Reverse Gibson_Vector_Primer_F5-Pol3-CaM | ACCTCCACCACTTCCATCAAAGCTCATAATACGCAGCG |
| Forward Gibson_Vector_Primer_F5-Pol3-CaM | GGATCCGGTGGAGGTATTGAATGTGCAGGTCGTATTGGTGTTTTTCC |
| Forward Gibson_Fragment_Primer_F6-Pol3-CaM | ATTGCATATACCCTGGGAAGTGGTGGAGGTATGGCG |
| Reverse Gibson_Fragment_Primer_F6-Pol3-CaM | ATCAACCATCAGACGACCTCCACCGGATCCTTTCG |
| Reverse Gibson_Vector_Primer_F6-Pol3-CaM | ACCTCCACCACTTCCCAGGGTATATGCAATATTATCATAGGTGGTGG |
| Forward Gibson_Vector_Primer_F6-Pol3-CaM | GGATCCGGTGGAGGTCGTCTGATGGTTGATTGTGGTATTGT |
| Forward Gibson_Fragment_Primer_F7-Pol3-CaM | AGTGATGATCAGTATGGAAGTGGTGGAGGTATGGCG |
| Reverse Gibson_Fragment_Primer_F7-Pol3-CaM | AACGGTTGCACCTTCACCTCCACCGGATCCTTTCG |
| Reverse Gibson_Vector_Primer_F7-Pol3-CaM | ACCTCCACCACTTCCATACTGATCATCACTTGCCTGGC |
| Forward Gibson_Vector_Primer_F7-Pol3-CaM | GGATCCGGTGGAGGTGAAGGTGCAACCGTTATTGAACCGAT |
| Forward Gibson_Fragment_Primer_Part 1_F8-Pol3-CaM | TGTTACCGGTGTTCCGTTTAGCTATGGAAGTGGTGGAGGTATGG |
| Reverse Gibson_Fragment_Primer_Part 1_F8-Pol3-CaM | ATCTGCTGACCACGTGCCAGCAGACCTCCACCGGATCCTTTCG |
| Forward Gibson_Fragment_Primer_Part 2_F8-Pol3-CaM | GGCACTGGTGAACTATACCGAAATGGCACGTGTTACCGGTGTTCCGTTTAGC |
| Reverse Gibson_Fragment_Primer_Part 2_F8-Pol3-CaM | CATTTACGAAACAGCTGGCTAACAACTTTAATCTGCTGACCACGTGCCAG |
| Reverse Gibson_Vector_Primer_F8-Pol3-CaM | CGTGCCATTTCGGTATAGTTCAC |
| Forward Gibson_Vector_Primer_F8-Pol3-CaM | TAAAGTTGTTAGCCAGCTGTTTCG |
| Forward Gibson_Fragment_Primer_F9-Pol3-CaM | GAAATTAAAGAAAGCGGAAGTGGTGGAGGTATGGCG |
| Reverse Gibson_Fragment_Primer_F9-Pol3-CaM | TTTGCTGCTAAACACACCTCCACCGGATCCTTTCG |
| Reverse Gibson_Vector_Primer_F9-Pol3-CaM | ACCTCCACCACTTCCGCTTTCTTTAATTTCTTGTTTCACGGTTTT |
| Forward Gibson_Vector_Primer_F9-Pol3-CaM | GGATCCGGTGGAGGTGTGTTTAGCAGCAAAGCATATGGCAC |
| Forward Gibson_Fragment_Primer_F10-Pol3-CaM | ATTCCGAATATGCAGGGAAGTGGTGGAGGTATGGCG |
| Reverse Gibson_Fragment_Primer_F10-Pol3-CaM | ATCACTTGCCTGGCTACCTCCACCGGATCCTTTCG |
| Reverse Gibson_Vector_Primer_F10-Pol3-CaM | ACCTCCACCACTTCCCTGCATATTCGGAATCACGGTATCAA |
| Forward Gibson_Vector_Primer_F10-Pol3-CaM | GGATCCGGTGGAGGTAGCCAGGCAAGTGATGATCAG |
| Forward Gibson_Fragment_Primer_F11-Pol3-CaM | ATTATTGGTGATAAAGGAAGTGGTGGAGGTATGGCG |
| Reverse Gibson_Fragment_Primer_F11-Pol3-CaM | CATACCATTGGCCTGACCTCCACCGGATCCTTTCG |
| Reverse Gibson_Vector_Primer_F11-Pol3-CaM | ACCTCCACCACTTCCTTTATCACCAATAATCGGTGCCACAATG |
| Forward Gibson_Vector_Primer_F11-Pol3-CaM | GGATCCGGTGGAGGTCAGGCCAATGGTATGTTCGTGG |
| Forward Gibson_Fragment_Primer_Part 1_F12-Pol3-CaM | CAAAAATGTGAATATTGATGGTCGTCTGCAGCTGGATGGAAG |
| Reverse Gibson_Fragment_Primer_Part 1_F12-Pol3-CaM | AACGCAGTTTATATTCACGCTGGATAAACTGCAGCAGACCTCCAC |
| Forward Gibson_Fragment_Primer_Part 2_F12-Pol3-CaM | TAGCAGCAAAGCATATGGCACCCGTGAAACCAAAAATGTGAATATTGATGGTCGTC |
| Reverse Gibson_Fragment_Primer_Part 2_F12-Pol3-CaM | AATGTGCGCTAACTGCATTCAGGGTATAGGAACGCAGTTTATATTCACGCTGG |
| Reverse Gibson_Vector_Primer_F12-Pol3-CaM | GTTTCACGGGTGCCATATGC |
| Forward Gibson_Vector_Primer_F12-Pol3-CaM | CCTATACCCTGAATGCAGTTAGC |
| Forward Gibson_Fragment_Primer_F13-Pol3-CaM | GTTACCGGCTTTAAAGGAAGTGGTGGAGGTATGGCG |
| Reverse Gibson_Fragment_Primer_F13-Pol3-CaM | CACATACAGATAGTTACCTCCACCGGATCCTTTCG |
| Reverse Gibson_Vector_Primer_F13-Pol3-CaM | ACCTCCACCACTTCCTTTAAAGCCGGTAACATTACACAGAACG |
| Forward Gibson_Vector_Primer_F13-Pol3-CaM | GGATCCGGTGGAGGTAACTATCTGTATGTGCCTGCACC |
| Forward Gibson_Fragment_Primer_F14-Pol3-CaM | GCAATGGATCTGGGCGGAAGTGGTGGAGGTATGGCG |
| Reverse Gibson_Fragment_Primer_F14-Pol3-CaM | TTTTGCTGCTTCGGTACCTCCACCGGATCCTTTCG |
| Reverse Gibson_Vector_Primer_F14-Pol3-CaM | ACCTCCACCACTTCCGCCCAGATCCATTGCTTCTTTCAG |
| Forward Gibson_Vector_Primer_F14-Pol3-CaM | GGATCCGGTGGAGGTACCGAAGCAGCAAAATATGTTAGCAC |
| Forward Gibson_Fragment_Primer_F15-Pol3-CaM | TTTGTTACCACCAAACGTCGTCGTGGTATTGGAAGTGGTGGAGGTATGGCG |
| Reverse Gibson_Fragment_Primer_F15-Pol3-CaM | GCTAATCAGTTCATCCAGAATAATCGGCAGACCTCCACCGGATCCTTTCG |
| Reverse Gibson_Vector_Primer_F15-Pol3-CaM | AATACCACGACGACGTTTGG |
| Forward Gibson_Vector_Primer_F15-Pol3-CaM | CTGCCGATTATTCTGGATGAACTG |
| Forward Gibson_Fragment_Primer_F16-Pol3-CaM | AAACTGATGGCACTGGGAAGTGGTGGAGGTATGGCG |
| Reverse Gibson_Fragment_Primer_F16-Pol3-CaM | TTCGGTATAGTTCACACCTCCACCGGATCCTTTCG |
| Reverse Gibson_Vector_Primer_F16-Pol3-CaM | ACCTCCACCACTTCCCAGTGCCATCAGTTTTTCCATCAG |
| Forward Gibson_Vector_Primer_F16-Pol3-CaM | GGATCCGGTGGAGGTGTGAACTATACCGAAATGGCACGTG |
| Forward Gibson_Fragment_Primer_F17-Pol3-CaM | GTTCCGATTGCCACCGGAAGTGGTGGAGGTATGGCG |
| Reverse Gibson_Fragment_Primer_F17-Pol3-CaM | GCTATTAAAATCCAGACCTCCACCGGATCCTTTCG |
| Reverse Gibson_Vector_Primer_F17-Pol3-CaM | ACCTCCACCACTTCCGGTGGCAATCGGAACATCATAATAACC |
| Forward Gibson_Vector_Primer_F17-Pol3-CaM | GGATCCGGTGGAGGTCTGGATTTTAATAGCCTGTATCCGAGCA |
| Forward Gibson_Fragment_Primer_F18-Pol3-CaM | CGTCAGCTGGCACTGGGAAGTGGTGGAGGTATGGCG |
| Reverse Gibson_Fragment_Primer_F18-Pol3-CaM | ATTTGCGCTAATTTTACCTCCACCGGATCCTTTCG |
| Reverse Gibson_Vector_Primer_F18-Pol3-CaM | ACCTCCACCACTTCCCAGTGCCAGCTGACGAC |
| Forward Gibson_Vector_Primer_F18-Pol3-CaM | GGATCCGGTGGAGGTAAAATTAGCGCAAATAGCGTTTATGGTTT |
| Forward Gibson_Fragment_Primer_F19-Pol3-CaM | GATCAGTATGAAGGTGGAAGTGGTGGAGGTATGGCG |
| Reverse Gibson_Fragment_Primer_F19-Pol3-CaM | TTCAATAACGGTTGCACCTCCACCGGATCCTTTCG |
| Reverse Gibson_Vector_Primer_F19-Pol3-CaM | ACCTCCACCACTTCCACCTTCATACTGATCATCACTTGCCT |
| Forward Gibson_Vector_Primer_F19-Pol3-CaM | GGATCCGGTGGAGGTGCAACCGTTATTGAACCG |
